# Coordination of Tissue Cell Polarity by Auxin Transport and Signaling

**DOI:** 10.1101/680090

**Authors:** Carla Verna, Sree Janani Ravichandran, Megan G. Sawchuk, Nguyen Manh Linh, Enrico Scarpella

## Abstract

Coordination of polarity between cells in tissues is key to multicellular organism development. In animals, coordination of this tissue cell polarity often requires direct cell-cell interactions and cell movements, which are precluded in plants by a wall that separates cells and holds them in place; yet plants coordinate the polarity of hundreds of cells during the formation of the veins in their leaves. Overwhelming experimental evidence suggests that the plant signaling molecule auxin coordinates tissue cell polarity to induce vein formation, but how auxin does so is unclear. The prevailing hypothesis proposes that GNOM, a regulator of vesicle formation during protein trafficking, positions auxin transporters of the PIN-FORMED family to the correct side of the plasma membrane. The resulting cell-to-cell, polar transport of auxin would coordinate tissue cell polarity and would induce vein formation. Here we tested this hypothesis by means of a combination of cellular imaging, molecular genetic analysis, and chemical induction and inhibition. Contrary to predictions of the hypothesis, we find that auxin-induced vein formation occurs in the absence of PIN-FORMED proteins or any known intercellular auxin transporter, that the residual auxin-transport-independent vein-patterning activity relies on auxin signaling, and that a *GNOM*-dependent signal that coordinates tissue cell polarity to induce vein formation acts upstream of both auxin transport and signaling. Our results reveal synergism between auxin transport and signaling, and their unsuspected control by *GNOM*, in the coordination of tissue cell polarity during vein patterning, one of the most spectacular and informative expressions of tissue cell polarization in plants.

## Introduction

How the polarity of the cells in a tissue is coordinated is a central question in biology. In animals, the coordination of this tissue cell polarity requires direct cell-cell communication and often cell movements (Goodrich and Strutt, 2011), both of which are precluded in plants by a wall that holds cells in place; therefore, tissue cell polarity is coordinated by a different mechanism in plants.

Plant veins are an expression of coordinated tissue cell polarity (Sachs, 1991b; Sachs, 2000; Boutte et al., 2007; Nakamura et al., 2012). This is reflected in the relation between the parts of the vein, and between the veins and the parts of the plant: vascular elements are elongated along the length of the vein and are connected to one another at their ends (Esau, 1942), and veins primarily connect shoot organs with roots (Dengler, 2006); therefore, veins and their elements are unequal at their ends — one end connects to shoot tissues, the other to root tissues — and are thus polar (Sachs, 1975). Not all the veins in closed networks such as those of Arabidopsis leaves have unambiguous shoot-to-root polarity, but the vein networks themselves are polar (Sachs, 1975).

Just as veins are an expression of coordinated tissue cell polarity, their formation is an expression of coordination of tissue cell polarity; this is most evident in developing leaves. Consider, for example, the formation of the midvein at the center of the cylindrical leaf primordium. Initially, the plasma-membrane (PM)-localized PIN-FORMED1 (PIN1) protein of Arabidopsis (Galweiler et al., 1998), which catalyzes cellular efflux of the plant signal auxin (Petrasek et al., 2006), is expressed in all the inner cells of the leaf primordium (Benkova et al., 2003; Reinhardt et al., 2003; Heisler et al., 2005; Scarpella et al., 2006; Wenzel et al., 2007; Bayer et al., 2009; Verna et al., 2015); over time, however, PIN1 expression becomes gradually restricted to the file of cells that will form the midvein. PIN1 localization at the PM of the inner cells is initially isotropic, or nearly so, but as PIN1 expression becomes restricted to the site of midvein formation, PIN1 localization becomes polarized: in the cells surrounding the developing midvein, PIN1 localization gradually changes from isotropic to medial, i.e. toward the developing midvein, to mediobasal; and in the cells of the developing midvein, PIN1 becomes uniformly localized toward the base of the leaf primordium, where the midvein will connect to the pre-existing vasculature. Both the restriction of PIN1 expression and the polarization of PIN1 localization initiate and proceed away from pre-existing vasculature and are thus polar.

The correlation between (1) coordination of tissue cell polarity, as expressed by the coordination of PIN1 polar localization between cells, (2) polar auxin transport, as expressed by the auxin-transport-polarity-defining localization of PIN1 (Wisniewska et al., 2006), and (3) the polar formation of veins, themselves polar, does not seem to be coincidental. Auxin application to developing leaves induces the formation of broad expression domains of isotropically localized PIN1; such domains become restricted to the sites of auxin-induced vein formation, and PIN1 localization becomes polarized toward the pre-existing vasculature (Scarpella et al., 2006). Both the restriction of PIN1 expression domains and the polarization of PIN1 localization are delayed by chemical inhibition of auxin transport (Scarpella et al., 2006; Wenzel et al., 2007), which induces vein pattern defects similar to, though stronger than, those of *pin1* mutants (Mattsson et al., 1999; Sieburth, 1999; Sawchuk et al., 2013).

Therefore, available evidence suggests that auxin coordinates tissue cell polarizaty to induce polar-vein-formation, and it seems that such coordinative and inductive property of auxin strictly depends on the function of *PIN1* and possibly other *PIN* genes. How auxin precisely coordinates tissue cell polarity to induce polar-vein-formation is unclear, but the current hypothesis is that the GNOM (GN) guanine-nucleotide exchange factor for ADP-rybosilation-factor GTPases, which regulates vesicle formation in membrane trafficking, controls the cellular localization of PIN1 and other PIN proteins; the resulting cell-to-cell, polar transport of auxin would coordinate tissue cell polarity and control polar developmental processes such as vein formation (reviewed in, e.g., (Berleth et al., 2000; Richter et al., 2010; Nakamura et al., 2012; Linh et al., 2018)).

Here we tested this hypothesis by a combination of cellular imaging, molecular genetic analysis and chemical induction and inhibition. Contrary to predictions of the hypothesis, we found that auxin-induced polar-vein-formation occurs in the absence of PIN proteins or any known intercellular auxin transporter, that the residual auxin-transport-independent vein-patterning activity relies on auxin signaling, and that a *GN*-dependent tissue-cell-polarizing signal acts upstream of both auxin transport and signaling.

## Results

### Contribution of the *GNOM* Gene to Coordination of Tissue Cell Polarity During Arabidopsis Vein Formation

The current hypothesis of how auxin coordinates tissue cell polarity to induce polar-vein-formation proposes that the GNOM (GN) guanine-nucleotide exchange factor for ADP-ribosylation-factor GTPases, which regulates vesicle formation in membrane trafficking, controls the cellular localization of PIN1; the resulting cell-to-cell, polar transport of auxin would coordinate cell polarity between cells, and control polar developmental processes such as vein formation (reviewed in, e.g., (Berleth et al., 2000; Richter et al., 2010; Nakamura et al., 2012; Linh et al., 2018)). As such, the hypothesis predicts that the restriction of PIN1 expression domains and coordination of PIN1 polar localization that normally occur during vein formation (Benkova et al., 2003; Reinhardt et al., 2003; Heisler et al., 2005; Scarpella et al., 2006; Wenzel et al., 2007; Bayer et al., 2009; Sawchuk et al., 2013; Marcos and Berleth, 2014; Verna et al., 2015) would occur abnormally, or fail to occur altogether, during *gn*-mutant leaf development.

We first tested this prediction by imaging expression domains of PIN1::PIN1:YFP (PIN1:YFP fusion protein expressed by the PIN1 promoter (Xu et al., 2006)) in WT and in the new strong allele *gn*-13 (Table S1) during first-leaf development.

Consistent with previous reports (Benkova et al., 2003; Reinhardt et al., 2003; Heisler et al., 2005; Scarpella et al., 2006; Wenzel et al., 2007; Bayer et al., 2009; Sawchuk et al., 2013; Marcos and Berleth, 2014; Verna et al., 2015), in WT leaves PIN1::PIN1:YFP was expressed in all the cells at early stages of tissue development; over time, epidermal expression became restricted to the basal-most cells and inner tissue expression became restricted to files of vascular cells (Fig. 1A-J).

**Figure 1.**
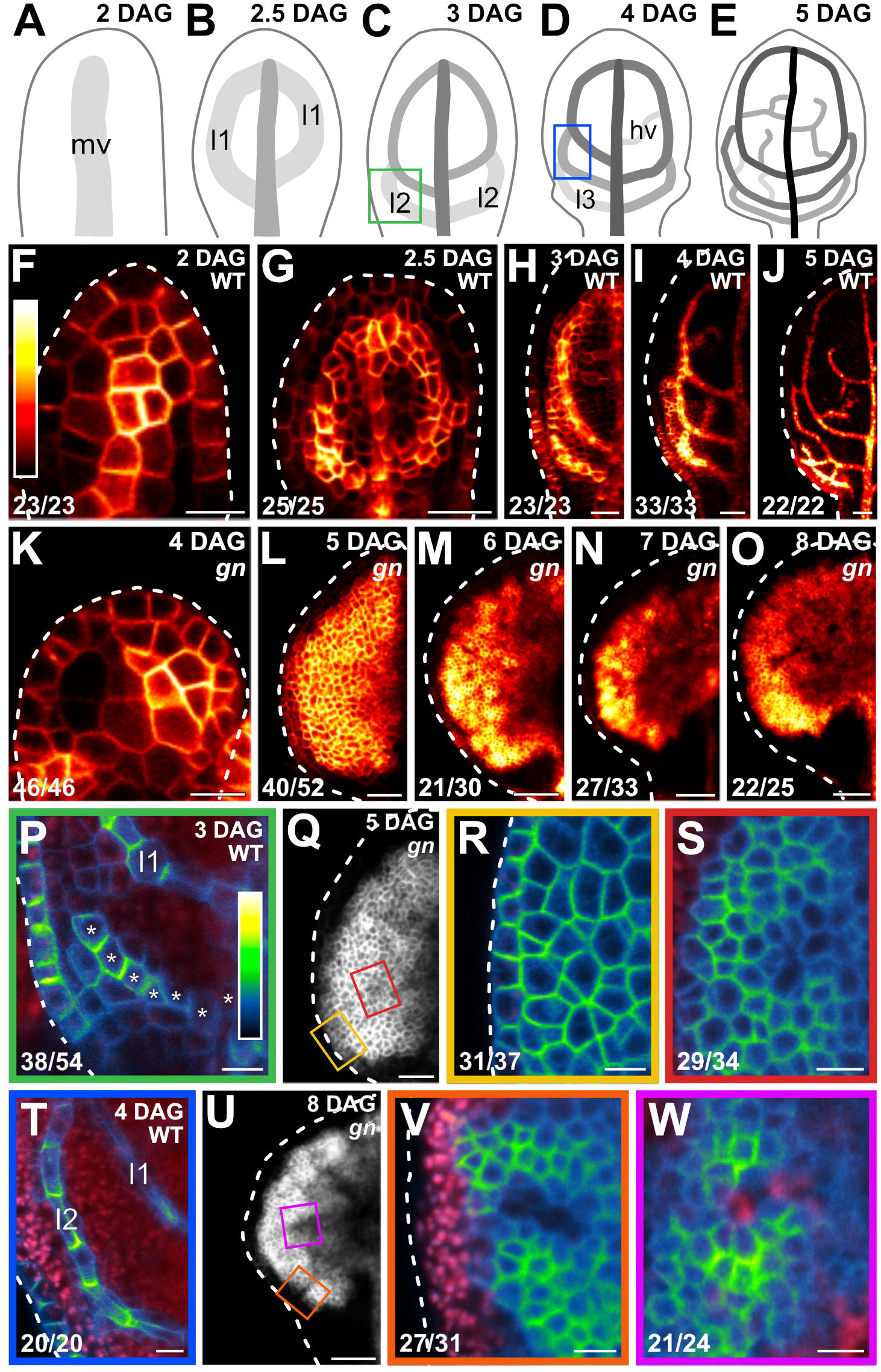
Contribution of the *GNOM* Gene to Coordination of Tissue Cell Polarity During Arabidopsis Vein Formation. (A–Q,T,U) Top right: leaf age in days after germination (DAG). (A–E) Veins form sequentially during Arabidopsis leaf development: the formation of the midvein (mv) is followed by the formation of the first loops of veins (“first loops”; 11), which in turn is followed by the formation of second loops (12) and minor veins (hv) (Mattsson et al., 1999; Sieburth, 1999; Kang and Dengler, 2004; Scarpella et al., 2004). Loops and minor veins differentiate in a tip-to-base sequence during leaf development. Increasingly darker grays depict progressively later stages of vein development. Boxes in C and D illustrate positions of closeups in P and T. 13: third loop. (F–W) Confocal laser scanning microscopy. First leaves. For simplicity, only half-leaves are shown in H–J and L–O. Dashed white line in F–R, T, U and V delineates leaf outline. (F–Q,T,U) Top right: genotype. (F–P,R–T,V,W) Bottom left: reproducibility index. (F–O) PIN1::PIN1:YFP expression; look-up table (ramp in F) visualizes expression levels. (P,R–T,V,W) PIN1::PIN1:GFP expression; look-up table (ramp in P) visualizes expression levels. Red: autofluorescence. Stars in P label cells of the developing second loop. (Q,U) PIN1::PIN1:YFP expression. Boxes in Q and in U illustrate positions of closeups in R and S and in V and W, respectively. Bars: (F,P,R–T,V,W) 10 μm; (G,I,L,Q) 30 μm; (H,K) 20 μm; (J,M–O,U) 60 μm.

In *gn* leaves too, PIN1::PIN1:YFP was expressed in all the cells at early stages of tissue development and over time epidermal expression became restricted to the basal-most cells; however, inner tissue expression failed to become restricted to files of vascular cells and instead remained nearly ubiquitous even at very late stages of leaf development (Fig. 1K-O).

We next tested the prediction by imaging cellular localization of expression of PIN1::PIN1:GFP (Benkova et al.,2003) in WT and during first-leaf development. Hereafter, we use “basal” to describe localization of PIN::PIN1:GFP expression oriented toward pre-existing veins, irrespective of how these veins are positioned within a leaf.

Consistent with previous reports (Benkova et al., 2003; Reinhardt et al., 2003; Heisler et al., 2005; Scarpella et al., 2006; Wenzel et al., 2007; Bayer et al., 2009; Sawchuk et al., 2013; Marcos and Berleth, 2014; Verna et al., 2015), in the cells of the second pair of vein loops (“second loop” hereafter) at early stages of its development in WT leaves, PIN1::PIN1:GFP expression was mainly localized to the basal side of the plasma membrane (PM), toward the midvein; in the inner cells flanking the developing loop, PIN1::PIN1:GFP expression was mainly localized to the side of the PM facing the developing loop; and in the inner cells further away from the developing loop, PIN1::PIN1:GFP expression was localized isotropically, or nearly so, at the PM (Fig. 1C,P). At later stages of second-loop development, by which time PIN1::PIN1:GFP expression had become restricted to the sole, elongated cells of the developing loop, PIN1::PIN1:GFP expression was localized to the basal side of the PM, toward the midvein (Fig. 1D,T).

At early stages of development of the tissue that in *gn* leaves corresponds to that from which the second loop forms in WT leaves, PIN1::PIN1:GFP was expressed uniformly in the outermost inner tissue and expression was localized isotropically, or nearly so, at the PM (Fig. 1Q,R). PIN1::PIN1:GFP was expressed more heterogeneously in the innermost inner tissue, but expression remained localized isotropically, or nearly so, at the PM, except in cells near the edge of higher-expression domains (Fig. 1Q,S); in those cells, localization of PIN1::PIN1:GFP expression at the PM was weakly polar, but such weak cell polarities pointed in seemingly random directions (Fig. 1Q,S).

At late stages of *gn* leaf development, heterogeneity of PIN1::PIN1:GFP expression had spread to the outermost inner tissue, but expression remained localized isotropically, or nearly so, at the PM, except in cells near the edge of higher-expression domains (Fig. 1U,V); in those cells, localization of PIN1::PIN1:GFP expression at the PM was weakly polar, but such weak cell polarities pointed in seemingly random directions (Fig. 1U,V). Heterogeneity of PIN1::PIN1:GFP expression in the innermost inner tissue had become more pronounced at late stages of *gn* leaf development, and the weakly polar localization of PIN1:: PIN1:GFP expression at the PM had spread to the center of the higher-expression domains (Fig. 1U,W); nevertheless, such weak cell polarities still pointed in seemingly random directions (Fig. 1U,W). Finally, none of the cells had acquired the elongated shape characteristic of vascular cells in WT (Fig. 1U-W).

In conclusion, consistent with previous observations (Steinmann et al., 1999; Kleine-Vehn et al., 2008), both restriction of PIN1 expression domains and coordination of PIN1 polar localization occur only to a very limited extent or fail to occur altogether during *gn* leaf development, which is consistent with the current hypothesis of how auxin coordinates tissue cell polarity to induce polar-vein-formation.

### Contribution of *GN to* Vein Patterning

We asked whether the very limited or altogether absent restriction of PIN1 expression domains and coordination of PIN1 polar localization occurring during *gn* leaf development (Figure 1) were associated with vein pattern defects in mature *gn* leaves.

WT Arabidopsis grown under normal conditions forms separate leaves whose vein networks are defined by at least four reproducible features (Telfer and Poethig, 1994; Nelson and Dengler, 1997; Kinsman and Pyke, 1998; Candela et al., 1999; Mattsson et al., 1999; Sieburth, 1999; Steynen and Schultz, 2003; Sawchuk et al., 2013; Verna et al., 2015) (Fig. 2A,B): (1) a narrow I-shaped midvein that runs the length of the leaf; (2) lateral veins that branch from the midvein and join distal veins to form closed loops; (3) minor veins that branch from midvein and loops, and either end freely or join other veins; (4) minor veins and loops that curve near the leaf margin, lending a scalloped outline to the vein network.

**Figure 2.**
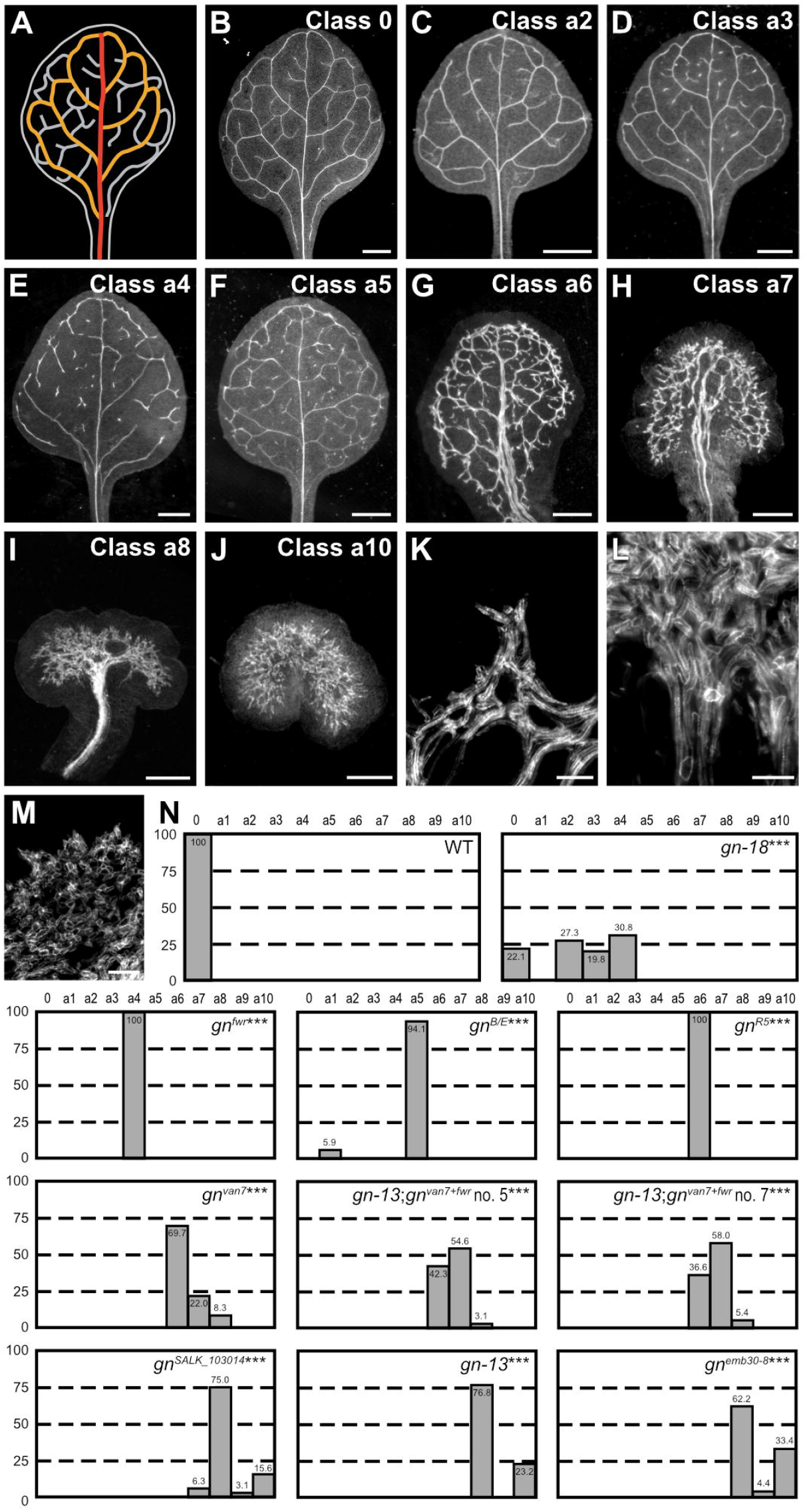
Contribution of *GN* to Vein Patterning. (A,B) Vein pattern of WT mature first leaf. In A: red, midvein; orange, loops; gray, minor veins. (B–J) Dark-field illumination of mature first leaves illustrating phenotype classes (top right): class 0, narrow I-shaped midvein and scalloped vein-network outline (B); class a1, dense vein network and apically thickened vein-network outline (not shown); class a2, open vein-network outline (C); class a3, fragmented vein network (D); class a4, open vein-network outline and fragmented vein network (E); class a5, open vein-network outline, fragmented vein network and apically thickened vein-network outline (F); class a6, wide midvein, dense network of thick veins and jagged vein-network outline (G); class a7, dense network of thick veins that fail to join the midvein in the bottom half of the leaf and pronouncedly jagged vein-network outline (H); class a8, wide midvein and shapeless vascular cluster (I); class a9, fused leaves with wide midvein and shapeless vascular cluster (not shown); class a10, shapeless vascular cluster (J). (K–M) Details of vascular clusters illustrating vascular elements uniformly oriented perpendicular to the leaf margin (K) (class a6), vascular elements oriented seemingly randomly at the distal side of the cluster and parallel to the leaf axis at the proximal side of the cluster (L) (classes a8 and a9), and seemingly random orientation of vascular elements (M) (classed a8–a10). (N) Percentages of leaves in phenotype classes. Difference between *gn-18* and WT, between *gn^fwr^* and WT, between *gn^B/E^* and WT, between *gn^R5^* and WT, between *gn^van7^* and WT, between *gn^van7+fwr^*;*gn-13* and WT, between *gn^SALK_103014^* and WT, between *gn-13* and WT, and between *emb30-8* and WT was significant at P<0.001 (***) by Kruskal-Wallis and Mann-Whitney test with Bonferroni correction. Sample population sizes: WT, 58; *gn-18*, 172; *gn^fwr^*, 43; *gn^B/E^*, 80; *gn^R5^*, 93; *gn^van7^*, 109; *gn^van7+fwr^;gn-13* no. 5, 97; *gn^van7+fwr^;gn-13* no. 7, 93; *gn^SALK_103014^*, 32; *gn-13*, 56; *gn^emb30-8^*, 45. Bars: (B–F) 1 mm; (G) 0.75 mm; (H,I) 0.5 mm; (J) 0.25 mm; (K–M) 50 μm.

In ~25% of the leaves of the new weak allele *gn*-18 (Table S1) (Figure S1) closed loops were often replaced by open loops, i.e. loops that contact the midvein or other loops at only one of their two ends (Fig. 2C,N). Moreover, in ~50% of *gn-18* leaves veins were often replaced by “vein fragments”, i.e. stretches of vascular elements that fail to contact other stretches of vascular elements at either one of their two ends (Fig. 2D,E,N). Loops were open and veins were fragmented also in the leaves of both *gn^fwr^* (Okumura et al., 2013) and *gn^B/E^* (Geldner et al., 2004) (Fig. 2N). In addition, the vein network of *gn^B/E^* leaves was denser and its outline was thicker near the leaf tip (Fig. 2F,N).

The vein network was denser also in all the leaves of *gn^R5^* (Geldner et al., 2004), in nearly 70% of those of *gn^van7^* (Koizumi et al., 2000) and in ~40% of those of *gn^van7+fwr^*;*gn-13* — in which we had combined the *van7* and *fwr* mutations (Table S1) (Fig. 2G,N). However, in the leaves of these backgrounds — unlike in those of *gn^B/E^* — all the veins were thicker; lateral veins failed to join the midvein but ran parallel to it to form a “wide midvein”; and the vein network outline was jagged because of narrow clusters of vascular elements that were oriented perpendicular to the leaf margin and that were laterally connected by veins (Fig. 2G,K,N). These features were enhanced in ~20% of the leaves of *gn^van7^*, in ~55% of those of *gn^van7+fwr^;gn-13* and in ~5% of those of *gn^SALK_103014^* (Okumura et al., 2013): the vein network was denser, veins failed to join the midvein in the bottom half of the leaf, and the vein network outline was pronouncedly jagged (Fig. 2H,N).

Consistent with previous observations (Shevell et al., 2000), in the few remaining leaves of *gn^van7^* and *gn^van7+fwr^*; *gn-13*, and in most of those of *gn^SALK_103014^, gn-13* and *gn^emb30-8^* (Franzmann et al., 1989; Moriwaki et al., 2014), a central, shapeless vascular cluster was connected with the basal part of the leaf by a wide midvein, and vascular elements were oriented seemingly randomly at the distal side of the cluster and progressively more parallel to the leaf axis toward the proximal side of the cluster (Fig. 2I,L–N).

Finally, in the remaining leaves of *gn^SALK_103014^, gn-13* and *gn^emb30-8^*, vascular differentiation was limited to a central, shapeless cluster of seemingly randomly oriented vascular elements (Fig. 2J,M,N).

We conclude that defects in coordination of PIN1 polar localization and possible derived defects in polar auxin transport during *gn* leaf development are associated with vein pattern defects in mature *gn* leaves.

### Contribution of Plasma-Membrane-Localized PIN Proteins to Vein Patterning

Were the vein pattern defects of *gn* the sole result of loss of PIN1-mediated polar auxin-transport induced by defects in coordination of PIN1 polar localization, the vein pattern defects of *gn* would be phenocopied by simultaneous mutation in all the *PIN* genes with function in *PIN1*-dependent vein patterning; we asked whether that were so.

In Arabidopsis, the PIN family of auxin transporters is composed of eight members (Paponov et al., 2005; Krecek et al., 2009; Viaene et al., 2012): PIN5, PIN6 and PIN8, which are primarily localized to the endoplasmic reticulum (ER) (Mravec et al., 2009; Bosco et al., 2012; Ding et al., 2012; Sawchuk et al., 2013), and PIN1, PIN2, PIN3, PIN4 and PIN7 are primarily localized to the plasma membrane (PM) and catalyze cellular auxin efflux (Chen et al., 1998; Galweiler et al., 1998; Luschnig et al., 1998; Muller et al., 1998; Friml et al.,2002a; Friml et al., 2002b; Friml et al., 2003; Petrasek et al., 2006; Yang and Murphy, 2009; Zourelidou et al., 2014). Sequence analysis divides the PM-localized subfamily of PIN (PM-PIN) proteins into three groups: the PIN1 group, the PIN2 group and the PIN3 group, which also contains PIN4 and PIN7 (Krecek et al., 2009; Viaene et al., 2012).

Mutants of *PIN1* are the only *pin* single mutants with vein pattern defects, and the vein pattern defects of double mutants between *pin1* and mutants of *PIN2, PIN3, PIN4* or *PIN7* are no different from those of *pin1* single mutants (Sawchuk et al., 2013), suggesting that either *PIN2, PIN3, PIN4* and *PIN7* have no function in *PIN1*-dependent vein patterning or their function in this process is redundant. To discriminate between these possibilities, we first assessed the collective contribution to *PIN1*-dependent vein patterning of the *PM-PIN* genes of the *PIN3* group (*PIN3, PIN4* and *PIN7*), whose translational fusions to GFP (Zadnikova et al., 2010; Bennett et al., 2016; Belteton et al., 2018) (Table S1) are all expressed — as are translational fusions of PIN1 to GFP (Benkova et al., 2003; Heisler et al., 2005; Scarpella et al., 2006; Wenzel et al., 2007; Bayer et al., 2009; Marcos and Berleth, 2014) — in both epidermal and inner cells of the developing leaf (Fig. 3A,C-E).

**Figure 3.**
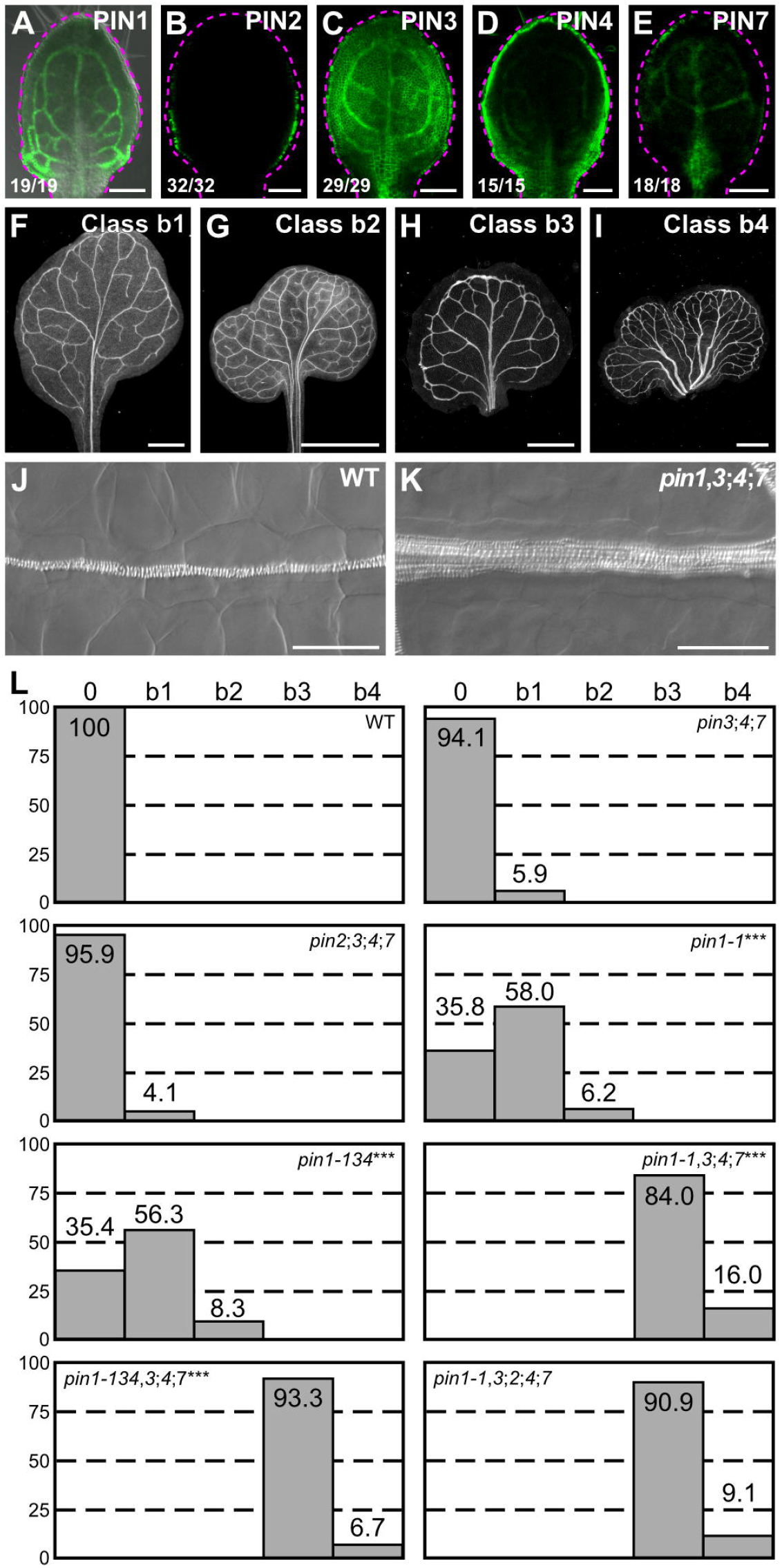
Contribution of Plasma-Membrane-Localized PIN Proteins to Vein Patterning. (A–K) Top right: expression-reported gene, phenotype class or genotype. (B–E) Bottom left: reproducibility index. (A–E) Confocal laser scanning microscopy with (A) or without (B–E) transmitted light; 4-day-old first leaves. Dashed magenta line delineates leaf outline. (A) PIN1::PIN1:GFP expression. (B) PIN2::PIN2:GFP expression. (C) PIN3::PIN3:GFP expression. (D) PIN4::PIN4:GFP expression. (E) PIN7::PIN7:GFP expression. (F–I) Dark-field illumination images of mature first leaves illustrating phenotype classes: class b1, Y-shaped midvein and scalloped vein-network outline (F); class b2, fused leaves with scalloped vein-network outline (G); class b3, thick veins and scalloped vein-network outline (H); class b4, fused leaves with thick veins and scalloped vein-network outline (I). (J,K) Differential interference images of details of WT (J) or *pin1-1,3;4;7* (K) illustrating normal (classes 0, b1 and b2) or thick (classes b3 and b4) veins, respectively. (L) Percentages of leaves in phenotype classes. Difference between *pin1-1* and WT, between *pin1-134* and WT, between *pin1-1,3;4;7* and *pin1-1*, and between *pin1-134,3;4;7* and *pin1-134*-wassignificantat P<0.001 (***) by Kruskal-Wallis and Mann-Whitney test with Bonferroni correction. Sample population sizes: WT, 58; *pin2;5;4;7*, 49; *pin3,4;7*, 102; *pin1-1*, 81; *pin1-134*, 48; *pin1-1,3;4;7,7*ĩ>; *pin1-134,3;4;7*, 45, *pin1-1,3;2;4;7*, 99. Bars: (A–E) 0.1 mm; (F–H) 1 mm; (I) 5 mm; (J,K) 50 μm.

Consistent with previous reports (Sawchuk et al., 2013; Verna et al., 2015), the vein patterns of most of the *pin1* leaves were abnormal (Fig. 3F,G,L). *pin3;pin4;pin7* (*pin3;4;7* hereafter) embryos were viable and developed into seedlings (Table S2) whose vein patterns were no different from those of WT (Fig. 3L). *pin1,3;4;7* embryes were viable (Table S3) and developed into seedlings (Table S4) that were smaller than *pin1* seedlings (Fig. S2A,B). The cotyledon pattern defects of *pin1,3;40;7* were more severe than those of *pin1* (Fig. S3A–H), and the vein pattern defects of *pin1,3;4;7* were more severe than those of *pin1*: no *pin1,3;4;7* leaf had a WT vein pattern; *pin1,3;4;7* veins were thicker; and ~15% of *pin1,3;4;7* leaves were fused (Fig. 3H–L). However, as in WT, in *pin1,3;4;7* vascular elements were elongated and aligned along the length of the vein (Fig. 3J,K).

Next, we asked whether mutation of *PIN2* — whose translational fusion to GFP (Xu and Scheres, 2005) is only expressed in epidermal cells in the developing leaf (Fig. 3B) — changed the spectrum of vein pattern defects of *pin1,3;4;7*.

*pin2;3;4;7* embryos were viable and developed into seedlings (Table S2) whose vein patterns were no different from those of WT (Fig. 3L). *pin1,3;2;4;7* embryos were viable (Table S3) and developed into seedlings (Table S4) whose vein pattern defects were no different from those of *pin1,3;4;7* (Fig. 3L). The cotyledon pattern defects of *pin1,3;2;4;7* were more severe than those of *pin1,3;4;7* (Fig. S3A–H), but the size of *pin1,3;2;4;7* seedlings was similar to that of *pin1,3;4;7* seedlings (Fig. S2A–C).

In conclusion, the *PIN3* group of *PM-PIN* genes (*PIN3, PIN4* and *PIN7*) provides no nonredundant function in vein patterning, but it contributes to *PIN1*-dependent vein patterning; *PIN1* and the *PIN3* group of *PM-PIN* genes redundantly restrict vascular differentiation to narrow zones; and *PIN2* seems to have no function in any of these processes. Most important, loss of *PM-PIN* function fails to phenocopy the vein pattern defects of *gn*.

### Contribution of *PIN* Genes to Vein Patterning

Expression and genetic analyses suggest that PIN1, PIN3, PIN4 and PIN7 redundantly define a single auxin-transport pathway with vein patterning functions whose loss fails to phenocopy the vein pattern defects of *gn* (Figure 2; Figure 3). The ER-localized PIN (ER-PIN) proteins PIN6 and PIN8, but not the ER-PIN protein PIN5, define a distinct auxin-transport pathway with vein patterning functions that overlap with those of *PIN1* (Sawchuk et al., 2013; Verna et al., 2015). We asked what the collective contribution of these two auxin-transport pathways were to vein patterning and whether simultaneous mutation in all the *PIN* genes with vein patterning function phenocopied the vein pattern defects of *gn*.

As previously reported (Sawchuk et al., 2013), the vein pattern of *pin6;8* was no different from that of WT (Fig. 4C). *pin1,3,6;4;7;8* embryos were viable (Table S3) and developed into seedlings (Table S4) whose vein patterns differed from those of *pin1,3,4;7* in four respects: (1) the vein network comprised more lateral veins; (2) lateral veins failed to join the midvein but ran parallel to it to form a wide midvein; (3) lateral veins ended in a marginal vein that closely paralleled the leaf margin, lending a smooth outline to the vein network; (4) veins were thicker (Figure 3; Fig. 4A–C). Simultaneous mutation of *PIN6* and *PIN8* in the *pin1,3;4;7* background shifted the distribution of *pin1,3;4;7* cotyledon pattern phenotypes toward stronger classes (Fig. S3A–H), but the size of *pin1,3,6;4;7;8* seedlings was similar to that of *pin1,3;4;7* seedlings (Fig. S2A,B,D).

**Figure 4.**
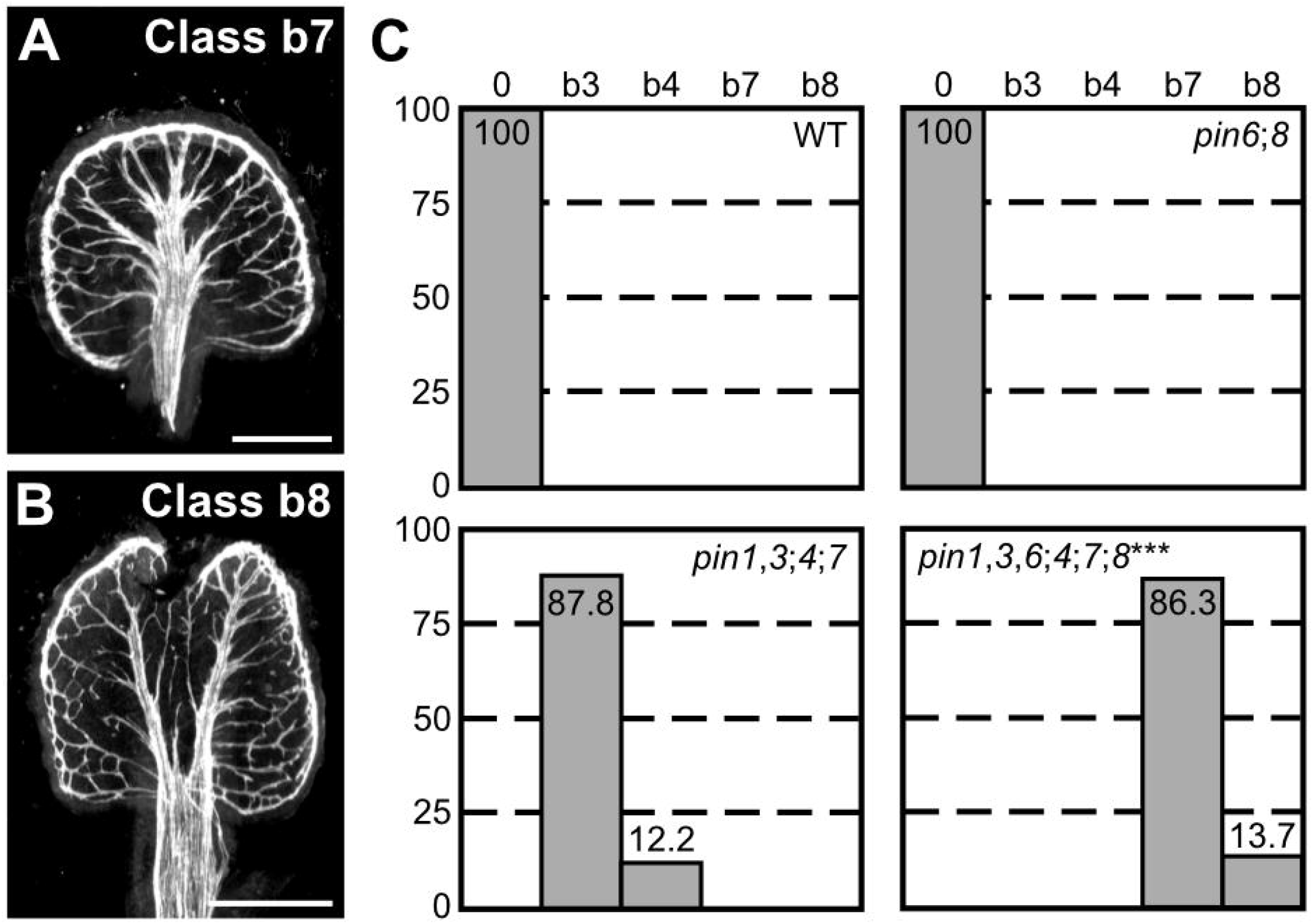
Contribution of *PIN* Genes to Vein Patterning. (A,B) Dark-field illumination of mature first leaves illustrating phenotype classes (top right): class b7, wide midvein, more lateral-veins and conspicuous marginal vein (A); class b8, fused leaves with wide midvein, more lateral-veins and conspicuous marginal vein (B). (C) Percentages of leaves in phenotype classes (Classes 0, b3 and b4 defined in Figures 2 and 3). Difference between *pin1-1,3;4;7;8* and *pin1;4;7* was significant at P<0.001 (***) by Kruskal-Wallis and Mann-Whitney test with Bonferroni correction. Sample population sizes: WT, 51; *pin6;8*, 47; *pin1-1,3;4;7*, 49; *pin1-1,3,6;4;7;8*, 73. Bars: (A,B) 0.5 mm.

Because *pin6;8* synthetically enhanced vein pattern defects of *pin1,3;4;7*, we conclude that the auxin-transport pathway mediated by PIN1, PIN3, PIN4 and PIN7 and that mediated by PIN6 and PIN8 provide overlapping functions in vein patterning. Nevertheless, loss of *PIN*-dependent vein patterning function fails to phenocopy the vein pattern defects of *gn*.

### Genetic Versus Chemical Inhibition of Auxin Transport During Vein Patterning

Loss of *PIN*-dependent vein patterning function fails to phenocopy the vein pattern defects of *gn* (Figure 2; Figure 4), suggesting that these latter are not the sole result of loss of *PIN*-dependent polar auxin-transport induced by defects in coordination of PIN polar localization. However, it is possible that the vein pattern defects of *gn* result from additional or exclusive defects in *PIN*-independent polar auxin-transport pathways; we asked whether that were so.

Cellular auxin efflux is inhibited by a class of structurally related compounds referred to as phytotropins, exemplified by N-1-naphthylphthalamic acid (NPA) (Cande and Ray, 1976; Katekar and Geissler, 1980; Sussman and Goldsmith, 1981). Because PM-PIN proteins catalyze cellular auxin efflux (Chen et al., 1998; Petrasek et al., 2006; Yang and Murphy, 2009; Zourelidou et al., 2014), we first asked whether defects resulting from simultaneous mutation of all the *PM-PIN* genes with vein patterning function were phenocopied by growth of WT in the presence of NPA. To address this question, we compared defects of *pin1,3;4;7* to those induced in WT by growth in the presence of 100 μM NPA, which is the highest concentration of NPA without toxic, auxin-efflux-unrelated effects (Petrasek et al., 2003; Dhonukshe et al., 2008). Because leaves develop more slowly at this concentration of NPA (Mattsson et al., 1999; Sieburth, 1999), to ensure maximal vascular differentiation we allowed them to grow for four weeks before analysis.

Consistent with previous reports (Mattsson et al., 1999; Sieburth, 1999), NPA only rarely induced leaf fusion in WT (see Fig. 6I for one such rare occurrence) but reproducibly induced characteristic vein-pattern defects: (1) the vein network comprised more lateral veins; (2) lateral veins failed to join the midvein but ran parallel to it to form a wide midvein; (3) lateral veins ended in a marginal vein that closely paralleled the leaf margin, lending a smooth outline to the vein network; (4) veins were thicker, though vascular elements were elongated and aligned along the length of the vein (Fig. 5A,D,E,H).

**Figure 5.**
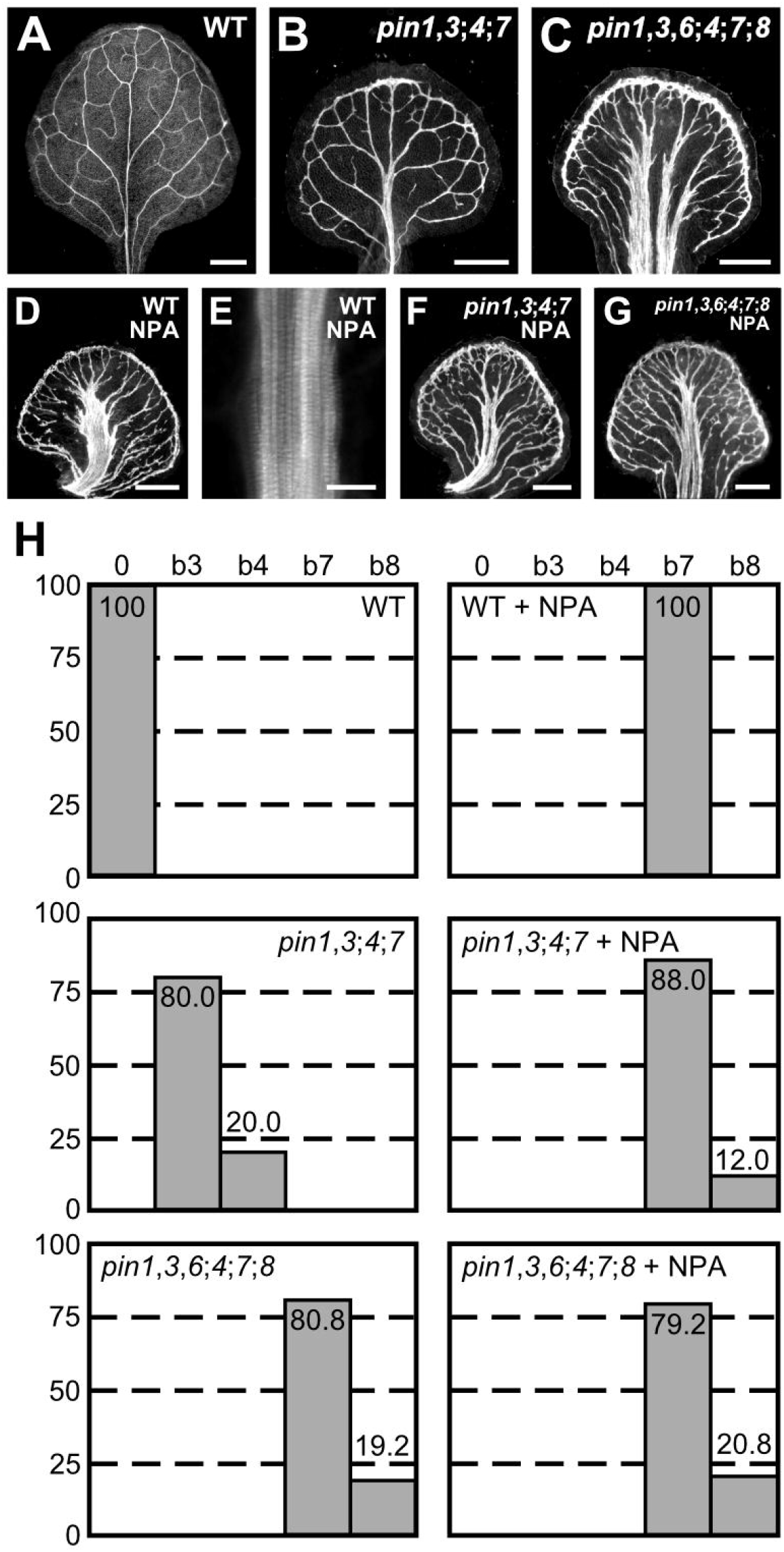
Genetic Versus Pharmacological Inhibition of Auxin Transport. (A–G) Top right: genotype and treatment. (A–G) Dark-field illumination (A–D,F,G) or confocal laser scanning microscopy (E) of mature first leaves. (A) WT. (B) *pin1-1,3;4;7*. (C) *pin1-1,3,6;4;7;8*. (D) NPA-grown WT. (E) Detail illustrating thick veins in NPA-grown WT (compare with Fig. 3J). (F) NPA-grown *pin1-1,3;4;7*. (G) NPA-grown *pin1-1,3,6;4;7;8*. (G) Percentages of leaves in phenotype classes (defined in Figures 2-4). Sample population sizes: WT, 38; *pin1-1,3;4;7*, 30; *pin1-1,3,6;4;7;8*, 73; NPA-grown WT, 41; NPA-grown *pin1-1,3;4;7*, 58; NPA-grown *pin1-1,3,6;4;7;8*, 48. Bars: (A-D,F,G) 0.5 mm, (E) 25 μm.

By contrast, 20% of *pin1,3;4;7* leaves were fused, and though *pin1,3;4;7* veins were thick, *pin1,3;4;7* vein patterns lacked all the other characteristic defects induced in WT by NPA (Fig. 5B,H). However, such defects were induced in *pin1,3;4;7* by NPA (Fig. 5F,H), suggesting that this background has residual NPA-sensitive vein-patterning activity. The vein pattern defects induced in WT or *pin1,3;4;7* by NPA were no different from those of *pin1,3,6;4;7;8* (Fig. 5C,D–F,H). Because no additional defects were induced in *pin1,3,6;4;7;8* by NPA (Fig. 5G,H), the residual NPA-sensitive vein-patterning activity of *pin1,3;4;7* is likely provided by *PIN6* and *PIN8*.

In conclusion, our results suggest that growth in the presence of NPA phenocopies defects of loss of *PIN*-dependent vein patterning function, that in the absence of this function any residual NPA-sensitive vein-patterning activity — if existing — becomes inconsequential, and that loss of neither *PIN*-dependent vein-patterning function nor NPA-sensitive vein-patterning activity phenocopies the vein pattern defects of *gn*.

### Contribution of *ABCB* Genes to Vein Patterning

Loss of *PIN*-dependent vein-patterning function or of NPA-sensitive vein-patterning activity fails to phenocopy the vein pattern defects of *gn* (Figure 2; Figure 5), suggesting that these latter are not the sole result of loss of *PIN*-dependent or NPA-sensitive polar auxin-transport induced by defects in coordination of PIN polar localization. However, it is possible that the vein pattern defects of *gn* result from additional or exclusive defects in another polar auxin-transport pathway; we asked whether that were so.

Cellular auxin efflux is catalyzed not only by PM-PIN proteins but by the PM-localized ATP-BINDING CASSETTE B1 (ABCB1) and ABCB19 proteins (Geisler et al., 2005; Bouchard et al., 2006; Petrasek et al., 2006; Blakeslee et al., 2007; Yang and Murphy, 2009), whose fusions to GFP (Dhonukshe et al., 2008; Mravec et al., 2008) are expressed at early stages of leaf development (Fig. 6A,B). We asked whether ABCB1/19-mediated auxin efflux were required for vein patterning.

**Figure 6.**
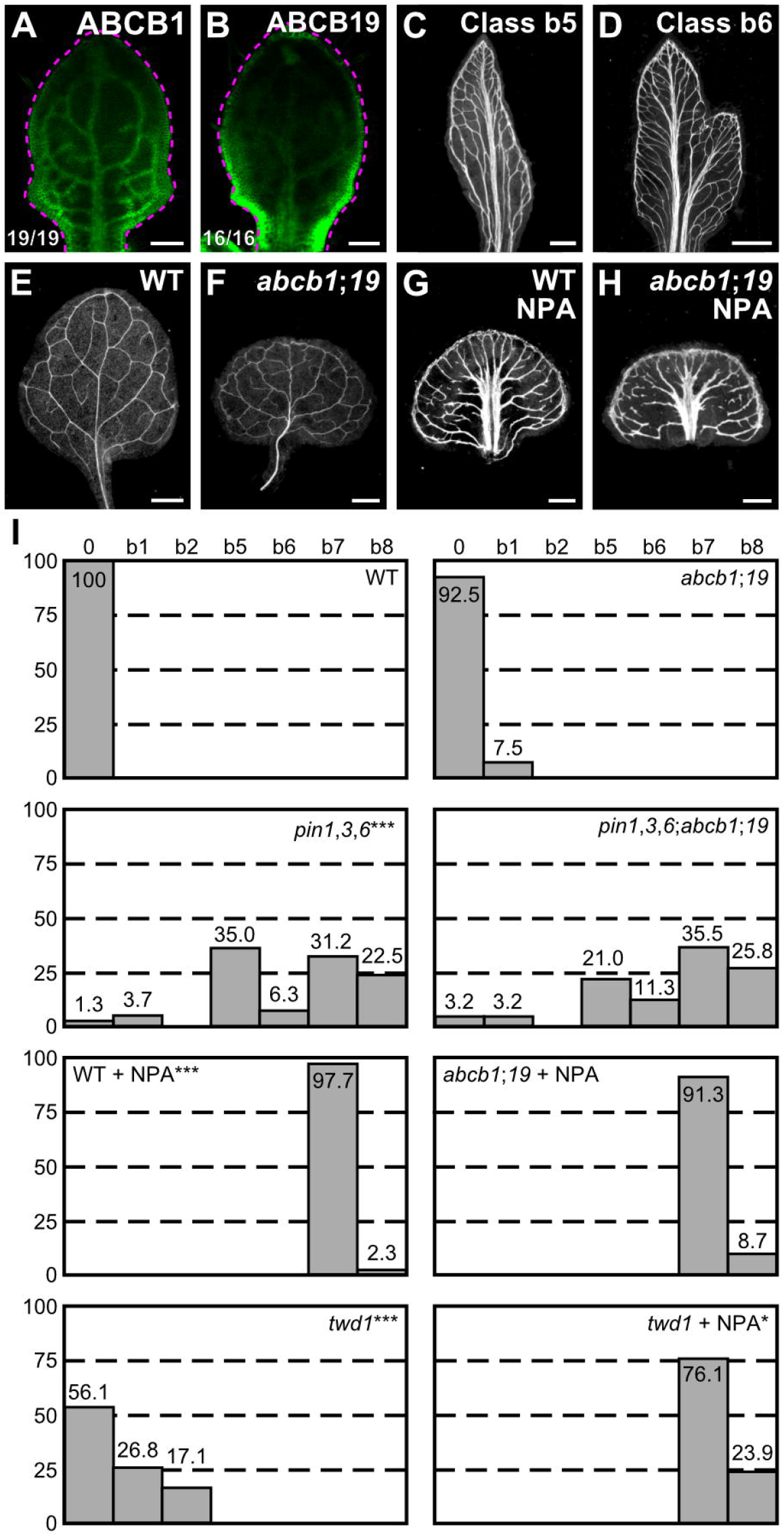
Contribution of *ABCB* Genes to Vein Patterning. (A,B,E–H) Top right: expression-reported gene, genotype and treatment. (A–B) Bottom left: reproducibility index. (A–B) Confocal laser scanning microscopy; 5-day-old first leaves. Dashed magenta line delineates leaf outline. (A) ABCB1::ABCB1:GFP expression. (B) ABCBI9::ABCBI9:GFP expression. (C–H) Dark-field illumination of mature first leaves. (C,D) Phenotype classes: class b5, thick veins and conspicuous marginal vein (C); class b6, fused leaves with thick veins and conspicuous marginal vein (D). (I) Percentages of leaves in phenotype classes (Classes 0, b1, b2, b7 and b8 defined in Figures 2-4). Difference between *pin1-1,3,6* and WT, between *twd1* and WT, and between NPA-grown WT and WT was significant at P<0.001 (***), and between NPA-grown *twd1* and NPA-grown WT was significant at P<0.05 (*) by Kruskal-Wallis and Mann-Whitney test with Bonferroni correction. Sample population sizes: WT, 41; *abcb1;19*, 40; *pin1-1,3,6*, 80; *pin1-1,3,6;abcb1;19*, 62; NPA-grown WT, 43; NPA-grown *abcb1;19*, 46; *twd1*, 41; NPA-grown *twd1*, 46. Bars: (A–B) 0.1 mm; (C–H) 0.5 mm.

The embryos of *abcb1* and *abcb19* were viable, but ~15% of *abcb1;19* embryos died during embryogenesis (Table S5); nevertheless, the vein patterns of *abcb1, abcb19* and *abcb1;19* were no different from the vein pattern of WT (Fig. 6E,F,I), suggesting that ABCB1/19-mediated auxin efflux is dispensable for vein patterning.

Developmental functions of ABCB1/19-mediated auxin transport overlap with those of PIN-mediated auxin transport (Blakeslee et al., 2007; Mravec et al., 2008). We therefore asked whether vein pattern defects resulting from simultaneous mutation of *PIN1, PIN3* and *PIN6*, or induced in WT by 100 μM NPA — which phenocopies loss of *PIN*-dependent vein-patterning function (Figure 5) — were enhanced by simultaneous mutation of *ABCB1* and *ABCB19*.

*pin1,3,6* embryos were viable (Table S6) and developed into seedlings (Table S7). The proportion of embryos derived from the self-fertilization of *PIN1,PIN3,PIN6*/*pin1,pin3,pin6;abcb1/abcb1;abcb19/abcb19* that died during embryogenesis was no different from the proportion of embryos derived from the self-fertilization of *abcb1/abcb1;abcb19/abcb19* that died during embryogenesis (Table S6), suggesting no nonredundant functions of *PIN1, PIN3* and *PIN6* in *ABCB1/ABCB19*-dependent embryo viability.

Consistent with previous reports (Blakeslee et al., 2007; Mravec et al., 2008), simultaneous mutation of *ABCB1* and *ABCB19* in the *pin1,3,6* background shifted the distribution of *pin1,3,6* cotyledon pattern phenotypes toward stronger classes (Figure S4). However, the spectrum of vein pattern phenotypes of *pin1,3,6;abcb1;19* was no different from that of *pin1,3,6*, and the vein pattern defects induced in *abcb1;19* by NPA were no different from those induced in WT by NPA (Fig. 6C,D,G–I), suggesting no vein-patterning function of *ABCB1* and *ABCB19* in the absence of function of *PIN1, PIN3* and *PIN6* or of NPA-sensitive, *PIN*-dependent vein-patterning function.

ABCB1 and ABCB19 are members of a large family (Geisler and Murphy, 2006); therefore, vein patterning functions of ABCB1/19-mediated auxin efflux might be masked by redundant functions provided by other ABCB transporters. The TWISTED DWARF1/ULTRACURVATA2 (TWD1/UCU2; TWD1 hereafter) protein (Kamphausen et al., 2002; Perez-Perez et al., 2004) is a positive regulator of ABCB-mediated auxin transport (Geisler et al., 2003; Bouchard et al., 2006; Bailly et al., 2008; Wu et al., 2010; Wang et al., 2013). Consistent with this observation, defects of *twd1* are more severe than, though similar to, those of *abcb1;19* (Geisler et al., 2003; Bouchard et al., 2006; Bailly et al., 2008; Wu et al., 2010; Wang et al., 2013). We therefore reasoned that analysis of *twd1* vein patterns might uncover vein patterning functions of ABCB-mediated auxin transport that could not be inferred from the analysis of *abcb1;19*.

Approximately 25% of *twd1* leaves had Y-shaped midveins and ~15% of *twd1* leaves were fused (Fig. 6I), suggesting possible vein-patterning functions of *TWD1*-dependent ABCB-mediated auxin transport. However, vein pattern defects induced in *twd1* by 100 μM NPA were no different from those induced in WT or *abcb1;19* by NPA (Fig. 6I), suggesting that vein patterning functions of *TWD1*-dependent ABCB-mediated auxin transport — if existing — become inconsequential in the absence of NPA-sensitive, *PIN*-dependent vein-patterning function. By contrast, NPA enhanced leaf separation defects of *twd1* (Fig. 6I), suggesting overlapping functions of *TWD1*-dependent ABCB-mediated auxin transport and NPA-sensitive, *PIN*-dependent auxin transport in leaf separation.

In conclusion, the residual vein patterning activity in *pin* mutants or in their NPA-induced phenocopy is not provided by *ABCB1, ABCB19* or *TWD1*-dependent ABCB-mediated auxin transport, and loss of PIN- and ABCB-mediated auxin transport fails to phenocopy vein pattern defects of *gn*.

### Contribution of *AUX1/LAX* Genes to Vein Patterning

Loss of PIN- and ABCB-mediated auxin transport fails to phenocopy vein pattern defects of *gn* (Figure 2; Figure 6), suggesting that these latter are not the sole result of loss of *PIN*-dependent, NPA-sensitive or *ABCB*-dependent polar auxin-transport. However, it is possible that the vein pattern defects of *gn* result from additional or exclusive defects in yet another auxin-transport pathway; we asked whether that were so.

Auxin is predicted to enter the cell by diffusion and through an auxin influx carrier (Rubery and Sheldrake, 1974; Raven, 1975). In Arabidopsis, auxin influx activity is encoded by the *AUX1, LAX1, LAX2* and *LAX3* (*AUX1/LAX*) genes (Parry et al., 2001; Yang et al., 2006; Swarup et al., 2008; Peret et al., 2012). We asked whether AUX1/LAX-mediated auxin influx were required for vein patterning.

*aux1;lax1;2;3* embryos were viable (Table S8). Because the vein patterns of *aux1;lax1;2;3* were no different from those of WT (Fig. 7A,C,D), we conclude that *AUX1/LAX* function is dispensable for vein patterning.

**Figure 7.**
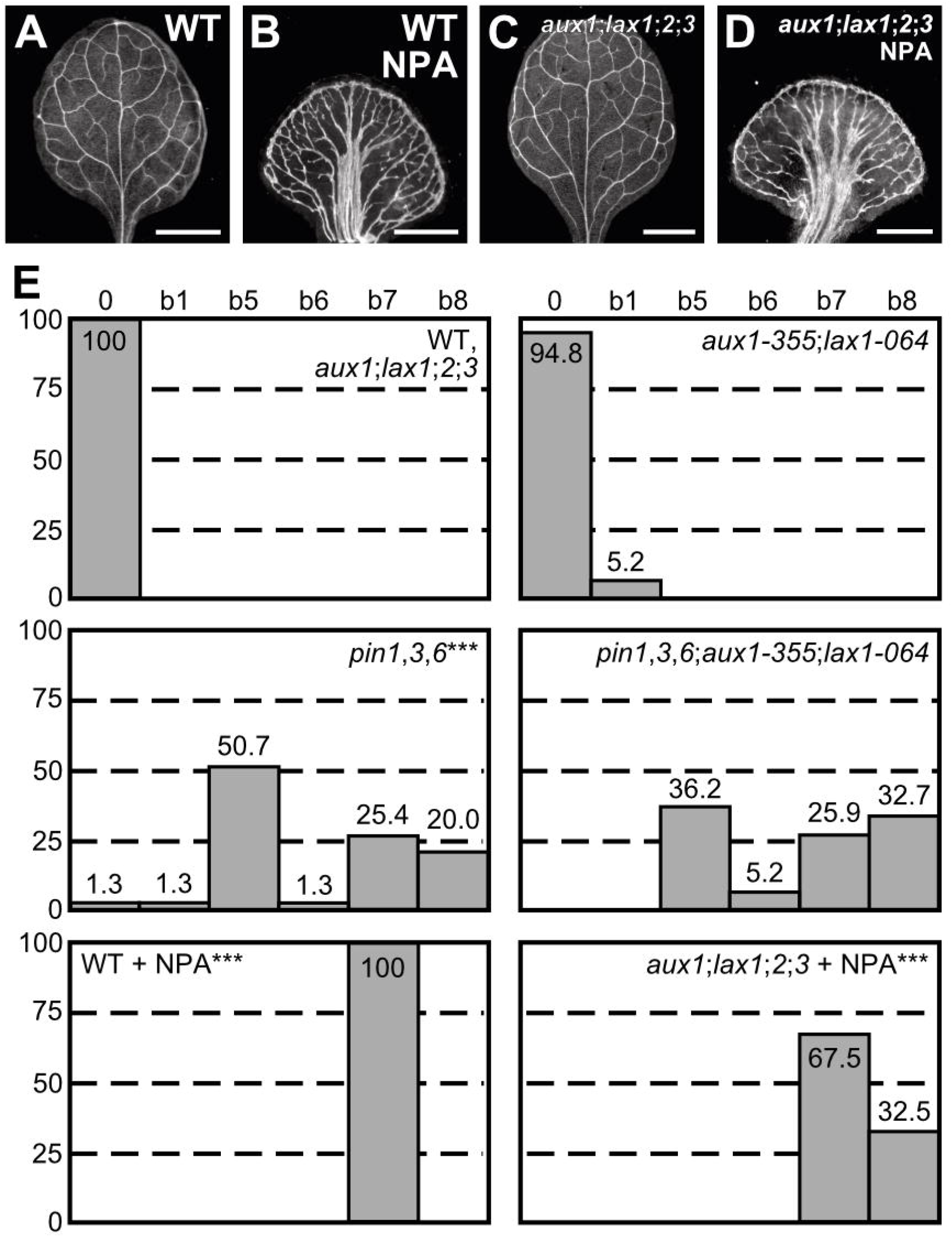
Contribution of *AUX1/LAX* Genes to Vein Patterning. (A–D) Dark-field illumination of mature first leaves. Top right: genotype and treatment. (E) Percentages of leaves in phenotype classes (defined in Figures 2-4 and 6). Difference between *pin1-1,3,6* and WT, between NPA-grown WT and WT, and between NPA-grown *aux1-21;lax1;2;3* and NPA-grown WT was significant at P<0.001 (***) by Kruskal-Wallis and Mann-Whitney test with Bonferroni correction. Sample population sizes: WT, 53; *aux1-21;lax1;2;3*, 60; *aux1-355;lax1-064*, 77; *pin1-1,3;6*, 75; *pin1-1,3,6;aux1-355;lax1-064*, 58; NPA-grown WT, 46; NPA-grown *aux1-21;lax1;2;3*, 40. Bars: (A–D) 1 mm.

We next asked whether contribution of *AUX1/LAX* genes to vein patterning only became apparent in conditions of extremely reduced PIN-mediated auxin transport. To address this question, we tested whether vein pattern defects resulting from simultaneous loss of function of *PIN1, PIN3* and *PIN6* or induced in WT by 100 μM NPA, which phenocopies simultaneous mutation of all the *PIN* genes with vein patterning function (Figure 4), were enhanced by simultaneous mutation of *AUX1* and *LAX1* — the two *AUX1/LAX* genes that most contribute to shoot organ patterning (Bainbridge et al., 2008) — or of all *AUX1/LAX* genes, respectively.

The embryos derived from the self-fertilization of *PIN1,pin3,PIN6/pin1,pin3,pin6;aux1/aux1;lax1/lax1* were viable (Table S9) and developed into seedlings (Table S10). The spectrum of vein pattern phenotypes of *pin1,3,6;aux1;lax1* was no different from that of *pin1,3,6* and the vein pattern defects induced in *aux1;lax1;2;3* by NPA were no different from those induced in WT by NPA (Fig. 7B,D,E), suggesting no vein-patterning function of *AUX1/LAX* genes in conditions of extremely reduced auxin transport. On the other hand, simultaneous mutation of *AUX1* and *LAX1* in the *pin1,3,6* background shifted the distribution of *pin1,3,6* cotyledon pattern phenotypes toward stronger classes (Figure S4), and NPA induced leaf fusion in *aux1;lax1;2;3* but not in WT (Fig. 7E), suggesting that AUX1/LAX-mediated auxin influx and NPA-sensitive, *PIN*-dependent, auxin transport have overlapping functions in cotyledon and leaf separation and that — consistent with previous observations (Reinhardt et al., 2003; Bainbridge et al., 2008; Kierzkowski et al., 2013) — AUX1/LAX-mediated auxin influx contributes to maintaining cotyledon and leaves separate in conditions of reduced auxin transport. Nevertheless, loss of PIN- and AUX1/LAX-mediated auxin transport fails to phenocopy the vein pattern defects of *gn*.

### Genetic Interaction Between *GN* and *PIN* Genes

The vein pattern defects of *gn* are not the sole result of loss of *PIN*-dependent auxin transport (Figure 2; Figure 4; Figure 5); however, they could be the result of abnormal polarity of PIN-mediated auxin transport induced by defects in coordination of PIN polar localization. Were that so, the vein pattern defects of *gn* would depend on *PIN* genes, and therefore the vein pattern defects of *gn;pin* mutants would resemble those of *pin* mutants; we tested whether that were so.

We first asked what the phenotype were of the quintuple mutant between the strong allele *gn-13* (Figure 2) and mutation in *PIN1, PIN3, PIN4* and *PIN7* — i.e. the *PM-PIN* genes with vein patterning function (Figure 3).

Consistent with previous observations (Mayer et al., 1993; Shevell et al., 1994), in *gn* seedlings hypocotyl and root were replaced by a basal peg, and the cotyledons were most frequently fused (Fig. S5A,C; Fig S6B; Fig. S7A,B). As shown above (Fig. S2A,B; Fig. S3A,H), *pin1,3;4;7* seedlings had hypocotyl, short root, and a single cotyledon or two — either separate or fused — cotyledons (Fig. S5A,B; Fig S6C,D; Fig. S7B).

*gn;pin1,3;4;7* embryos were viable (Table S11). A novel phenotype segregated in approximately one-sixteenth of the progeny of plants homozygous for *pin3, pin4* and *pin7* and heterozygous for *pin1* and *gn* — no different from the one-sixteenth frequency expected for the *gn;pin1,3;4;7* homozygous mutants by Pearson’s chi-squared (χ^2^) goodness-of-fit test (Table S12). We genotyped 10 of the seedlings with the novel mutant phenotype and found they were *gn;pin1,3;4;7* homozygous mutants. *gn;pin1,3;4;7* seedlings had hypocotyl, no root and the cotyledons were fused (Fig. S5A,D; Fig S6E; Fig. S7B).

WT cotyledons have a I-shaped midvein and three or four loops (Fig. S8A,B,K). All the veins of *pin1,3;4;7* cotyledons were thick, and all *pin1,3;4;7* cotyledons had three or four loops (Fig. S8C,D,K). In *pin1,3;4;7* cotyledons, the proximal end of the first loops joined the midvein more basally than in WT, and minor veins branched from midvein and loops (Fig. S8C,D,K). Approximately 60% of *pin1,3;4;7* cotyledons had an I-shaped midvein, while the remaining ~40% of them had a Y-shaped midvein (Fig. S8C,D,K).

Consistent with previous observations (Mayer et al., 1993; Shevell et al., 1994), in ~70% of *gn* cotyledons short stretches of vascular elements connected the proximal side of a central, shapeless cluster of seemingly randomly oriented vascular elements with the basal part of the cotyledon, while vascular differentiation was limited to a central, shapeless vascular cluster in the remaining ~30% of *gn* cotyledons (Fig. S8F,G,K). The vein pattern defects of *gn;pin1,3;4;7* cotyledons were no different from those of *gn*; cotyledons (Fig. S8H,K), suggesting that the vein pattern phenotype of *gn* cotyledons is epistatic to that of *pin1,3;4;7* cotyledons. Likewise, the vein pattern defects of *gn;pin1,3;4;7* leaves were no different from those of *gn* leaves (Fig. 8A,B,E), suggesting that the vein pattern phenotype of *gn* leaves is epistatic to that of *pin1,3;4;7* leaves.

**Figure 8.**
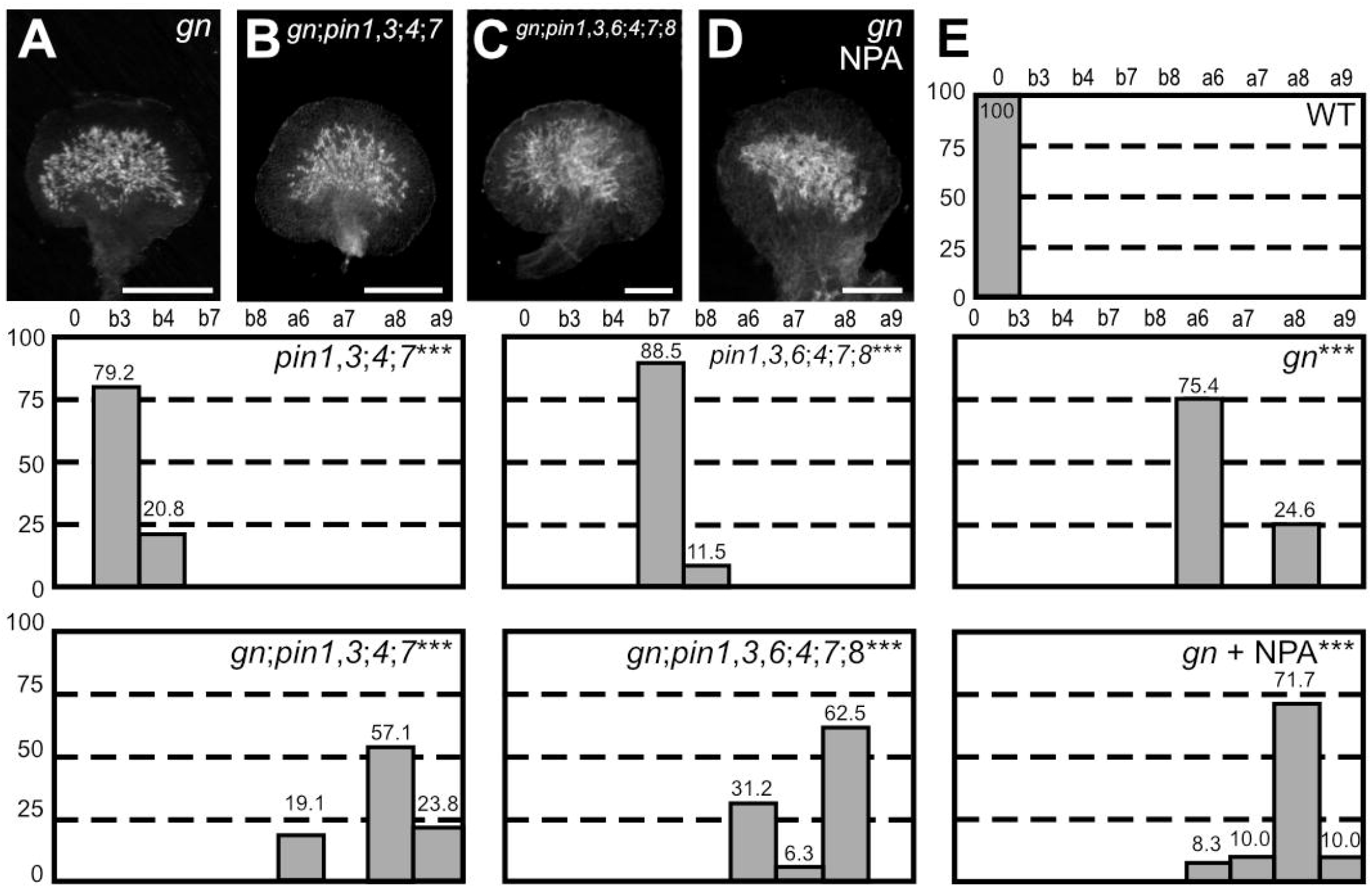
Genetic Interaction Between *GN* and *PIN* Genes. (A–D) Dark-field illumination of mature first leaves. Top right: genotype and treatment. (E) Percentages of leaves in phenotype classes (defined in Figures 2-4). Difference between *pin1-1,3;4;7* and WT, between *pin1-1,3,6;4;7;8* and WT, between *gn* and WT, between *gn-13;pin1-1,3;4;7* and *pin1-1,3;4;7*, between *gn-13;pin1-1,3;6;4;7;8* and *pin1-1,3,6;4;7;8* and between NPA-grown *gn-13* and *pin1-1,3,6;4;7;8*, was significant at P<0.001 (***) by Kruskal-Wallis and Mann-Whitney test with Bonferroni correction. Sample population sizes: WT, 63; *pin1-1,3;4;7*, 53; *pin1-1,3,6,4;7;8*, 52; *gn-13*, 69; *gn-13;pin1-1,3;4;7*, 21; *gn-13;pin1-1,3,6;4;7;8*, 15; NPA-grown *gn-13*, 60. Bars: (A–D) 0.5 mm.

We next asked what the phenotype were of the septuple mutant between the strong allele *gn-13* (Figure 2) and mutation in all the *PIN* genes with vein patterning function (Figure 4).

As shown above (Fig. S2A,D; Fig. S3A,H), *pin1,3,6;4;7;8* seedlings had hypocotyl, short root and a single cotyledon or two fused cotyledons (Fig. S5A,E; Fig S6G,H; Fig. S7B).

*gn;pin1,3,6;4;7;8* embryos were viable (Table S11). A phenotype similar to that of *gn;pin1,3;4;7* segregated in approximately one-sixteenth of the progeny of plants homozygous for *pin3, pin4, pin6, pin7* and *pin8* and heterozygous for *pin1* and *gn* — no different from the one-sixteenth frequency expected for the *gn;pin1,3,6;4;7;8* homozygous mutants by Pearson’s χ^2^ goodness-of-fit test (Table S12). We genotyped 10 of the seedlings with the novel mutant phenotype and found they were *gn;pin1,3,6;4;7;8* homozygous mutants. Like *gn;pin1,3;4;7* seedlings, *gn;pin1,3,6;4;7;8* seedlings had hypocotyl and no root, but unlike *gn;pin1,3;4;7* seedlings ~90% of *gn;pin1,3,6;4;7;8* seedlings had completely fused cup-shaped cotyledons (Fig. S5A,F; Fig S6I; Fig. S7B).

The vein pattern defects of *pin1,3,6;4; 7;8* cotyledons were similar to those of *pin1,3;4;7* cotyledons, but in ~85% of *pin1,3,6;4;7;8* cotyledons the loops joined the midvein at the base of the cotyledon and the top half of the vein network outline was thick (Fig. S8C-E,K). The vein pattern defects of *gn;pin1,3,6;4;7;8* cotyledons were no different from those of *gn* cotyledons (Fig. S8I-K), suggesting that the vein pattern phenotype of *gn* cotyledons is epistatic to that of *pin1,3,6;4;7;8* cotyledons. Likewise, the vein pattern defects of *gn;pin1,3,6;4;7;8* leaves were no different from those of *gn* leaves (Fig. 8C,E), suggesting that the vein pattern phenotype of *gn* leaves is epistatic to that of *pin1,3,6;4;7;8* leaves. Finally, 100 μM NPA, which phenocopies loss of *PIN*-dependent vein-patterning function (Figure 5), failed to induce additional vein pattern defects in *gn* leaves (Fig. 8D,E).

In conclusion, our results suggest that the vein pattern defects of *gn* are not the result of either the sole loss of PIN-mediated auxin transport or the sole abnormal polarity of PIN-mediated auxin transport induced by defects in coordination of PIN polar localization.

### Response of *pin* Leaves to Auxin Application

The uniform vein-pattern phenotype of *pin1,3,6;4;7;8* was phenocopied by growth of WT in the presence of high concentration of NPA (Figure 5). Moreover, the vein-pattern phenotype of *pin1,3,6;4;7;8* was unchanged by NPA treatment, and the NPA-induced vein-pattern phenocopy of *pin1,3,6;4;7;8* was unchanged by mutation in any other known intercellular auxin-transporter (Figure 6; Figure 7). These observations suggest that the function of known intercellular auxin-transporters in vein patterning is dispensable in the absence of the auxin transport activity of PIN1, PIN3, PIN4, PIN6, PIN7 and PIN8. Because auxin transport is thought to be essential for auxin-induced vascular-strand formation (reviewed in (Sachs, 1981; Berleth et al., 2000; Aloni, 2010; Sawchuk and Scarpella, 2013)), we asked whether auxin-induced vein formation in *pin1,3,6;4;7;8* and consequently whether veins were formed by an auxin-dependent mechanism in *pin1,3,6;4;7;8*. To address this question, we applied lanolin paste containing 1% of the natural auxin indole-3-acetic acid (IAA) to one side of developing leaves of WT and *pin1,3,6;4;7;8*, and recorded tissue response in mature leaves.

Consistent with previous reports (Scarpella et al., 2006; Sawchuk et al., 2007), IAA induced formation of extra veins in ~70% of WT leaves (27/38) (Fig. 9A,B), while ~30% of WT leaves (9/38) failed to respond to IAA application.

**Figure 9.**
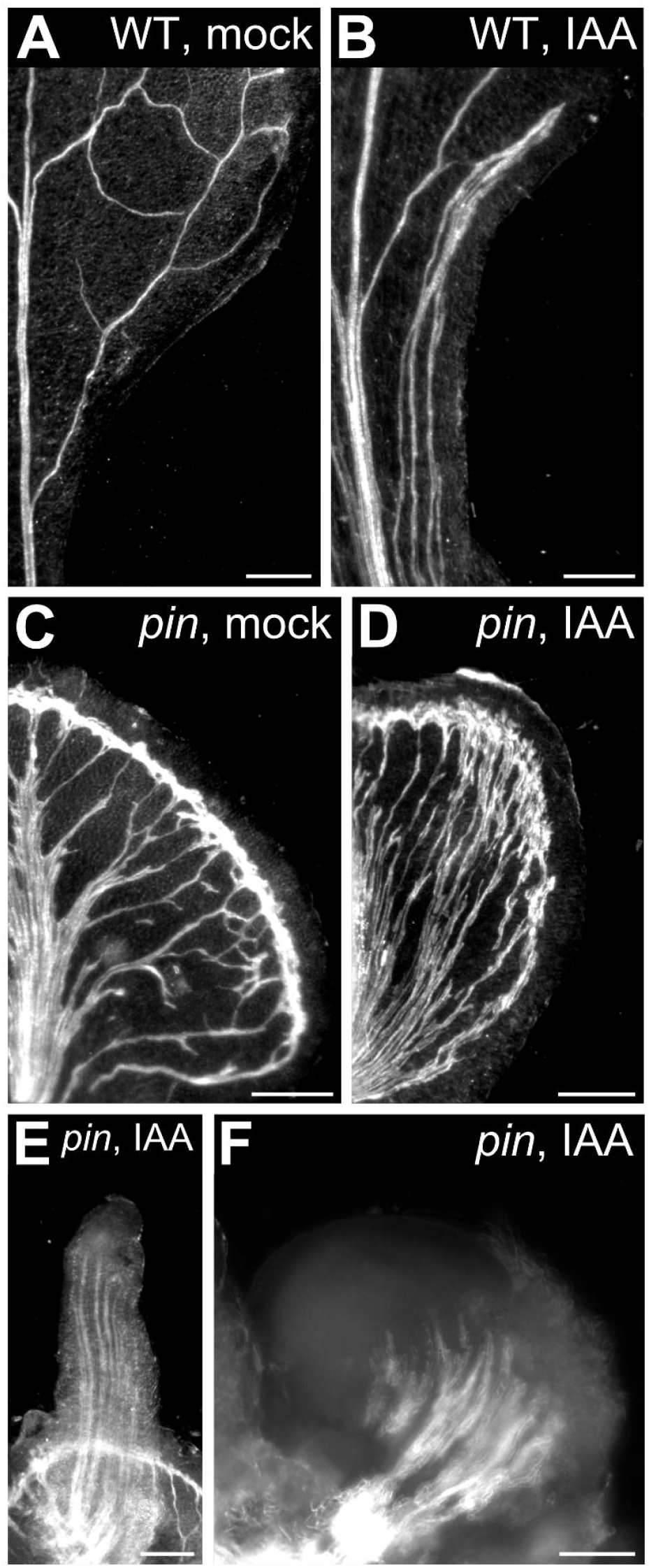
Response of *pin* Leaves to Auxin Application. (A–F) Top right: genotype and treatment. Dark-field illumination of mature first leaves of WT (A,B) or *pin1-1,3,6;4;7;8* (C–F) at side of application of lanolin paste (A,C) or lanolin paste containing 1% IAA (B,D–F). Bars: (A) 0.5 mm; (B–E) 0.25 mm; (F) 0.1 mm.

The effects of IAA on *pin1,3,6;4;7;8* leaves were variable. In 40% of the leaves (28/70), IAA induced formation of extra veins (Fig. 9C,D). In ~60% of the leaves in which IAA induced formation of extra veins (17/28), IAA also induced tissue outgrowth of varied shape (Fig. 9E,F). In 30% of *pin1,3,6;4;7;8* leaves (21/70), IAA induced tissue outgrowth but failed to induce formation of extra veins in the leaf; however, in nearly 80% of the *pin1,3,6;4;7;8* leaves in which IAA induced tissue outgrowth [30/(17+21)=30/38], IAA also induced formation of vascular strands in the outgrowth (Fig. 9E,F). Finally, as in WT, 30% of *pin1,3,6;4;7;8* leaves (21/70) failed to respond to IAA application in any noticeable way.

We conclude that *pin1,3,6;4;7;8* leaves respond to vein-formation-inducing auxin signals and consequently that veins are formed by an auxin-dependent mechanism in the absence of PIN-mediated auxin transport.

### Contribution of Auxin Signaling to Vein Patterning

Leaves of *pin1,3,6;4;7;8* respond to vein-formation-inducing auxin signals (Figure 9), suggesting that the residual vein-patterning activity in those leaves may be provided by an auxin-dependent mechanism. We therefore asked what the contribution of auxin signaling to vein patterning were in the absence of *PIN*-dependent vein patterning activity.

To address this question, we used mutants in *AUXIN-RESISTANT1* (*AXR1*), which lack a required post-translational modification of the auxin receptor complex (reviewed in (Calderon-Villalobos et al., 2010; Schwechheimer, 2018)); double mutants in *TRANSPORT INHIBITOR RESPONSE1* (*TIR1*) and *AUXIN SIGNALING F-BOX2* (*AFB2*), which lack the two auxin receptors that most contribute to auxin signaling (Dharmasiri et al., 2005); and phenylboronic acid (PBA), which inhibits auxin signaling (Matthes and Torres-Ruiz, 2016).

The embryos of *axr1* and *tir1;afb2* were viable (Table S13). In ~40–65% of the leaves of *axr1*, of *tir1;afb2* and of WT grown in the presence of 10 μM PBA — as in leaves of weak *gn* alleles (Figure 2) — loops were open (Fig. 10A,B,H). Furthermore, in ~20–50% of the leaves of *axr1*, of *tir1;afb2* and of WT grown in the presence of 10 μM PBA — again as in leaves of weak *gn* alleles (Figure 2) — veins were fragmented (Fig. 10A,B,H).

**Figure 10.**
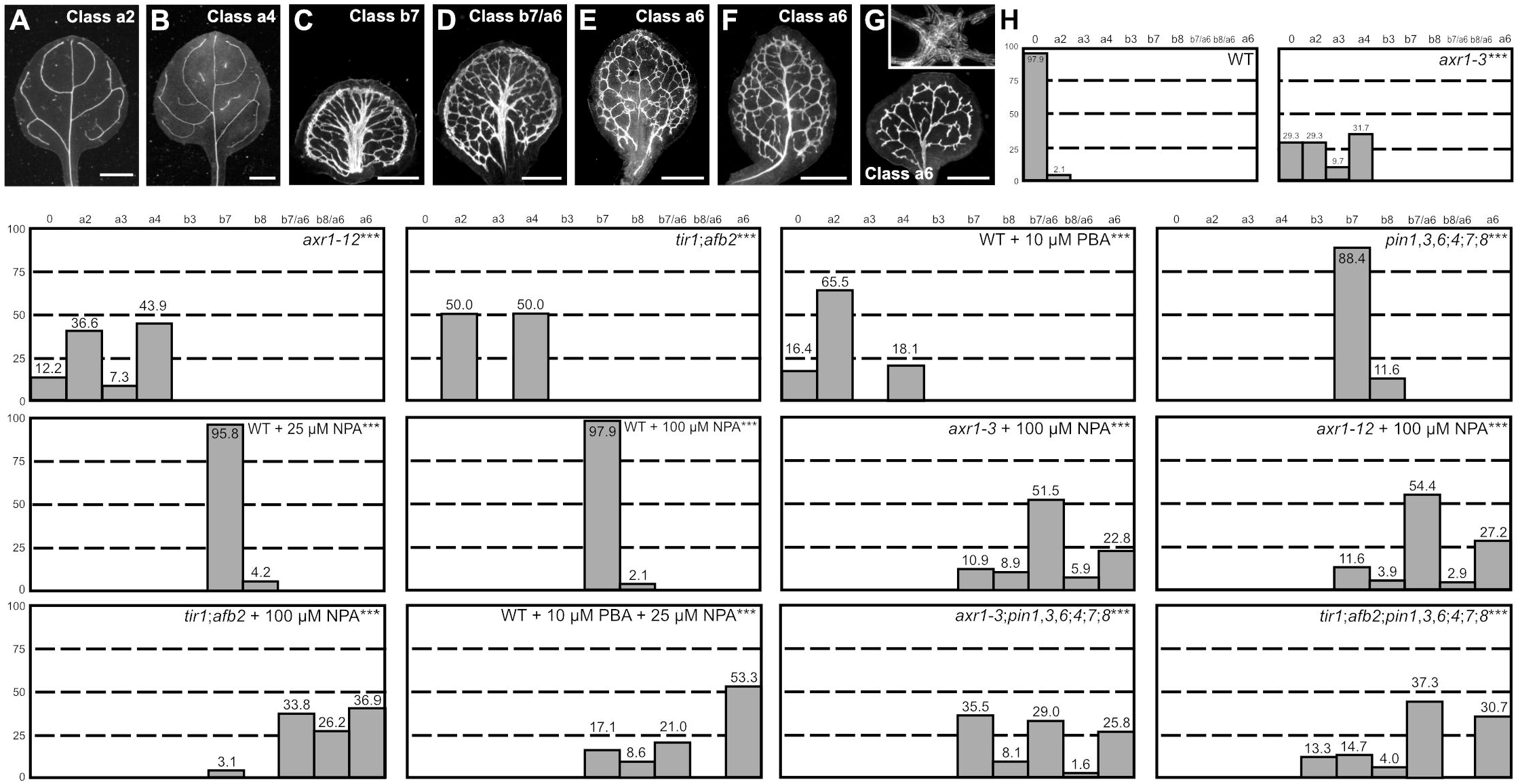
Contribution of Auxin Signaling to Vein Patterning. (A–G) Dark-field illumination of mature leaves illustrating phenotype classes (A–F, top right; G, bottom left): class a2 (*axr1-3*; A); class a4 (*tir1;afb2*; B); class b7 (NPA-grown WT; C); class b7/a6, wide midvein, more lateral-veins, dense network of thick veins and conspicuous marginal vein (NPA-grown *axr1-12*; D); class b8/a6, fused leaves with wide midvein, more lateral-veins, dense network of thick veins and conspicuous marginal vein (not shown); class a6 (E: PBA- and NPA-grown WT; F: NPA-grown *tir1;afb2*; G: *tir1;afb2;pin1-1,3,6;4;7;8*); inset in (G) illustrates cluster of seemingly randomly oriented vascular elements. (H) Percentages of leaves in phenotype classes (Classes 0, a2, a3, a4, a6, b7 and b8 defined in Figures 2 and 4). Difference between *axr1-3* and WT, between *axr1-12* and WT, between *tir1;afb2* and WT, between PBA-grown WT and WT, between *pin1-1,3,6;4;7;8* and WT, between NPA-grown WT and WT, between NPA-grown *axr1-3* and NPA-grown WT, between NPA-grown *axr1-12* and NPA-grown WT, between NPA-grown *tir1;afb2* and NPA-grown WT, between PBA- and NPA-grown WT and NPA-grown WT, between *axr1-3;pin1-1,3,6;4;7;8*, and *pin1-1,3,6;4;7;8* and between *tir1;afb2;pin1-1,3,6;4;7;8* and *pin1-1,3,6;4;7;8* was significant at *P*<0.001 (***) by Kruskal-Wallis and Mann-Whitney test with Bonferroni correction. Sample population sizes: WT, 47; *axr1-3*, 41; *axr1-12*, 41; *tir1;afb2*, 42; PBA-grown WT, 58; *pin1-1,3,6;4;7;8*, 63; NPA-grown WT, 48 (25 μM) or 146 (100 μM); NPA-grown *axr1-3*, 101; NPA-grown *axr1-12*, 103; NPA-grown *tir1;afb2*, 65; PBA- and NPA-grown WT, 105; *axr1-3;pin1-1,3,6;4;7;8*, 62; *tir1;afb2;pin1-1,3,6;4;7;8*, 75. Bars: (A,B) 1 mm; (C–E) 0.75 mm (F,G) 0.5 mmm.

We next asked whether PBA, mutation of *AXR1* or simultaneous mutation of *TIR1* and *AFB2* enhanced the vein pattern defects induced by NPA, which phenocopies loss of *PIN*-dependent vein-patterning activity (Figure 5).

Approximately 3-25% of the leaves of NPA-grown *axr1*, NPA-grown *tir1;afb2* and NPA- and PBA-grown WT resembled those of NPA-grown WT or of *pin1,3,6;4;7;8* (Fig. lOC,H). However, ~25-50% of the leaves of NPA-grown *axr1*, NPA-grown *tir1;afb2* and NPA- and PBA-grown WT resembled those of intermediate *gn* alleles: veins were thicker; the vein network was denser; and its outline was jagged because of narrow clusters of vascular elements that were oriented perpendicular to the leaf margin and that were laterally connected by veins or that, in the most severe cases, were aligned in seemingly random orientations (Figure 2; Fig. 10E,F,H; Fig. 10G, inset). Finally, ~20-60% of the leaves of NPA-grown *axr1*, NPA-grown *tir1;afb2* and NPA- and PBA-grown WT had features intermediate between those of NPA-grown WT or of *pin1,3,6;4;7;8* and those of intermediate *gn* alleles (Fig. 10D,H).

We next asked whether the spectrum of vein pattern defects of NPA-grown *axr1* and *tir1;afb2* were recapitulated by *axr1;pin1,3,6;4;7;8* and *tir1;afb2;pin1,3,6;4;7;8*.

*axr1;pin1,3,6;4;7;8* embryos were viable (Table S14) and developed into seedlings (Table S15) that resembled *pin1,3,6;4;7;8* seedlings (Figure S9; Figure S10). Also *tir1;afb2;pin1,3,6;4;7;8* embryos were viable (Table S14), but they developed into seedlings (Table S15) whose cotyledon pattern defects were more severe than those of *pin1,3,6;4;7;8* seedlings (Figure S10; Figure S11) and whose root was replaced by a basal peg (Fig. S11C), as in strong *gn* alleles (Mayer et al., 1993) (Fig. S6B). Nevertheless, the spectrum of vein pattern defects of *axr1;pin1,3,6;4;7;8* and *tir1;afb2;pin1,3,6;4;7;8* was no different from that of NPA-grown *axr1* and NPA-grown *tir1;afb2* (Fig. 10C–H).

These observations suggest that the residual vein-patterning activity in *pin1,3,6;4;7;8* is provided, at least in part, by AXR1- and TIR1/AFB2-mediated auxin signaling. Because reduction of AXR1- and TIR1/AFB2-mediated auxin signaling synthetically enhanced vein pattern defects resulting from loss of *PIN*-dependent vein-patterning function, we conclude that PIN-mediated auxin transport and AXR1- and TIR1/AFB2-mediated auxin signaling provide overlapping functions in vein patterning. Finally, the similarity between the vein pattern defects of NPA-grown *axr1* and *tir1;afb2*, of NPA- and PBA-grown WT, and of *axr1;pin1,3,6;4;7;8* and *tir1;afb2;pin1,3,6;4;7;8*, on the one hand, and those of intermediate *gn* alleles, on the other, suggests that the vein pattern defects of *gn* are caused by simultaneous defects in auxin transport and signaling.

### Contribution of *GN* to Auxin Signaling

Were the vein pattern defects of *gn* not only the result of abnormal polarity or loss of PIN-mediated auxin transport but that of defects in auxin signaling, the vein pattern defects of *gn* might be associated with reduced auxin response, and the reduced auxin response of *gn* would be recapitulated by NPA-grown *axr1*; we asked whether that were so.

To address this question, we imaged expression of the auxin response reporter DR5rev::nYFP (Heisler et al., 2005; Sawchuk et al., 2013) in developing first-leaves of WT, *pin1,3,6;4;7;8*, NPA-grown WT, *axr1, gn* and NPA-grown *axr1*.

As previously shown (Sawchuk et al., 2013; Verna et al., 2015), strong DR5rev::nYFP expression was mainly associated with developing veins in WT (Fig. 11A). In *pin1,3,6;4;7;8* and NPA-grown WT, DR5rev::nYFP expression was weaker and mainly confined to areas near the margin of the leaf (Fig. 11B–E). DR5rev::nYFP expression was weaker also in *axr1* but was still associated with developing veins (Fig. 11F,G). Finally, in both *gn* and NPA-grown *axr1*, DR5rev::nYFP expression was much weaker and scattered across large areas of the leaf (Fig. 11H-K), suggesting that the vein pattern defects of *gn* are associated with reduced auxin response and that the reduced auxin response of *gn* is recapitulated by NPA-grown *axr1*.

**Figure 11.**
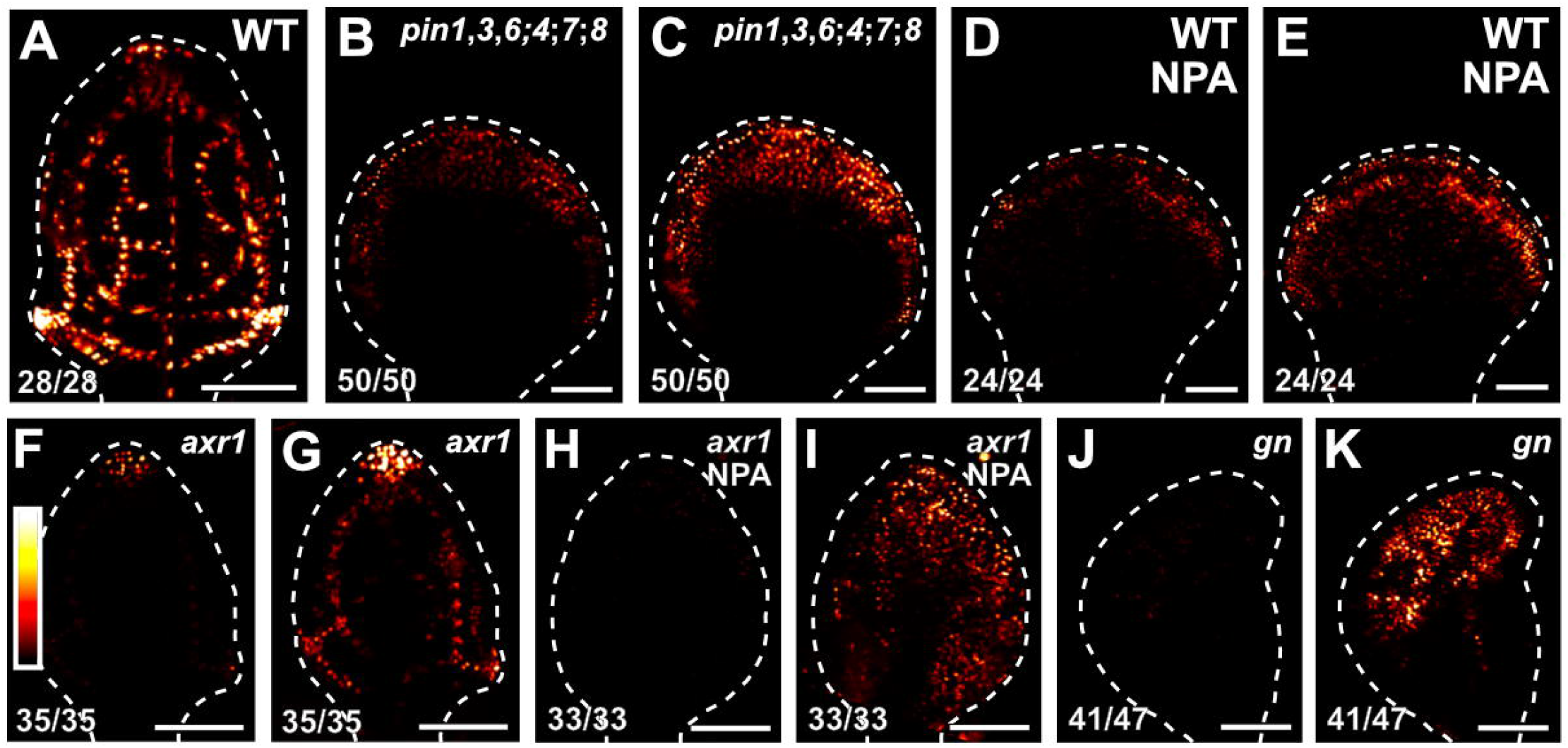
Auxin Response in Developing Leaves. (A–K) Confocal laser scanning microscopy; first leaves 4 (A,D,E), 5 (B,C,F–I) or 6 (J,K) days after germination. DR5rev::nYFP expression; look-up table (ramp in F) visualizes expression levels. Top right: genotype and treatment. Bottom left: reproducibility index. Dashed white line delineates leaf outline. Images in A,B,D,F,H,J were taken at identical settings. Images in A,C,E,G,I,K were taken by matching signal intensity to detector’s input range (~5% saturated pixels). Bars: (A–K) 100 μm.

Were the vein pattern defects of *gn* caused by simultaneous defects in auxin transport and signaling and did *GN* control auxin signaling as it controls auxin transport, the vein pattern defects of *gn;axr1* should resemble those of *gn*, just as the vein pattern defects of *gn;pin1,3;4;7* and *gn;pin1,3,6;4;7;8* resemble those of *gn*; we tested whether that were so.

*gn;axr1* embryos were viable (Table S16) and developed into seedlings (Table S17) that resembled *gn* seedlings (Figure S12; Figure S13), and the vein pattern defects of *gn;axr1* were no different from those of *gn* (Fig. 12A–C), suggesting that the phenotype of *gn* is epistatic to that of *axr1*.

**Figure 12.**
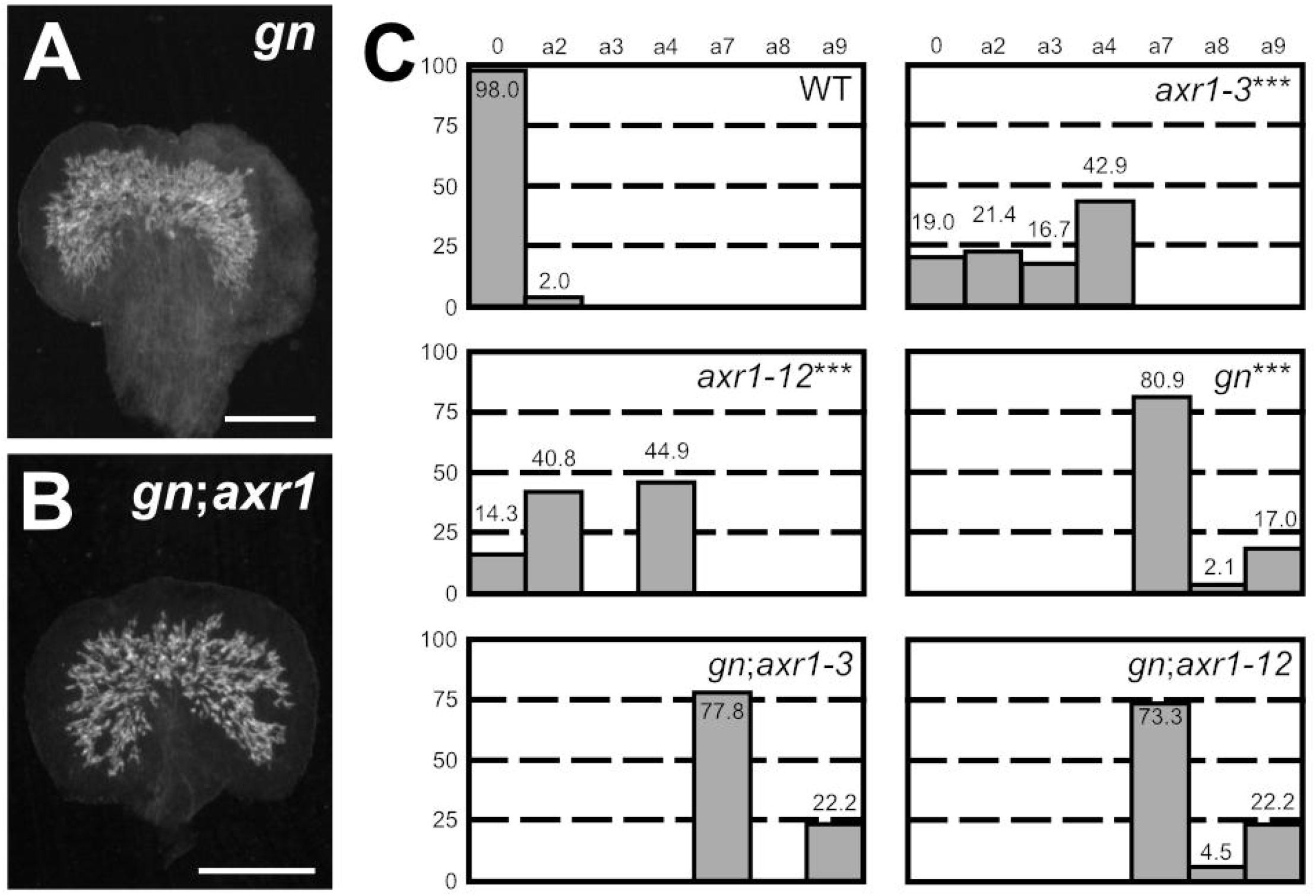
Genetic Interaction Between *GN* and *AXR1*. (A,B) Dark-field illumination of mature first leaves. Top right: genotype. (C) Percentages of leaves in phenotype classes (defined in Figures 2). Difference between *axr1-3* and WT, between *axr1-12* and WT, and between *gn-13* and WT was significant at *P*<0.001 (***) by Kruskal-Wallis and Mann-Whitney test with Bonferroni correction. Sample population sizes: WT, 49; *axr1-3*, 42; *axr1-12*, 49; *gn-13*, 47; *gn-13;axr1-3*, 45; *gn-13;axr1-12*, 45. Bars: (A,B) 0.75 mm.

We conclude that the vein pattern defects of *gn* are caused by simultaneous defects in auxin transport and signaling and that *GN* controls both auxin signaling and auxin transport.

### Contribution of Auxin Transport and Signaling to Coordination of Tissue Cell Polarity During Vein Formation

The vein pattern defects of *gn* are caused by simultaneous defects in auxin transport and signaling. We finally asked whether simultaneous defects in auxin transport and signaling recapitulated *gn* defects in coordination of tissue cell polarity.

To address this question, we imaged cellular localization of PIN1::PIN1:GFP expression during first-leaf development in WT, *tir1;afb2*, NPA-grown WT, NPA-grown *tir1;afb2*, and *gn^van7^*.

Consistent with previous reports (Benkova et al., 2003; Reinhardt et al., 2003; Heisler et al., 2005; Scarpella et al., 2006; Wenzel et al., 2007; Bayer et al., 2009; Sawchuk et al., 2013; Marcos and Berleth, 2014; Verna et al., 2015), and as shown above (Fig. 1P,T), in the cells of the second loop at early stages of its development in WT leaves, PIN1::PIN1:GFP expression was mainly localized to the basal side of the PM, toward the midvein; in the inner cells flanking the developing loop, PIN1::PIN1:GFP expression was mainly localized to the side of the PM facing the developing loop; and in the inner cells further away from the developing loop, PIN1::PIN1:GFP expression was localized isotropically, or nearly so, at the PM (Fig. 13B). At later stages of second-loop development, by which time PIN1::PIN1:GFP expression had become restricted to the sole, elongated cells of the developing loop, PIN1::PIN1:GFP expression was localized to the basal side of the PM, toward the midvein (Fig. 13H). We observed a similar pattern of localization of PIN1::PIN1:GFP expression in *tir1;afb2*, but in this background stages of second-loop development comparable to those in WT appeared at later stages of leaf development, and nearly 70% (24/35) of second loops failed to connect to the first loop (Fig. 13C,I).

**Figure 13.**
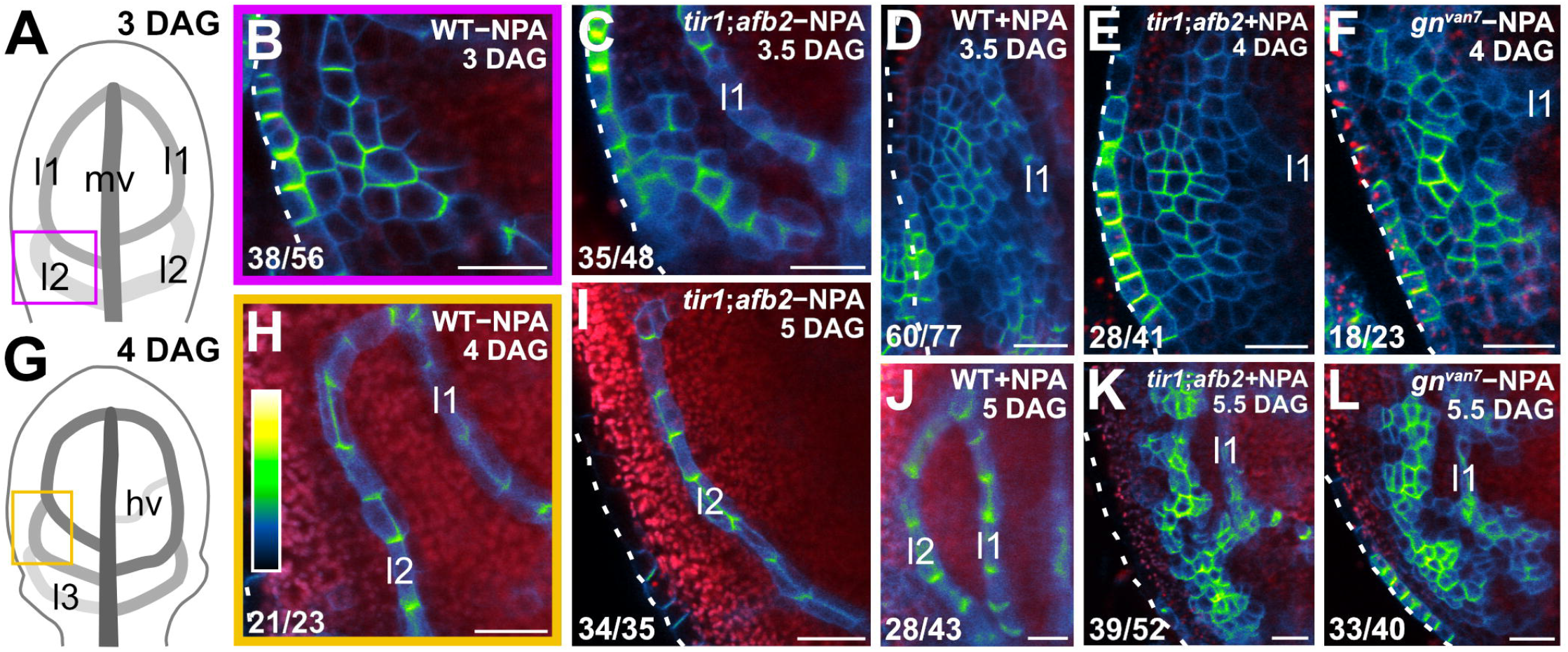
Contribution of Auxin Transport and Signaling to Coordination of Tissue Cell Polarity During Vein Formation. (A,G) Increasingly darker grays depict progressively later stages of vein development Boxes illustrate positions of closeups in B and H, respectively, hv: minor vein; 11, 12 and 13: first, second and third loops; mv: midvein. (B–F,H–L) Confocal laser scanning microscopy. First leaves. Top right: genotype, treatment and leaf age in days after germination (DAG). Dashed white line delineates leaf outline. Bottom left: reproducibility index. PIN1::PIN1:GFP expression; look-up table (ramp in H) visualizes expression levels. Red: autofluorescence. Bars: (B–F,H–L) 20 μm.

Consistent with previous reports (Scarpella et al., 2006; Wenzel et al., 2007), PIN1::PIN1:GFP expression domains were broader at early stages of development of the tissue that in NPA-grown WT corresponds to that from which the second loop forms in WT; PIN1::PIN1:GFP expression was localized isotropically, or nearly so, at the PM in the outermost inner cells but was mainly localized to the basal side of the PM in the innermost inner cells (Fig. 13D). At later stages of second-loop development in NPA-grown WT, by which time PIN1::PIN1:GFP expression had become restricted to the sole, elongated cells of the developing loop, PIN1::PIN1:GFP expression was localized to the basal side of the PM (Fig. 13J).

As in NPA-grown WT, in both *gn^van7^* and NPA-grown *tir1;afb2* PIN1::PIN1:GFP expression domains were broader at early stages of development of the tissue that corresponds to that from which the second loop forms in WT, but PIN1::PIN1:GFP was expressed more heterogeneously in *gn^van7^* and NPA-grown *tir1;afb2* than in NPA-grown WT (Fig. 13E,F). Nevertheless, as in NPA-grown WT, in both *gn^van7^* and NPA-grown *tir1;afb2* PIN1::PIN1:GFP expression remained localized isotropically, or nearly so, at the PM, except in cells near the edge of higher-expression domains; in those cells, localization of PIN1::PIN1:GFP expression atthe PM was weakly polar, but such weak cell polarities pointed in seemingly random directions (Fig. 13E,F). At later stages of second-loop development of both *gn^van7^* and NPA-grown *tir1;afb2*, heterogeneity of PIN1::PIN1:GFP expression had become more pronounced, and PIN1::PIN1:GFP expression had become restricted to narrow clusters of cells; in those cells, localization of PIN1::PIN1:GFP expression at the PM was weakly polar, but such weak cell polarities still pointed in seemingly random directions (Fig. 13K,L).

In conclusion, simultaneous defects in auxin transport and signaling recapitulate *gn* defects in coordination of PIN1 polar localization, suggesting not only that the vein pattern defects of *gn* are caused by simultaneous defects in auxin transport and signaling, but that simultaneous defects in auxin transport and signaling recapitulate *gn* defects in coordination of tissue cell polarity during vein formation.

## Discussion

The current hypothesis of how auxin coordinates tissue cell polarity to induce polar-vein-formation proposes that GN controls the cellular localization of PIN1 and other PIN proteins; the resulting cell-to-cell, polar transport of auxin would coordinate tissue cell polarity and control polar developmental processes such as vein formation (reviewed in, e.g., (Berleth et al., 2000; Richter et al., 2010; Nakamura et al., 2012; Linh et al., 2018)).

Contrary to predictions of the hypothesis, we find that auxin-induced polar-vein-formation occurs in the absence of PIN proteins or any known intercellular auxin transporter, that the residual auxin-transport-independent vein-patterning activity relies on auxin signaling, and that a *GN*-dependent signal that coordinates tissue cell polarity to induce polar-vein-formation acts upstream of both auxin transport and signaling (Fig. S14).

### Control of Vein Patterning by Carrier-Mediated Polar Auxin-Transport

Overwhelming experimental evidence places polar auxin transport at the core of the mechanism that defines sites of vein formation (reviewed in (Sachs, 1981; Sachs, 1991a; Berleth et al., 2000; Sachs, 2000; Sawchuk and Scarpella, 2013)). The polarity of auxin transport is determined by the asymmetric localization of efflux carriers of the PIN family at the PM of auxin-transporting cells (Wisniewska et al., 2006). Therefore, loss of function of all the PM-PIN proteins should lead to loss of reproducible vein-pattern features or even, in the most extreme case, to the inability to form veins. Neither prediction is, however, supported by evidence: mutants in all the *PM-PIN* genes with vein patterning function — *PIN1, PIN3, PIN4* and *PIN7* — or in all the *PM-PIN* genes — *PIN1–PIN4* and *PIN7* — form veins, and these veins are arranged in reproducible, albeit abnormal, patterns. The most parsimonious account for the discrepancy between the observed and expected mutant defects is that vein patterning is controlled by additional, PM-PIN-independent auxin-transport pathways.

The existence of PM-PIN-independent auxin-transport pathways with vein patterning function can also be inferred from the discrepancy between the vein pattern defects of *pin1,3;4;7*, or of *pin1,3;2;4;7*, and those induced by NPA, which is thought to be a specific inhibitor of carrier-mediated cellular auxin-efflux (Cande and Ray, 1976; Sussman and Goldsmith, 1981; Petrasek et al., 2003; Dhonukshe et al., 2008). The vein pattern defects of WT grown in the presence of NPA are more severe than those of *pin1,3;4;7* or *pin1,3;2;4;7*, suggesting the existence of an NPA-sensitive auxin-transport pathway with vein patterning function in addition to that controlled by PM-PIN proteins, a suggestion that is supported by the observation that growth in the presence of NPA enhances the vein pattern defects of *pin1,3;4;7* to match those induced in WT by NPA.

Such PM-PIN-independent NPA-sensitive auxin-transport pathway with vein patterning function depends on the activity of the ER-PIN proteins PIN6 and PIN8, as inferred from the identity of the vein pattern defects induced in WT by NPA and those of *pin1,3,6;4;7;8*, and from the inability of NPA to induce further defects in *pin1,3,6;4;7;8*. Moreover, that NPA-grown WT phenocopies *pin1,3,6;4;7;8*, that no further defects can be induced in *pin1,3,6;4;7;8* by NPA, and that the vein patterns of *pin1,3,6;4;7;8* and NPA-grown WT fall into the same single phenotype-class suggest the absence of NPA-sensitive vein-patterning activity beyond that provided by PIN1, PIN3, PIN4, PIN6, PIN7 and PIN8, and hence the existence of NPA-insensitive vein-patterning pathways. It is of course possible that PIN6 and PIN8 are partially localized to the PM, and PM-localization of PIN5 and PIN6 has indeed been reported (Ganguly et al., 2014; Bennett et al., 2016; Simon et al., 2016; Ditengou et al., 2018); most important, however, that observation would not argue against the existence of NPA-insensitive vein patterning pathways, which is a logical conclusion, not a hypothesis.

These NPA-insensitive vein-patterning pathways are unlikely to be mediated by known intercellular auxin transporters — the AUX1/LAX influx carriers (Yang et al., 2006; Swarup et al., 2008; Peret et al., 2012) and the ABCB efflux carriers (Geisler et al., 2005; Bouchard et al., 2006; Petrasek et al., 2006) — as their mutation fails to enhance the vein pattern defects of *pin1,3,6* and of the NPA-induced phenocopy of *pin1,3,6;4;7;8*. Though it remains unexplored whether the NPA-insensitive vein-patterning pathways depend on the function of the PIN-LIKES intracellular auxin-transporters (Barbez et al., 2012), and though we cannot rule out the existence of unknown auxin transporters, it is unlikely that the NPA-insensitive vein-patterning pathways depend on NPA-insensitive carrier-mediated auxin-transport because as little as 10 μM NPA (a fraction of the concentration we used) is sufficient to inhibit carrier-mediated polar auxin-transport completely in tissue segments (Okada et al., 1991; Kaneda et al., 2011). Whatever the molecular nature of the NPA-insensitive vein-patterning pathways, they do contribute to the polar propagation of the inductive auxin signal: application of auxin to *pin1,3,6;4;7;8* leaves, just as to WT leaves, induces the formation of veins that connect the applied auxin to the pre-existing vasculature basal to the site of auxin application.

### Control of Vein Patterning by Auxin Signaling

The residual NPA-insensitive auxin-dependent vein-patterning activity of *pin1,3,6;4;7;8* relies, at least in part, on the signal transduction mediated by the TIR1/AFB auxin receptors and their post-translational regulator AXR1. Loss of *AXR1* or of *TIR1* and *AFB2*, the two auxin receptors that most contribute to auxin signaling (Dharmasiri et al., 2005), or growth in the presence of the auxin signaling inhibitor PBA (Matthes and Torres-Ruiz, 2016), induces entirely new vein-pattern defects in *pin1,3,6;4;7;8* or in its NPA-induced phenocopy, defects never observed in *pin1,3,6;4;7;8* or NPA-grown WT: in the more-severely affected leaves of *axr1;pin1,3,6;4;7;8, tir1;afb2;pin1,3,6;4;7;8*, NPA-grown *axr1*, NPA-grown *tir1;afb2* and NPA- and PBA-grown WT, the end-to-end alignment of vascular elements oriented with their axis along the axis of the vein is often replaced by the clustered differentiation of abnormally oriented vascular elements. Not only are these defects never observed in *pin1,3,6;4;7;8* or NPA-grown WT, but they are more severe than the predicted sum of the defects of *pin1,3,6;4;7;8* or NPA-grown WT, on the one hand, and of *axr1, tir1;afb2* or PBA-grown WT, on the other. These observations are particularly interesting because genetic analysis of auxin signaling components has so far implicated auxin signaling only in the differentiation of normally patterned veins (Przemeck et al., 1996; Hardtke and Berleth, 1998; Hardtke et al., 2004; Alonso-Peral et al., 2006; Candela et al., 2007; Esteve-Bruna et al., 2013). Instead, the mutual synthetic enhancement between the vein pattern defects caused by reduced auxin signaling and those caused by reduced auxin transport suggests non-homologous redundancy of auxin signaling and auxin transport in vein patterning, a conclusion which is consistent with observations in the shoot apical meristem (Schuetz et al., 2008). Unlike in the shoot apical meristem, however, in the leaf such redundancy is unequal: whereas auxin transport is required for vein patterning even in the presence of normal auxin signaling, the vein patterning activity of auxin signaling is only exposed in conditions of compromised auxin transport.

How auxin signaling, inherently non-directional (Leyser, 2018), could contribute to the polar propagation of the inductive auxin signal in the absence of carrier-mediated polar auxin-transport is unclear. One possibility is that auxin signaling promotes the passive diffusion of auxin through the tissue by controlling, for example, the proton gradient across the PM (Fendrych et al., 2016). However, it is difficult to conceive how auxin diffusion through a specific side of the PM would positively feed back on the ability of auxin to diffuse through that specific side of the PM — a positive feedback that would be required to drain neighboring cells from auxin and thus to form veins, i.e. channels of preferential auxin movement (Sachs, 1969).

One other possibility is that auxin signaling promotes the facilitated diffusion of auxin through the plasmodesmata intercellular channels, a possibility that had previously been suggested (Mitchison, 1980) and that has recently received some experimental support (Han et al., 2014). Here, how auxin movement through a specific side of the PM could positively feed back on the ability of the cell to move auxin through that specific side of the PM is conceivable (e.g., (Cieslak et al., 2015)), but no experimental evidence exists of such feedback or that auxin movement through plasmodesmata controls vein patterning.

Yet another possibility is that auxin signaling activates an unknown mobile signal. Such signal need not be chemical and alternatives, for example a mechanical signal, have been suggested (Couder et al., 2002; Laguna et al., 2008; Corson et al., 2009; Lee et al., 2014) and have been implicated in other auxin-driven processes (e.g., (Hamant et al., 2008; Heisler et al., 2010; Peaucelle et al., 2011; Nakayama et al., 2012; Braybrook and Peaucelle, 2013)). However, whether a mechanical signal controls vein patterning remains to be tested.

### A Tissue-Cell-Polarizing Signal Upstream of Auxin Transport and Signaling

The vein pattern defects of leaves in which both transport and transduction of the auxin signal are compromised are never observed in leaves in which either process is; yet those defects are not unprecedented: they are observed — though in more extreme form — in leaves of *gn* mutants, suggesting that *GN* controls both transport and transduction of the auxin signal during vein patterning.

That *GN* controls PM-PIN-mediated auxin transport during vein patterning is also suggested by the very limited or altogether missing restriction of PIN1 expression domains and coordination of PIN1 polar localization during *gn* leaf development, which is consistent with observations in embryos and roots (Steinmann et al., 1999; Kleine-Vehn et al., 2008). However, if failure to coordinate the polarization of the localization of PIN1 — and possibly other PM-PIN proteins — were the sole cause of the vein pattern defects of *gn*, these defects would be dependent on *PM-PIN*function and would therefore be masked by those of *pin1,3;4;7* in the *gn;pin1,3;4;7* mutant. The epistasis of the vein pattern defects of *gn* to those of *pin1,3;4;7* instead suggests that the vein pattern defects of *gn* are independent of *PM-PIN* function, and therefore that they are not the sole result of loss or abnormal polarity of PM-PIN-mediated auxin transport and that *GN*-acts upstream of *PM-PIN* genes in vein patterning. Moreover, the epistasis of the vein pattern defects of *gn* to those of *pin1,3,6;4;7;8* and the inability of NPA, which phenocopies the vein pattern defects of *pin1,3,6;4;7;8*, to induce additional defects in *gn* suggest that the vein pattern defects of *gn* are independent of all the *PIN* genes with vein patterning function, and therefore that those defects are not the sole result of loss or abnormal polarity of PIN-mediated auxin transport, and that *GN* acts upstream of all the *PIN* genes in vein patterning. Whereas mechanisms by which *GN* may control PM-PIN-mediated auxin transport have been suggested (e.g., (Richter et al., 2010; Luschnig and Vert, 2014; Naramoto et al., 2014)), it is unclear how *GN* could control auxin transport mediated by the ER-localized PIN6 and PIN8; it is possible, however, that such control is mediated by *GN* function in ER-Golgi trafficking (Richter et al., 2007; Teh and Moore, 2007; Nakano et al., 2009).

These observations suggest that the function of *GN* in coordination of tissue cell polarity and vein patterning entails more than the regulation of PIN-mediated auxin transport, a conclusion which is consistent with functions of *GN* that do not seem to be related to auxin transport or mediated by *PIN* genes (Shevell et al., 2000; Fischer et al., 2006; Irani et al., 2012; Nielsen et al., 2012; Moriwaki et al., 2014).

The auxin-transport-, *PIN*-independent functions of *GN* in coordination of tissue cell polarity and vein patterning are, at least in part, provided by TIR1/AFB2- and AXR1-mediated auxin signaling. This conclusion is suggested by the ability of simultaneous reduction in auxin transport and signaling to phenocopy defects in coordination of tissue cell polarity, auxin response and vein patterning of *gn*; it is also supported by the epistasis of the vein pattern defects of *gn* to those of *axr1*, which is consistent with genetic analysis placing *GN* upstream of auxin signaling in the formation of apical-basal polarity in the embryo (Mayer et al., 1993).

Though it is unclear how *GN* controls auxin signaling during vein patterning, the most parsimonious account is that *GN* controls the coordinated localization of proteins produced in response to auxin signaling. Auxin signaling has indeed been shown to control the production of proteins that are polarly localized at the plasma membrane of root cells (e.g., (Scacchi et al., 2009; Scacchi et al., 2010; Yoshida et al., 2019)), and at least some of these proteins act synergistically with PIN-mediated auxin transport in the root (e.g., (Marhava et al., 2018)); however, it remains to be tested whether such proteins have vein patterning activity, whether their localization is controlled by GN, and whether they mediate *GN* function in auxin signaling during vein patterning.

Alternatively, because cell wall composition and properties are abnormal in *gn* (Shevell et al., 2000), *GN* could control the production, propagation or interpretation of a mechanical signal that has been proposed to be upstream of both auxin signaling and transport in the shoot apical meristem (Heisler et al., 2010; Nakayama et al., 2012); however, whether a mechanical signal controls vein patterning and whether such signal acts downstream of *GN* remains to be tested.

Irrespective of the mechanism of action, our results reveal synergism between auxin transport and signaling, and their unsuspected control by *GN*, in the coordination of tissue cell polarity during vein patterning, a control whose logic is unprecedented in multicellular organisms.

## Materials & Methods

### Notation

In agreement with (Crittenden et al., 1996), linked genes or mutations (<2,500 kb apart, which in Arabidopsis corresponds, on average, to ~10 cM (Lukowitz et al., 2000)) are separated by a comma, unlinked ones by a semicolon and homologous chromosomes by a slash.

### Plants

Origin and nature of lines, genotyping strategies and oligonucleotide sequences are in Tables 1, 18 and 19. Seeds were sterilized and sown as in (Sawchuk et al., 2008). Stratified seeds were germinated and seedlings were grown at 22°C under continuous fluorescent light (~80 μmol m^-2^ s^-1^). Plants were grown at 25°C under fluorescent light (~110 μmol m^-2^ s^-1^) in a 16-h-light/8-h-dark cycle. Plants were transformed and representative lines were selected as in (Sawchuk et al., 2008).

### Chemicals

N-1-naphthylphthalamic acid and phenylboronic acid were dissolved in dimethyl sulfoxide and water, respectively; dissolved chemicals were added to growth medium just before sowing. Indole-3-acetic acid was dissolved in melted (55°C) lanolin; the IAA-lanolin paste was applied to first leaves 4 days after germination and was reapplied weekly.

### RT-PCR

Total RNA was extracted as in (Chomczynski and Sacchi, 1987) from 4-day-old seedlings grown as in (Odat et al., 2014). RT-PCR was performed as in (Odat et al., 2014) with the “GN_qFb” and “GN_qRb” oligonucleotides (Table S19), and with the “ROC1 F” and “ROC1 R” oligonucleotides (Beeckman et al., 2002) (Table S19).

### Imaging

Developing leaves were mounted and imaged as in (Sawchuk et al., 2013), except that emission was collected from ~2.5-μm-thick optical slices. Light paths are in Table S20.

Mature leaves were fixed in 3 : 1 or 6 : 1 ethanol: acetic acid, rehydrated in 70% ethanol and water, cleared briefly (few seconds to few minutes) — when necessary — in 0.4 M sodium hydroxide, washed in water, mounted in 80% glycerol or in 1 : 2 : 8 or 1 : 3 : 8 water: glycerol: chloral hydrate and imaged as in (Odat et al., 2014). Grayscaled RGB color images were turned into 8-bit images, look-up-tables were applied, and brightness and contrast were adjusted by linear stretching of the histogram in the Fiji distribution (Schindelin et al., 2012) of ImageJ (Schneider et al., 2012; Schindelin et al., 2015; Rueden et al., 2017).

## Acknowledgements

We thank the Arabidopsis Biological Resource Center for *emb30-8, gn-13, pin1-1, eir1-1, pin6, pin8-1, pgp1-100, mdr1-101, ucu2-4, aux1-355, lax1-064, axr1-3* and *axr1-12*; Hidehiro Fukaki for *fwr* and *gn^SALK_103014^*; Sandra Richter and Gerd Jürgens for *gn^B/E^* and *gn^R5^*; Satoshi Naramoto and Hiroo Fukuda for *van7/emb30-7*; Eva Benková and Jiří Friml for PIN1::PIN1:GFP; Jian Xu and Ben Scheres for PIN1::PIN1:YFP and PIN2::PIN2:GFP; Jian Xu, Miyo Morita and Masao Tasaka for PIN3::PIN3:GFP; Ikram Blilou and Ben Scheres for *Atpin1::En134, pin3-3, pin4-2* and *pin7^En^*; Venkatesan Sundaresan for *toz-1*, Thomas Berleth for *mp^G12^*; Markus Geisler and Jiří Friml for ABCB1::ABCB1:GFP and ABCB19::ABCB19:GFP; Cris Kuhlemeier for *aux1-21;lax1;2-1;3*; Michael Prigge and Mark Estelle for *tir1-1;afb2-3*; and Marcus Heisler and Elliot Meyerowitz for DR5rev::nYFP. This work was supported by Discovery Grants of the Natural Sciences and Engineering Research Council of Canada (NSERC) to E.S. C.V. was supported, in part, by a University of Alberta Doctoral Recruitment Scholarship. M.G.S was supported, in part, by an NSERC CGS-M Scholarship and an NSERC CGS-D Scholarship.

## References

Aloni, R. (2010). The induction of vascular tissues by auxin. In Plant hormones: biosynthesis, signal transduction, action (ed. D. PJ), pp. 485–506. Dordrecht: Kluwer Academic Publishers.

Alonso-Peral, M. M., Candela, H., del Pozo, J. C., Martinez-Laborda, A., Ponce, M. R. and Micol, J. L. (2006). The HVE/CAND1 gene is required for the early patterning of leaf venation in Arabidopsis. Development 133, 3755–3766.

Bailly, A., Sovero, V., Vincenzetti, V., Santelia, D., Bartnik, D., Koenig, B. W., Mancuso, S., Martinoia, E. and Geisler, M. (2008). Modulation of P-glycoproteins by auxin transport inhibitors is mediated by interaction with immunophilins. J. Biol. Chem. 283, 21817–21826.

Bainbridge, K., Guyomarc’h, S., Bayer, E., Swarup, R., Bennett, M., Mandel, T. and Kuhlemeier, C. (2008). Auxin influx carriers stabilize phyllotactic patterning. Genes Dev. 22, 810–823.

Barbez, E., Kubes, M., Rolcik, J., Beziat, C., Pencik, A., Wang, B., Rosquete, M. R., Zhu, J., Dobrev, P. I., Lee, Y. et al. (2012). A novel putative auxin carrier family regulates intracellular auxin homeostasis in plants. Nature 485, 119–122.

Bayer, E. M., Smith, R. S., Mandel, T., Nakayama, N., Sauer, M., Prusinkiewicz, P. and Kuhlemeier, C. (2009). Integration of transport-based models for phyllotaxis and midvein formation. Genes Dev. 23, 373–384.

Beeckman, T., Przemeck, G. K., Stamatiou, G., Lau, R., Terŗyn, N., De Rycke, R., Inze, D. and Berleth, T. (2002). Genetic complexity of cellulose synthase a gene function in Arabidopsis embryogenesis. Plant Physiol 130, 1883–1893.

Belteton, S. A., Sawchuk, M. G., Donohoe, B. S., Scarpella, E. and Szymanski, D. B. (2018). Reassessing the Roles of PIN Proteins and Anticlinal Microtubules during Pavement Cell Morphogenesis. Plant Physiol 176, 432–449.

Benkova, E., Michniewicz, M., Sauer, M., Teichmann, T., Seifertova, D., Jurgens, G. and Friml, J. (2003). Local, efflux-dependent auxin gradients as a common module for plant organ formation. cell 115, 591–602.

Bennett, T., Hines, G., van Rongen, M., Waldie, T., Sawchuk, M. G., Scarpella, E., Ljung, K. and Leyser, O. (2016). Connective Auxin Transport in the Shoot Facilitates Communication between Shoot Apices. PLoS Biol. 14, e1002446.

Berleth, T., Mattsson, J. and Hardtke, C. S. (2000). Vascular continuity and auxin signals. Trends Plant Sci 5, 387–393.

Blakeslee, J. J., Bandyopadhyay, A., Lee, O. R., Mravec, J., Titapiwatanakun, B., Sauer, M., Makam, S. N., Cheng, Y., Bouchard, R., Adamec, J. et al. (2007). Interactions among PIN-FORMED and P-glycoprotein auxin transporters in Arabidopsis. Plant Cell 19, 131–147.

Bosco, C. D., Dovzhenko, A., Liu, X., Woerner, N., Rensch, T., Eismann, M., Eimer, S., Hegermann, J., Paponov, I. A., Ruperti, B. et al. (2012). The endoplasmic reticulum localized PIN8 is a pollen specific auxin carrier involved in intracellular auxin homeostasis. Plant J 71 860-870.

Bouchard, R., Bailly, A., Blakeslee, J. J., Oehring, S. C., Vincenzetti, V., Lee, O. R., Paponov, I., Palme, K., Mancuso, S., Murphy, A. S. et al. (2006). Immunophilin-like TWISTED DWARF1 modulates auxin efflux activities of Arabidopsis P-glycoproteins. J. Biol. Chem. 281, 30603–30612.

Boutte, Y., Ikeda, Y. and Grebe, M. (2007). Mechanisms of auxin-dependent cell and tissue polarity. Curr Opin Plant Biol 10, 616–623.

Braybrook, S. A. and Peaucelle, A. (2013). Mechano-chemical aspects of organ formation in Arabidopsis thaliana: the relationship between auxin and pectin. PLoS One 8, e57813.

Calderon-Villalobos, L. I., Tan, X., Zheng, N. and Estelle, M. (2010). Auxin perception-structural insights. Cold Spring Harb Perspect Biol 2, a005546.

Cande, W. Z. and Ray, P. I. M. (1976). Nature of cell-to-cell transfer of auxin in polar transport. Planta 129, 43–52.

Candela, H., Alonso-Peral, M. M., Ponce, M. R. and Micol, J. L. (2007). Role of HEMIVENATA and the Ubiquitin Pathway in Venation Pattern Formation. Plant Signal Behav 2, 258–259.

Candela, H., Martinez-Laborda, A. and Micol, J. L. (1999). Venation pattern formation in Arabidopsis thaliana vegetative leaves. Dev. Biol. 205, 205–216.

Chen, R., Hilson, P., Sedbrook, J., Rosen, E., Caspar, T. and Masson, P. H. (1998). The arabidopsis thaliana AGRAVITROPIC 1 gene encodes a component of the polar-auxin-transport efflux carrier. Proc. Natl. Acad. Sci. U S A 95, 15112–15117.

Chomczynski, P. and Sacchi, N. (1987). Single-step method of RNA isolation by acid guanidinium thiocyanate-phenol-chloroform extraction. Anal. Biochem. 162, 156–159.

Cieslak, M., Runions, A. and Prusinkiewicz, P. (2015). Auxin-driven patterning with unidirectional fluxes. J Exp Bot 66, 5083–5102.

Corson, F., Adda-Bedia, M. and Boudaoud, A. (2009). In silico leaf venation networks: growth and reorganization driven by mechanical forces. J Theor Biol 259, 440–448.

Couder, Y., Pauchard, L., Allain, C. and Adda-Bedia, M. and Douady, S. (2002). The leaf venation as formed in a tensorial field. The European Physical Journal B 28, 135-138.

Crittenden, L. B. and Bitgood, J. J. Burt, D. W., Ponce de Leon, F. A. and Tixier-Boichard, M. (1996). Nomenclature for naming loci, alleles, linkage groups and chromosomes to be used in poultry genome publications and databases. Genet Sel Evol 28, 289-297.

Dengler, N. G. (2006). The shoot apical meristem and development of vascular architecture. Canadian Journal of Botany 84, 1660–1671.

Dharmasiri, N., Dharmasiri, S., Weijers, D., Lechner, E., Yamada, M., Hobbie, L., Ehrismann, J. S., Jurgens, G. and Estelle, M. (2005). Plant Development Is Regulated by a Family of Auxin Receptor F Box Proteins. Dev. Cell 9, 109–119.

Dhonukshe, P., Grigoriev, I., Fischer, R., Tominaga, M., Robinson, D. G., Hasek, J., Paciorek, T., Petrasek, J., Seifertova, D., Tejos, R. et al. (2008). Auxin transport inhibitors impair vesicle motility and actin cytoskeleton dynamics in diverse eukaryotes. Proc. Natl. Acad. Sci. U S A 105, 4489–4494.

Ding, Z., Wang, B., Moreno, I., Duplakova, N., Simon, S., Carraro, N., Reemmer, J., Pencik, A., Chen, X., Tejos, R. et al. (2012). ER-localized auxin transporter PIN8 regulates auxin homeostasis and male gametophyte development in Arabidopsis. Nat Commun 3, 941.

Ditengou, F. A., Gomes, D., Nziengui, H., Kochersperger, P., Lasok, H., Medeiros, V., Paponov, I. A., Nagy, S. K., Nádai, T. V., Mészáros, T. et al. (2018). Characterization of auxin transporter PIN6 plasma membrane targeting reveals a function for PIN6 in plant bolting. New Phytol 217, 1610–1624.

Esau, K. (1942). Vascular Differentiation in the Vegetative Shoot of Linum. I. The Procambium. American Journal of Botany 29, 738–747.

Esteve-Bruna, D., Perez-Perez, J. M., Ponce, M. R. and Micol, J. L. (2013). incurvata13, a novel allele of AUXIN RESISTANT6, reveals a specific role for auxin and the SCF complex in Arabidopsis embryogenesis, vascular specification, and leaf flatness. Plant Physiol 161, 1303–1320.

Fendrych, M., Leung, J. and Friml, J. (2016). TIR1/AFB-Aux/IAA auxin perception mediates rapid cell wall acidification and growth of Arabidopsis hypocotyls. Elife 5, e19048.

Fischer, U., Ikeda, Y., Ljung, K., Serralbo, O., Singh, M., Heidstra, R., Palme, K., Scheres, B. and Grebe, M. (2006). Vectorial information for Arabidopsis planar polarity is mediated by combined AUX1, EIN2, and GNOM activity. Curr. Biol. 16, 2143–2149.

Franzmann, L., Patton, D. A. and Meinke, D. W. (1989). Invitro Morphogenesis of Arrested Embryos from Lethal Mutants of Arabidopsis-Thaliana. Theoretical and Applied Genetics 77, 609–616.

Friml, J., Benkova, E., Blilou, I., Wisniewska, J., Hamann, T., Ljung, K., Woody, S., Sandberg, G., Scheres, B., Jurgens, G. et al. (2002a). AtPIN4 mediates sink-driven auxin gradients and root patterning in Arabidopsis. Cell 108, 661–673.

Friml, J., Vieten, A., Sauer, M., Weijers, D., Schwarz, H., Hamann, T., Offringa, R. and Jurgens, G. (2003). Efflux-dependent auxin gradients establish the apical-basal axis of Arabidopsis. Nature 426, 147–153.

Friml, J., Wisniewska, J., Benkova, E., Mendgen, K. and Palme, K. (2002b). Lateral relocation of auxin efflux regulator PIN3 mediates tropism in Arabidopsis. Nature 415, 806–809.

Galweiler, L., Guan, C., Muller, A., Wisman, E, Mendgen, K., Yephremov, A. and Palme, K. (1998). Regulation of polar auxin transport by AtPIN1 in Arabidopsis vascular tissue. Science 282, 2226–2230.

Ganguly, A., Park, M., Kesawat, M. S. and Cho, H.-T. (2014). Functional analysis of the hydrophilic loop in intracellular trafficking of Arabidopsis PIN-FORMED proteins. The Plant Cell 26, 1570–1585.

Geisler, M., Blakeslee, J. J., Bouchard, R., Lee, O. R., Vincenzetti, V., Bandyopadhyay, A., Titapiwatanakun, B., Peer, W. A., Bailly, A., Richards, E. L. et al. (2005). Cellular efflux of auxin catalyzed by the Arabidopsis MDR/PGP transporter AtPGP1. Plant J 44, 179–194.

Geisler, M., Kolukisaoglu, H. U., Bouchard, R., Billion, K., Berger, J., Saal, B., Frangne, N., Koncz-Kalman, Z., Koncz, C., Dudler, R. et al. (2003). TWISTED DWARF1, a unique plasma membrane-anchored immunophilin-like protein, interacts with Arabidopsis multidrug resistance-like transporters AtPGP1 and AtPGP19. Mol. Biol. Cell 14, 4238–4249.

Geisler, M. and Murphy, A. S. (2006). The ABC of auxin transport: the role of p-glycoproteins in plant development. FEBS Lett. 580, 1094–1102.

Geldner, N., Richter, S., Vieten, A., Marquardt, S., Torres-Ruiz, R. A., Mayer, U. and Jurgens, G. (2004). Partial loss-of-function alleles reveal a role for GNOM in auxin transport-related, post-embryonic development of Arabidopsis. Development, 131 389–400.

Goodrich, L. V. and Strutt, D. (2011). Principles of planar polarity in animal development. Development 138, 1877–1892.

Hamant, O., Heisler, M. G., Jonsson, H., Krupinski, P., Uyttewaal, M., Bokov, P., Corson, F., Sahlin, P., Boudaoud, A., Meyerowitz, E. M. et al. (2008). Developmental patterning by mechanical signals in Arabidopsis. Science 322, 1650–1655.

Han, X., Hyun, T. K., Zhang, M., Kumar, R., Koh, E. J., Kang, B. H., Lucas, W. J. and Kim, J. Y. (2014). Auxin-callose-mediated plasmodesmal gating is essential for tropic auxin gradient formation and signaling. Dev. Cell 28, 132–146.

Hardtke, C. S. and Berleth, T. (1998). The Arabidopsis gene MONOPTEROS encodes a transcription factor mediating embryo axis formation and vascular development. EMBO J. 17, 1405–1411.

Hardtke, C. S., Ckurshumova, W., Vidaurre, D. P., Singh, S. A., Stamatiou, G., Tiwari, S. B., Hagen, G., Guilfoyle, T. J. and Berleth, T. (2004). Overlapping and non-redundant functions of the Arabidopsis auxin response factors MONOPTEROS and NONPHOTOTROPIC HYPOCOTYL 4. Development 131, 1089–1100.

Heisler, M. G., Hamant, O., Krupinski, P., Uyttewaal, M., Ohno, C., Jonsson, H., Traas, J. and Meyerowitz, E. M. (2010). Alignment between PIN1 polarity and microtubule orientation in the shoot apical meristem reveals a tight coupling between morphogenesis and auxin transport. PLoS Biol 8, e1000516.

Heisler, M. G., Ohno, C., Das, P., Sieber, P., Reddy, G. V., Long, J. A. and Meyerowitz, E. M. (2005). Patterns of Auxin Transport and Gene Expression during Primordium Development Revealed by Live Imaging of the Arabidopsis Inflorescence Meristem. Curr. Biol 15, 1899–1911.

Irani, N. G., Di Rubbo, S., Mylle, E., Van den Begin, J., Schneider-Pizoń, J., Hniliková, J., Šíša, M., Buyst, D., Vilarrasa-Blasi, J. and Szatmári, A.-M. (2012). Fluorescent castasterone reveals BRI1 signaling from the plasma membrane. Nature chemical biology 8, 583.

Kamphausen, T., Fanghanel, J., Neumann, D., Schulz, B. and Rahfeld, J. U. (2002). Characterization of Arabidopsis thaliana AtFKBP42 that is membrane-bound and interacts with Hsp90. Plant 32, 263–276.

Kaneda, M., Schuetz, M., Lin, B. S., Chanis, C., Hamberger, B., Western, T. L., Ehlting, J. and Samuels, A. L. (2011). ABC transporters coordinately expressed during lignification of Arabidopsis stems include a set of ABCBs associated with auxin transport.J Exp Bot 62, 2063–2077.

Kang, J. and Dengler, N. (2004). Vein pattern development in adult leaves of Arabidopsis thaliana. International Journal of Plant Sciences 165, 231–242.

Katekar, G. F. and Geissler, A. E. (1980). Auxin Transport Inhibitors: IV. EVIDENCE OF A COMMON MODE OF ACTION FOR A PROPOSED CLASS OF AUXIN TRANSPORT INHIBITORS: THE PHYTOTROPINS. Plant Physiol 66, 1190–1195.

Kierzkowski, D., Lenhard, M., Smith, R. and Kuhlemeier, C. (2013). Interaction between meristem tissue layers controls phyllotaxis. Dev. Cell 26, 616–628.

Kinsman, E. A. and Pyke, K. A. (1998). Bundle sheath cells and cell-specific plastid development in Arabidopsis leaves. Development 125, 1815–1822.

Kleine-Vehn, J., Dhonukshe, P., Sauer, M., Brewer, P. B., Wisniewska, J., Paciorek, T., Benkova, E. and Friml, J. (2008). ARF GEF-dependent transcytosis and polar delivery of PIN auxin carriers in Arabidopsis. Curr. Biol. 18, 526–531.

Koizumi, K., Sugiyama, M. and Fukuda, H. (2000). A series of novel mutants of Arabidopsis thaliana that are defective in the formation of continuous vascular network: calling the auxin signal flow canalization hypothesis into question. Development 127, 3197–3204.

Krecek, P., Skupa, P., Libus, J., Naramoto, S., Tejos, R., Friml, J. and Zazimalova, E. (2009). The PIN-FORMED (PIN) protein family of auxin transporters. Genome Biol 10, 249.

Laguna, M. F., Bohn, S. and Jagla, E. A. (2008). The role of elastic stresses on leaf venation morphogenesis. PLoS Comput. Biol.

Lee, S. W., Feugier, F. G. and Morishita, Y. (2014). Canalization-based vein formation in a growing leaf. Journal of theoretical biology 353, 104-120.

Leyser, O. (2018). Auxin Signaling. Plant Physiol 176, 465–479.

Linh, N. M., Verna, C. and Scarpella, E. (2018). Coordination of cell polarity and the patterning of leaf vein networks. Curr Opin Plant Biol 41, 116–124.

Lukowitz, W., Gillmor, C. S. and Scheible, W. R. (2000). Positional cloning in Arabidopsis. Why it feels good to have a genome initiative working for you. Plant Physiol 123, 795–805.

Luschnig, C., Gaxiola, R. A., Grisafi, P. and Fink, G. R. (1998). EIR1, a root-specific protein involved in auxin transport, is required for gravitropism in Arabidopsis thaliana. Genes & Development 12, 2175–2187.

Luschnig, C. and Vert, G. (2014). The dynamics of plant plasma membrane proteins: PINs and beyond. Development 141, 2924–2938.

Marcos, D. and Berleth, T. (2014). Dynamic auxin transport patterns preceding vein formation revealed by live-imaging of Arabidopsis leaf primordia. Front Plant Sci 5, 235.

Marhava, P., Bassukas, A. E. L., Zourelidou, M., Kolb, M., Moret, B., Fastner, A., Schulze, W. X., Cattaneo, P., Hammes, U. Z., Schwechheimer, C. et al. (2018). A molecular rheostat adjusts auxin flux to promote root protophloem differentiation. Nature 558, 297–300.

Matthes, M. and Torres-Ruiz, R. A. (2016). Boronic acids induce monopteros phenocopies by affecting PIN1 membrane stability and polar auxin transport in Arabidopsis thaliana embryos. Development 143, 4053-4062.

Mattsson, J., Sung, Z. R. and Berleth, T. (1999). Responses of plant vascular systems to auxin transport inhibition. Development 126, 2979–2991.

Mayer, U., Buttner, G. and Jurgens, G. (1993). Apical-basal pattern formation in the Arabidopsis embryo: studies on the role of the gnom gene. Development 117, 149–162.

Mitchison, G. J. (1980). Model for Vein Formation in Higher-Plants. Proceedings of the Royal Society of London Series B-Biological Sciences 207, 79–109.

Moriwaki, T., Miyazawa, Y., Fujii, N. and Takahashi, H. (2014). GNOM regulates root hydrotropism and phototropism independently of PIN-mediated auxin transport. Plant Sci 215–216, 141–149.

Mravec, J., Kubes, M., Bielach, A., Gaykova, V., Petrasek, J., Skupa, P., Chand, S., Benkova, E., Zazimalova, E. and Friml, J. (2008). Interaction of PIN and PGP transport mechanisms in auxin distribution-dependent development. Development 135, 3345–3354.

Mravec, J., Skupa, P., Bailly, A., Hoyerova, K., Krecek, P., Bielach, A., Petrasek, J., Zhang, J., Gaykova, V., Stierhof, Y. D. et al. (2009). Subcellular homeostasis of phytohormone auxin is mediated by the ER-localized P1N5 transporter. Nature 459, 1136–1140.

Muller, A., Guan, C., Galweiler, L., Tanzler, P., Huijser, P., Marchant, A., Parry, G., Bennett, M., Wisman, E. and Palme, K. (1998). AtPIN2 defines a locus of Arabidopsis for root gravitropism control. EMBO J. 17, 6903–6911.

Nakamura, M., Kiefer, C. S. and Grebe, M. (2012). Planar polarity, tissue polarity and planar morphogenesis in plants. Curr Opin Plant Biol 15, 593–600.

Nakano, R. T., Matsushima, R., Ueda, H., Tamura, K., Shimada, T., Li, L., Hayashi, Y., Kondo, M., Nishimura, M. and Hara-Nishimura, I. (2009). GNOM-LIKE1/ERMO1 and SEC24a/ERMO2 are required for maintenance of endoplasmic reticulum morphology in Arabidopsis thaliana. Plant Cell 21, 3672–3685.

Nakayama, N., Smith, R. S., Mandel, T., Robinson, S., Kimura, S., Boudaoud, A. and Kuhlemeier, C. (2012). Mechanical regulation of auxin-mediated growth. Curr. Biol. 22, 1468–1476.

Naramoto, S., Otegui, M. S., Kutsuna, N., de Rycke, R., Dainobu, T., Karampelias, M., Fujimoto, M., Feraru, E., Miki, D., Fukuda, H. et al. (2014). Insights into the localization and function of the membrane trafficking regulator GNOM ARF-GEF at the Golgi apparatus in Arabidopsis. Plant Cell. l26, 3062–3076.

Nelson, T. and Dengler, N. (1997). Leaf Vascular Pattern Formation. Plant Cell 9, 1121–1135.

Nielsen, M. E., Feechan, A., Böhlenius, H., Ueda, T. and Thordal-Christensen, H. (2012). Arabidopsis ARF-GTP exchange factor, GNOM, mediates transport required for innate immunity and focal accumulation of syntaxin PEN1. Proceedings of the National Academy of Sciences 109, 11443–11448.

Odat, O., Gardiner, J., Sawchuk, M. G., Verna, C., Donner, T. J. and Scarpella, E. (2014). Characterization of an allelic series in the MONOPTEROS gene of Arabidopsis. Genesis 52, 127–133.

Okada, K., Ueda, J., Komaki, M. K., Bell, C. J. and Shimura, Y. (1991). Requirement of the Auxin Polar Transport System in Early Stages of Arabidopsis Floral Bud Formation. Plant Cell 3, 677–684.

Okumura, K., Goh, T., Toyokura, K., Kasahara, H., Takebayashi, Y., Mimura, T., Kamiya, Y. and Fukaki, H. (2013). GNOM/FEWER ROOTS is required for the establishment of an auxin response maximum for arabidopsis lateral root initiation. Plant Cell Physiol 54, 406–417.

Paponov, I. A., Teale, W. D., Trebar, M., Blilou, I. and Palme, K. (2005). The PIN auxin efflux facilitators: evolutionary and functional perspectives. Trends Plant Sci 10, 170–177.

Parry, G., Marchant, A., May, S., Swarup, R., Swarup, K., James, N., Graham, N., Allen, T., Martucci, T., Yemm, A. et al. (2001). Quick on the uptake: Characterization of a family of plant auxin influx carriers. Journal of Plant Growth Regulation 20, 217–225.

Peaucelle, A., Braybrook, S. A., Le Guillou, L., Bron, E., Kuhlemeier, C. and Höfte, H. (2011). Pectin-induced changes in cell wall mechanics underlie organ initiation in Arabidopsis. Curr. Biol. 21, 1720–1726.

Peret, B., Swarup, K., Ferguson, A., Seth, M., Yang, Y. D., Dhondt, S., James, N., Casimiro, I., Perry, P., Syed, A. et al. (2012). AUX/LAX Genes Encode a Family of Auxin Influx Transporters That Perform Distinct Functions during Arabidopsis Development. Plant Cell 24, 2874–2885.

Perez-Perez, J. M., Ponce, M. R. and Micol, J. L. (2004). The ULTRACURVATA2 gene of Arabidopsis encodes an FK506-binding protein involved in auxin and brassinosteroid signaling. Plant Physiol 134, 101–117.

Petrasek, J., Cerna, A., Schwarzerova, K., Elckner, M., Morris, D. A. and Zazimalova, E. (2003). Do phytotropins inhibit auxin efflux by impairing vesicle traffic? Plant Physiol 131, 254–263.

Petrasek, J., Mravec, J., Bouchard, R., Blakeslee, J. J., Abas, M., Seifertova, D., Wisniewska, J., Tadele, Z., Kubes, M., Covanova, M. et al. (2006). PIN proteins perform a rate-limiting function in cellular auxin efflux. Science 312, 914–918.

Przemeck, G. K., Mattsson, J., Hardtke, C. S., Sung, Z. R. and Berleth, T. (1996). Studies on the role of the Arabidopsis gene MONOPTEROS in vascular development and plant cell axialization. Planta 200, 229–237.

Raven, J. A. (1975). Transport of indole acetic acid in plant cells in relation to pH and electrical potential gradients, and its significance for polar IAA transport. New Phytologist 74, 163–172.

Reinhardt, D., Pesce, E. R., Stieger, P., Mandel, T., Baltensperger, K., Bennett, M., Traas, J., Friml, J. and Kuhlemeier, C. (2003). Regulation of phyllotaxis by polar auxin transport. Nature 426, 255–260.

Richter, S., Anders, N., Wolters, H., Beckmann, H., Thomann, A., Heinrich, R., Schrader, J., Singh, M. K., Geldner, N., Mayer, U. et al. (2010). Role of the GNOM gene in Arabidopsis apical-basal patterning-From mutant phenotype to cellular mechanism of protein action. Eur. J. Cell Biol. 89, 138–144.

Richter, S., Geldner, N., Schrader, J., Wolters, H., Stierhof, Y. D., Rios, G., Koncz, C., Robinson, D. G. and Jürgens, G. (2007). Functional diversification of closely related ARF-GEFs in protein secretion and recycling. Nature 448, 488–492.

Rubery, P. H. and Sheldrake, A. R. (1974). Carrier-Mediated Auxin Transport. Planta 118, 101–121.

Rueden, C. T., Schindelin, J., Hiner, M. C., DeZonia, B. E., Walter, A. E., Arena, E. T. and Eliceiri, K. W. (2017). ImageJ2:ImageJ for the next generation of scientific image data. BMC Bioinformatics 18, 529.

Sachs, T. (1969). Polarity and the Induction of Organized Vascular Tissues. Annals of Botany 33, 263–275.

Sachs, T. (1975). Control of Differentiation of Vascular Networks. Annals of Botany 39, 197–204.

Sachs, T. (1981). The control of the patterned differentiation of vascular tissues. Advances in Botanical Research 9, 151–262.

Sachs, T. (1991a). Cell Polarity and Tissue Patterning in Plants. Development Suppl 1, 83–93.

Sachs, T. (1991b). Pattern Formation in Plant Tissues. Cambridge: Cambridge University Press.

Sachs, T. (2000). Integrating Cellular and Organismic Aspects of Vascular Differentiation. Plant and Cell Physiology 41, 649–656.

Sawchuk, M. G., Donner, T. J., Head, P. and Scarpella, E. (2008). Unique and overlapping expression patterns among members of photosynthesis-associated nuclear gene families in Arabidopsis. Plant Physiol 148, 1908–1924.

Sawchuk, M. G., Edgar, A. and Scarpella, E. (2013). Patterning of leaf vein networks by convergent auxin transport pathways. PLoS Genet, 9, e1003294.

Sawchuk, M. G., Head, P., Donner, T. J. and Scarpella, E. (2007). Time-lapse imaging of Arabidopsis leaf development shows dynamic patterns of procambium formation. New Phytol 176, 560–571.

Sawchuk, M. G. and Scarpella, E. (2013). Polarity, continuity, and alignment in plant vascular strands. J Integr Plant Biol 55, 824–834.

Scacchi, E., Osmont, K. S., Beuchat, J., Salinas, P., Navarrete-Gómez, M., Trigueros, M., Ferrándiz, C. and Hardtke, C. S. (2009). Dynamic, auxin-responsive plasma membrane-to-nucleus movement of Arabidopsis BRX. Development 136, 2059–2067.

Scacchi, E., Salinas, P., Gujas, B., Santuari, L., Krogan, N., Ragni, L., Berleth, T. and Hardtke, C. S. (2010). Spatio-temporal sequence of cross-regulatory events in root meristem growth. Proc. Natl. Acad. Sci. U S A 107, 22734–22739.

Scarpella, E., Francis, P. and Berleth, T. (2004). Stage-specific markers define early steps of procambium development in Arabidopsis leaves and correlate termination of vein formation with mesophyll differentiation. Development 131, 3445–3455.

Scarpella, E., Marcos, D., Friml, J. and Berleth, T. (2006). Control of leaf vascular patterning by polar auxin transport. Genes Dev. 20, 1015–1027.

Schindelin, J., Arganda-Carreras, I., Frise, E., Kaynig, V., Longair, M., Pietzsch, T., Preibisch, S., Rueden, C., Saalfeld, S., Schmid, B. et al. (2012). Fiji: an open-source platform for biological-image analysis. Nat Methods 9, 676–682.

Schindelin, J., Rueden, C. T., Hiner, M. C. and Eliceiri, K. W. (2015). The ImageJ ecosystem: An open platform for biomedical image analysis. Mol. Reprod. Dev. 82, 518–529.

Schneider, C. A., Rasband, W. S. and Eliceiri, K. W. (2012). NIH Image to ImageJ: 25 years of image analysis. Nat Methods 9, 671–675.

Schuetz, M., Berleth, T. and Mattsson, J. (2008). Multiple MONOPTEROS-dependent pathways are involved in leaf initiation. Plant Physiol 148, 870–880.

Schwechheimer, C. (2018). NEDD8-its role in the regulation of Cullin-RING ligases. Curr Opin Plant Biol 45, 112–119.

Shevell, D. E., Kunkel, T. and Chua, N. H. (2000). Cell wall alterations in the arabidopsis emb30 mutant. Plant Cell 12, 2047–2060.

Shevell, D. E., Leu, W. M., Gillmor, C. S., Xia, G., Feldmann, K. A. and Chua, N. H. (1994). EMB30 is essential for normal cell division, cell expansion, and cell adhesion in Arabidopsis and encodes a protein that has similarity to Sec7. Cell 77, 1051–1062.

Sieburth, L. E. (1999). Auxin is required for leaf vein pattern in Arabidopsis. Plant Physiol 121, 1179–1190.

Simon, S., Skůpa, P., Viaene, T., Zwiewka, M., Tejos, R., Klíma, P., Čarná, M., Rolčík, J., De Rycke, R. and Moreno, I. (2016). PIN6 auxin transporter at endoplasmic reticulum and plasma membrane mediates auxin homeostasis and organogenesis in Arabidopsis. New Phytologist 211, 65–74.

Steinmann, T., Geldner, N., Grebe, M., Mangold, S., Jackson, C. L., Paris, S., Galweiler, L., Palme, K. and Jurgens, G. (1999). Coordinated polar localization of auxin efflux carrier PIN1 by GNOM ARF GEF. Science 286, 316–318.

Steynen, Q. J. and Schultz, E. A. (2003). The FORKED genes are essential for distal vein meeting in Arabidopsis. Development 130, 4695–4708.

Sussman, M. R. and Goldsmith, M. H. M. (1981). Auxin uptake and action of N-1-naphthylphthalamic acid in corn coleoptiles. Planta 151, 15–25.

Swarup, K., Benkova, E., Swarup, R., Casimiro, I., Peret, B., Yang, Y., Parry, G., Nielsen, E., De Smet, I., Vanneste, S. et al. (2008). The auxin influx carrier LAX3 promotes lateral root emergence. Nat. Cell Biol. 10, 946–954.

Teh, O.-k. and Moore, I. (2007). An ARF-GEF acting at the Golgi and in selective endocytosis in polarized plant cells. Nature 448, 493.

Telfer, A. and Poethig, R. S. (1994). Leaf development in Arabidopsis. In Arabidopsis (ed. E. M. Meyerowitz and C. R. Somerville), pp. 379–401. New York: Cold Spring Harbor Press.

Verna, C., Sawchuk, M. G., Linh, N. M. and Scarpella, E. (2015). Control of vein network topology by auxin transport. BMC Biol 13, 94.

Viaene, T., Delwiche, C. F., Rensing, S. A. and Friml, J. (2012). Origin and evolution of PIN auxin transporters in the green lineage. Trends Plant Sci 18, 5-10.

Wang, B., Bailly, A., Zwiewka, M., Henrichs, S., Azzarello, E., Mancuso, S., Maeshima, M., Friml, J., Schulz, A. and Geisler, M. (2013). Arabidopsis TWISTED DWARF1 functionally interacts with auxin exporter ABCB1 on the root plasma membrane. Plant Cell 25, 202–214.

Wenzel, C. L., Schuetz, M., Yu, Q. and Mattsson, J. (2007). Dynamics of MONOPTEROS and P1N-FORMED1 expression during leaf vein pattern formation in Arabidopsis thaliana. Plant J 49, 387–398.

Wisniewska, J., Xu, J., Seifertova, D., Brewer, P. B., Ruzicka, K., Blilou, I., Rouquie, D., Benkova, E., Scheres, B. and Friml, J. (2006). Polar PIN localization directs auxin flow in plants. Science 312, 883.

Wu, G., Otegui, M. S. and Spalding, E. P. (2010). The ER-localized TWD1 immunophilin is necessary for localization of multidrug resistance-like proteins required for polar auxin transport in Arabidopsis roots. Plant Cell 22, 3295–3304.

Xu, J., Hofhuis, H., Heidstra, R., Sauer, M., Friml, J. and Scheres, B. (2006). A molecular framework for plant regeneration. Science 311, 385–388.

Xu, J. and Scheres, B. (2005). Dissection of Arabidopsis ADP-R1BOSYLAT1ON FACTOR 1 function in epidermal cell polarity. Plant Cell 17, 525–536.

Yang, H. and Murphy, A. S. (2009). Functional expression and characterization of Arabidopsis ABCB, AUX 1 and PIN auxin transporters in Schizosaccharomyces pombe. Plant J 59, 179–191.

Yang, Y., Hammes, U. Z., Taylor, C. G., Schachtman, D. P. and Nielsen, E. (2006). High-affinity auxin transport by the AUX1 influx carrier protein. Curr. Biol. 16, 1123–1127.

Yoshida, S., van der Schuren, A., van Dop, M., van Galen, L., Saiga, S., Adibi, M., Möller, B., ten Hove, C. A., Marhavy, P., Smith, R. et al. (2019). A SOSEKI-based coordinate system interprets global polarity cues in Arabidopsis. Nature Plants 5, 160–166.

Zadnikova, P., Petrasek, J., Marhavy, P., Raz, V., Vandenbussche, F., Ding, Z., Schwarzerova, K., Morita, M. T., Tasaka, M., Hejatko, J. et al. (2010). Role of PIN-mediated auxin efflux in apical hook development of Arabidopsis thaliana. Development 137, 607–617.

Zourelidou, M., Absmanner, B., Weller, B., Barbosa, I. C., Willige, B. C., Fastner, A., Streit, V., Port, S. A., Colcombet, J., de la Fuente van Bentem, S. et al. (2014). Auxin efflux by PIN-FORMED proteins is activated by two different protein kinases, D6 PROTEIN KINASE and PINOID. Elife 3, e02860.

