## Supplemental Table 1 for "Coordination of Tissue Cell Polarity by Auxin Transport and Signaling"

**Table S1. Origin and nature of lines**

| **Line** | **Origin/Nature** |
| --- | --- |
| PIN1::PIN1:YFP | (Xu et al., 2006) |
| *gn-13* | ABRC; SALK_045424 (Alonso et al., 2003); contains a T-DNA insertion after +2835 of *GN* (AT1G13980) |
| PIN1::PIN1:GFP | (Benkova et al., 2003) |
| *gn-18* | ABRC; SALK_026031; contains a T-DNA insertion after -1047 of *GN* (AT1G13980) |
| *fwr* (*gn^fwr^*) | (Okumura et al., 2013) |
| *gn^B/E^* | (Geldner et al., 2004) |
| *gn^R5^* | (Geldner et al., 2004) |
| *van7*/*emb30-7* (*gn^van7^*) | (Koizumi et al., 2000) |
| *gn^van7+fwr^* | *gn^van7^* (-2127 to +5388; primers: ‘GN Fwd NotI’ and ‘GN Rev NotI’) containing the *fwr* mutation (primers: ‘fwr-mutagenesis F’ and ‘fwr-mutagenesis R’) |
| *gn^SALK_103014^* | ABRC; (Okumura et al., 2013) |
| *emb30-8* (*gn^emb30-8^*) | ABRC; (Franzmann et al., 1989; Moriwaki et al., 2013) |
| PIN2::PIN2:GFP | (Xu and Scheres, 2005) |
| PIN3::PIN3:GFP | (Zadnikova et al., 2010) |
| PIN4::PIN4:GFP | (Bennett et al., 2016; Belteton et al., 2018) |
| PIN7::PIN7:GFP | (Belteton et al., 2018) |
| *pin1-1* | ABRC; WT at the TTG1 (AT5G24520) locus (Goto N, 1987; Galweiler et al., 1998; Sawchuk et al., 2013) |
| *pin1-134* | Derived from *Atpin1::En134* (Galweiler et al., 1998); contains a 4-bp (AATT) insertion between +134 and +135 of *PIN1* (AT1G73590), resulting in a stop codon after amino acid 62. |
| *pin3-3* | (Friml et al., 2002b) |
| *pin4-2* | (Friml et al., 2002a) |
| *pin7^En^* | (Blilou et al., 2005) |
| *eir1-1* (*pin2*) | ABRC; (Roman et al., 1995; Luschnig et al., 1998) |
| *toz-1* | (Griffith et al., 2007) |
| *mp^G12^* | (Hardtke and Berleth, 1998) |
| *pin6* | ABRC; (Sawchuk et al., 2013) |
| *pin8-1* | ABRC; (Bosco et al., 2012) |
| ABCB1::ABCB1:GFP | (Dhonukshe et al., 2008; Mravec et al., 2008) |
| ABCB19::ABCB19:GFP | (Dhonukshe et al., 2008; Mravec et al., 2008) |
| *pgp1-100* (*abcb1*) | ABRC; (Lin and Wang, 2005) |
| *mdr1-101* (*abcb19*) | ABRC; (Lin and Wang, 2005) |
| *ucu2-4* (*twd1*) | ABRC; (Perez-Perez et al., 2004) |
| *aux1-21*;*lax1*;*2-1*;*3* | (Bainbridge et al., 2008) |
| *aux1-355* | ABRC; SALK_020355 (Alonso et al., 2003); contains a T-DNA insertion after +631 of *AUX1* (AT2G38120) |
| *lax1-064* | ABRC; SALK_071064 (Alonso et al., 2003); contains a T-DNA insertion after +814 of *LAX1* (AT5G01240) |
| *axr1-3* | ABRC; (Lincoln et al., 1990) |
| *axr1-12* | ABRC; (Lincoln et al., 1990) |
| *axl* | ABRC; SAIL_673_C11 (Sessions et al., 2002); contains a T-DNA insertion after +1390 of *AXL* (AT2G32410) |
| *tir1-1*;*afb2-3* | (Savaldi-Goldstein et al., 2008) |
| DR5rev::nYFP | (Heisler et al., 2005; Sawchuk et al., 2013) |

All gene coordinates are relative to the adenine (position +1) of the start codon.

**Alonso, J. M., Stepanova, A. N., Leisse, T. J., Kim, C. J., Chen, H., Shinn, P., Stevenson, D. K., Zimmerman, J., Barajas, P., Cheuk, R. et al.** (2003). Genome-wide insertional mutagenesis of Arabidopsis thaliana. *Science* **301**, 653-657.

**Bainbridge, K., Guyomarc’h, S., Bayer, E., Swarup, R., Bennett, M., Mandel, T. and Kuhlemeier, C.** (2008). Auxin influx carriers stabilize phyllotactic patterning. *Genes Dev.* **22**, 810-823.

**Belteton, S. A., Sawchuk, M. G., Donohoe, B. S., Scarpella, E. and Szymanski, D. B.** (2018). Reassessing the Roles of PIN Proteins and Anticlinal Microtubules during Pavement Cell Morphogenesis. *Plant Physiol* **176**, 432-449.

**Benkova, E., Michniewicz, M., Sauer, M., Teichmann, T., Seifertova, D., Jurgens, G. and Friml, J.** (2003). Local, efflux-dependent auxin gradients as a common module for plant organ formation. *Cell* **115**, 591-602.

**Bennett, T., Hines, G., van Rongen, M., Waldie, T., Sawchuk, M. G., Scarpella, E., Ljung, K. and Leyser, O.** (2016). Connective Auxin Transport in the Shoot Facilitates Communication between Shoot Apices. *PLoS Biol.* **14**, e1002446.

**Blilou, I., Xu, J., Wildwater, M., Willemsen, V., Paponov, I., Friml, J., Heidstra, R., Aida, M., Palme, K. and Scheres, B.** (2005). The PIN auxin efflux facilitator network controls growth and patterning in Arabidopsis roots. *Nature* **433**, 39-44.

**Bosco, C. D., Dovzhenko, A., Liu, X., Woerner, N., Rensch, T., Eismann, M., Eimer, S., Hegermann, J., Paponov, I. A., Ruperti, B. et al.** (2012). The endoplasmic reticulum localized PIN8 is a pollen specific auxin carrier involved in intracellular auxin homeostasis. *Plant J* **71**, 860-870.

**Dhonukshe, P., Grigoriev, I., Fischer, R., Tominaga, M., Robinson, D. G., Hasek, J., Paciorek, T., Petrasek, J., Seifertova, D., Tejos, R. et al.** (2008). Auxin transport inhibitors impair vesicle motility and actin cytoskeleton dynamics in diverse eukaryotes. *Proc. Natl. Acad. Sci. U S A* **105**, 4489-4494.

**Friml, J., Benkova, E., Blilou, I., Wisniewska, J., Hamann, T., Ljung, K., Woody, S., Sandberg, G., Scheres, B., Jurgens, G. et al.** (2002a). AtPIN4 mediates sink-driven auxin gradients and root patterning in Arabidopsis. *Cell* **108**, 661-673.

**Friml, J., Wisniewska, J., Benkova, E., Mendgen, K. and Palme, K.** (2002b). Lateral relocation of auxin efflux regulator PIN3 mediates tropism in Arabidopsis. *Nature* **415**, 806-809.

**Galweiler, L., Guan, C., Muller, A., Wisman, E., Mendgen, K., Yephremov, A. and Palme, K.** (1998). Regulation of polar auxin transport by AtPIN1 in Arabidopsis vascular tissue. *Science* **282**, 2226-2230.

**Geldner, N., Richter, S., Vieten, A., Marquardt, S., Torres-Ruiz, R. A., Mayer, U. and Jurgens, G.** (2004). Partial loss-of-function alleles reveal a role for GNOM in auxin transport-related, post-embryonic development of Arabidopsis. *Development* **131**, 389-400.

**Goto N, S. M., Kranz AR**. (1987). Effect of gibberllins on flower development of the pin-formed mutant of Arabidopsis thaliana. *Arabidopsis Information Service* **23**, 66-71.

**Griffith, M. E., Mayer, U., Capron, A., Ngo, Q. A., Surendrarao, A., McClinton, R., Jürgens, G. and Sundaresan, V.** (2007). The TORMOZ gene encodes a nucleolar protein required for regulated division planes and embryo development in Arabidopsis. *Plant Cell* **19**, 2246-2263.

**Hardtke, C. S. and Berleth, T.** (1998). The Arabidopsis gene MONOPTEROS encodes a transcription factor mediating embryo axis formation and vascular development. *EMBO J.* **17**, 1405-1411.

**Heisler, M. G., Ohno, C., Das, P., Sieber, P., Reddy, G. V., Long, J. A. and Meyerowitz, E. M.** (2005). Patterns of Auxin Transport and Gene Expression during Primordium Development Revealed by Live Imaging of the Arabidopsis Inflorescence Meristem. *Curr. Biol.* **15**, 1899-1911.

**Koizumi, K., Sugiyama, M. and Fukuda, H.** (2000). A series of novel mutants of Arabidopsis thaliana that are defective in the formation of continuous vascular network: calling the auxin signal flow canalization hypothesis into question. *Development* **127**, 3197-3204.

**Lin, R. and Wang, H.** (2005). Two homologous ATP-binding cassette transporter proteins, AtMDR1 and AtPGP1, regulate Arabidopsis photomorphogenesis and root development by mediating polar auxin transport. *Plant Physiol* **138**, 949-964.

**Lincoln, C., Britton, J. H. and Estelle, M.** (1990). Growth and development of the axr1 mutants of Arabidopsis. *Plant Cell* **2**, 1071-1080.

**Luschnig, C., Gaxiola, R. A., Grisafi, P. and Fink, G. R.** (1998). EIR1, a root-specific protein involved in auxin transport, is required for gravitropism in Arabidopsis thaliana. *Genes & Development* **12**, 2175-2187.

**Mravec, J., Kubes, M., Bielach, A., Gaykova, V., Petrasek, J., Skupa, P., Chand, S., Benkova, E., Zazimalova, E. and Friml, J.** (2008). Interaction of PIN and PGP transport mechanisms in auxin distribution-dependent development. *Development* **135**, 3345-3354.

**Okumura, K., Goh, T., Toyokura, K., Kasahara, H., Takebayashi, Y., Mimura, T., Kamiya, Y. and Fukaki, H.** (2013). GNOM/FEWER ROOTS is required for the establishment of an auxin response maximum for arabidopsis lateral root initiation. *Plant Cell Physiol* **54**, 406-417.

**Perez-Perez, J. M., Ponce, M. R. and Micol, J. L.** (2004). The ULTRACURVATA2 gene of Arabidopsis encodes an FK506-binding protein involved in auxin and brassinosteroid signaling. *Plant Physiol* **134**, 101-117.

**Roman, G., Lubarsky, B., Kieber, J. J., Rothenberg, M. and Ecker, J. R.** (1995). Genetic analysis of ethylene signal transduction in Arabidopsis thaliana: five novel mutant loci integrated into a stress response pathway. *Genetics* **139**, 1393-1409.

**Savaldi-Goldstein, S., Baiga, T. J., Pojer, F., Dabi, T., Butterfield, C., Parry, G., Santner, A., Dharmasiri, N., Tao, Y., Estelle, M. et al.** (2008). New auxin analogs with growth-promoting effects in intact plants reveal a chemical strategy to improve hormone delivery. *Proc. Natl. Acad. Sci. U S A* **105**, 15190-15195.

**Sawchuk, M. G., Edgar, A. and Scarpella, E.** (2013). Patterning of leaf vein networks by convergent auxin transport pathways. *PLoS Genet.* **9**, e1003294.

**Xu, J., Hofhuis, H., Heidstra, R., Sauer, M., Friml, J. and Scheres, B.** (2006). A molecular framework for plant regeneration. *Science* **311**, 385-388.

**Xu, J. and Scheres, B.** (2005). Dissection of Arabidopsis ADP-RIBOSYLATION FACTOR 1 function in epidermal cell polarity. *Plant Cell* **17**, 525-536.

**Zadnikova, P., Petrasek, J., Marhavy, P., Raz, V., Vandenbussche, F., Ding, Z., Schwarzerova, K., Morita, M. T., Tasaka, M., Hejatko, J. et al.** (2010). Role of PIN-mediated auxin efflux in apical hook development of Arabidopsis thaliana. *Development* **137**, 607-617.
