## Supplemental Table 2 for "Coordination of Tissue Cell Polarity by Auxin Transport and Signaling"

**Table S2. Embryo viability of WT, *pin3*;*4*;*7* and *pin2*;*3*;*4*;*7***

| **Genotype of self-fertilized parent** | **Proportion of viable embryos in siliques of self-fertilized parent (no. of non-aborted seeds / total no. of seeds)** | **Percentage of viable seeds in siliques of self-fertilized parent** |
| --- | --- | --- |
| WT | 293/293 | 100.0 |
| *pin3*/*pin3*;*pin4*/*pin4*;*pin7*/*pin7* | 275/276 | 99.6 |
| *pin2*/*pin2*;*pin3*/*pin3*;*pin4*/*pin4*;*pin7*/*pin7* | 271/271 | 100.0 |

Difference between *pin3*;*4*;*7* and WT and between *pin2*;*3*;*4*;*7* and WT was not significant by Kruskal-Wallis and Mann-Whitney test with Bonferroni correction.
