## Supplemental Table 3 for "Coordination of Tissue Cell Polarity by Auxin Transport and Signaling"

**Table S3. Embryo viability of *toz*, *mp*, *pin1*, *pin1*,*3*;*4*;*7*, *pin1*,*3*;*****2*;*4*;*7* and *pin1*,*3*,*6*;*4*;*7*;*8***

| **Genotype of self-fertilized parent** | **Proportion of viable embryos in siliques of self-fertilized parent (no. of non-aborted seeds / total no. of seeds)** | **Percentage of viable seeds in siliques of self-fertilized parent** |
| --- | --- | --- |
| *TOZ*/*toz-1* | 202/278 | 72.7 |
| *MP*/*mp^G12^* | 264/265^***^ | 99.6 |
| *PIN1*/*pin1-1* | 254/260^***^ | 97.7 |
| *PIN1*/*pin1-134* | 257/258^***^ | 99.6 |
| *PIN1*/*pin1-1*,*pin3*/*pin3*;*pin4*/*pin4*;*pin7*/*pin7* | 269/272^***^ | 98.9 |
| *PIN1*/*pin1-134*,*pin3*/*pin3*;*pin4*/*pin4*;*pin7*/*pin7* | 280/281^***^ | 99.6 |
| *PIN1*/*pin1-1*,*pin3*/*pin3*;*pin2*/*pin2*;*pin4*/*pin4*;*pin7*/*pin7* | 276/278^***^ | 99.3 |
| *PIN1*/*pin1-1*,*pin3*/*pin3*,*pin6*/*pin6*;*pin4*/*pin4*;*pin7*/*pin7*;*pin8*/*pin8* | 266/268^***^ | 99.2 |

Difference between negative control for completely penetrant embryo lethality (*mp^G12^*) and positive control for completely penetrant embryo lethality (*toz-1*), between *pin1-1* and *toz-1*, between *pin1-134* and *toz-1*, between *pin1-1*,*3*;*4*;*7* and *toz-1*, between *pin1-134*,*3*;*4*;*7* and *toz-1*, between *pin1-1*,*3*;*2*;*4*;*7* and *toz-1*, and between *pin1-1*,*3*,*6*;*4*;*7*;*8* and *toz-1* was significant at *P*<0.001 (***) by Kruskal-Wallis and Mann-Whitney test with Bonferroni correction. Difference between *pin1-1* and *mp^G12^*, between *pin1-134* and *mp^G12^*, between *pin1-1*,*3*;*4*;*7* and *mp^G12^*, between *pin1-134*,*3*;*4*;*7* and *mp^G12^*, between *pin1-1*,*3*;*2*;*4*;*7* and *mp^G12^*, and between *pin1-1*,*3*,*6*;*4*;*7*;*8* and *mp^G12^* was not significant by Kruskal-Wallis and Mann-Whitney test with Bonferroni correction.
