## Supplemental Table 4 for "Coordination of Tissue Cell Polarity by Auxin Transport and Signaling"

**Table S4. Embryo viability of *pin1*, *pin1*,*3*;*4*;*7*, *pin1*,*3*;*2*;*4*;*7* and *pin1*,*3*,*6*;*4*;*7*;*8***

| **Genotype of self-fertilized parent** | **Proportion of embryo-viable mutants in progeny of self-fertilized parent (no. of mutant seedlings / total no. of seedlings)** | **Percentage of embryo-viable mutants in progeny of self-fertilized parent** |
| --- | --- | --- |
| *PIN1*/*pin1-1* | 66/239 | 27.6 |
| *PIN1*/*pin1-134* | 53/227 | 23.3 |
| *PIN1*/*pin1-1*,*pin3*/*pin3*;*pin4*/*pin4*;*pin7*/*pin7* | 52/196 | 26.5 |
| *PIN1*/*pin1-1-134*,*pin3*/*pin3*;*pin4*/*pin4*;*pin7*/*pin7* | 56/228 | 24.6 |
| *PIN1*/*pin1-1*,*pin3*/*pin3*;*pin2*/*pin2*;*pin4*/*pin4*;*pin7*/*pin7* | 61/263 | 23.2 |
| *PIN1*/*pin1-1*,*pin3*/*pin3*,*pin6*/*pin6*;*pin4*/*pin4*;*pin7*/*pin7*;*pin8*/*pin8* | 65/260 | 25.0 |

Difference between observed and theoretical frequency distributions of embryo-viable mutants in the progeny of self-fertilized heterozygous parents was not significant by Pearson’s chi-squared (χ^2^) goodness-of-fit test (α=0.05, dF=1).
