## Supplemental Table 5 for "Coordination of Tissue Cell Polarity by Auxin Transport and Signaling"

**Table S5. Embryo viability of WT, *abcb1*, *abcb19*, *abcb1*;*19* and *twd1***

| **Genotype of self-fertilized parent** | **Proportion of viable embryos in siliques of self-fertilized parent (no. of non-aborted seeds / total no. of seeds)** | **Percentage of viable seeds in siliques of self-fertilized parent** |
| --- | --- | --- |
| WT | 294/294 | 100 |
| *abcb1*/*abcb1* | 269/272 | 98.9 |
| *abcb19*/*abcb19* | 271/276 | 98.2 |
| *abcb1*/*abcb1*;*abcb19*/*abcb19* | 276/332^***^ | 83.1 |
| *twd1*/*twd1* | 245/265^***^ | 92.4 |

Difference between *abcb1*;*19* and WT, and between *twd1* and WT was significant at *P*<0.001 (***). and between *abcb1* and WT, and between *abcb19* and WT was not significant by Kruskal-Wallis and Mann-Whitney test with Bonferroni correction.
