## Supplemental Table 6 for "Coordination of Tissue Cell Polarity by Auxin Transport and Signaling"

**Table S6. Embryo viability of *toz*, *mp*, *pin1*,*3*,*6* and *pin1*,*3*,*6*;*abcb1*;*19***

| **Genotype of self-fertilized parent** | **Proportion of viable embryos in siliques of self-fertilized parent (no. of non-aborted seeds / total no. of seeds)** | **Percentage of viable seeds in siliques of self-fertilized parent** |
| --- | --- | --- |
| *TOZ*/*toz-1* | 202/277 | 72.9 |
| *MP*/*mp^G12^* | 255/256^***^ | 99.6 |
| *PIN1*/*pin1-1*,*pin3*/*pin3*,*PIN6*/*pin6* | 263/266^***^ | 98.9 |
| *PIN1*/*pin1-1*,*PIN3*/*pin3*,*PIN6*/*pin6*;*abcb1*/*abcb1*;*abcb19*/*abcb19* | 240/284^*/***^ | 84.5 |

Difference between negative control for completely penetrant embryo lethality (*mp^G12^*) and positive control for completely penetrant embryo lethality (*toz-1*) and between *pin1-1*,*3*,*6* and *toz-1* was significant at *P*<0.001 (***), and between *pin1-1*,*3*,*6*;*abcb1*;*19* and *toz-1* was significant at *P*<0.05 (*) by Kruskal-Wallis and Mann-Whitney test with Bonferroni correction. Difference between *pin1-1*,*3*,*6*;*abcb1*;*19* and *mp^G12^* was significant at *P*<0.001 (***), and between *pin1-1*,*3*,*6* and *mp^G12^* was not significant by Kruskal-Wallis and Mann-Whitney test with Bonferroni correction. Linkage in *cis* between *pin1-1* and *pin6* in *PIN1*/*pin1-1*,*pin3*/*pin3*,*PIN6*/*pin6* was confirmed by phenotyping the progeny of the self-fertilized *PIN1*/*pin1-1*,*pin3*/*pin3*,*PIN6*/*pin6* plants used for the embryo viability analysis for the presence of seedlings with cup-shaped cotyledons, which are characteristic of *pin1*,*6* double homozygous mutant (Sawchuk et al., 2013). Linkage in *cis* between *pin1-1*, *pin3* and *pin6* in *PIN1*/*pin1-1*,*PIN3*/*pin3*,*PIN6*/*pin6*;*abcb1*/*abcb1*;*abcb19*/*abcb19* was confirmed by phenotyping the progeny of the self-fertilized *PIN1*/*pin1-1*,*PIN3*/*pin3*,*PIN6*/*pin6*;*abcb1*/*abcb1*;*abcb19*/*abcb19* plants used for the embryo viability analysis for the presence of seedlings with cup-shaped cotyledons and by genotyping those cup-shaped-cotyledon seedling for the *pin3* mutation.

**Sawchuk, M. G., Edgar, A. and Scarpella, E.** (2013). Patterning of leaf vein networks by convergent auxin transport pathways. *PLoS Genet.* **9**, e1003294.
