## Supplemental Table 7 for "Coordination of Tissue Cell Polarity by Auxin Transport and Signaling"

**Table S7. Embryo viability of *pin1*,*3*,*6* and *pin1*,*3*,*6*;*abcb1*;*19***

| **Genotype of self-fertilized parent** | **Proportion of embryo-viable mutants in progeny of self-fertilized parent (no. of mutant seedlings / total no. of seedlings)** | **Percentage of embryo-viable mutants in progeny of self-fertilized parent** |
| --- | --- | --- |
| *PIN1*/*pin1-1*,*pin3*/*pin3*,*PIN6*/*pin6* | 80/361 | 22.2 |
| *PIN1*/*pin1-1*,*PIN3*/*pin3*,*PIN6*/*pin6*;*abcb1*/*abcb1*;*abcb19*/*abcb19* | 74/335 | 22.1 |

Difference between observed and theoretical frequency distributions of embryo-viable mutants in the progeny of self-fertilized heterozygous parents was not significant by Pearson’s chi-squared (χ2) goodness-of-fit test (α=0.05, dF=1). Genotype of the mutants seedlings of *PIN1*/*pin1-1*,*pin3*/*pin3*,*PIN6*/*pin6* was confirmed by genotyping all mutant seedlings for *pin1-1* and *pin6* mutation. Genotype of the mutants seedlings of *PIN1*/*pin1-1*,*PIN3*/*pin3*,*PIN6*/*pin6*;*abcb1*/*abcb1*;*abcb19*/*abcb19* was confirmed by genotyping all mutant seedlings for *pin1-1*, *pin3* and *pin6* mutation.
