## Supplemental Table 8 for "Coordination of Tissue Cell Polarity by Auxin Transport and Signaling"

**Table S8. Embryo viability of WT, *aux1*, *lax1*, *aux1*;*lax1* and *aux1*;*lax1*;*2*;*3***

| **Genotype of self-fertilized parent** | **Proportion of viable embryos in siliques of self-fertilized parent (no. of non-aborted seeds / total no. of seeds)** | **Percentage of viable seeds in siliques of self-fertilized parent** |
| --- | --- | --- |
| WT | 272/274 | 99.3 |
| *aux1*/*aux1-355* | 266/267 | 99.6 |
| *lax1*/*lax1-064* | 265/267 | 99.2 |
| *aux1*/*aux1-355*;*lax1*/*lax1-064* | 278/281 | 98.9 |
| *aux1*/*aux1-21*;*lax1*/*lax1*;*lax2*/*lax2-1*;*lax3*/*lax3* | 261/262 | 99.6 |

Difference between *aux1-355* and WT, between *lax1-064* and WT, between *aux1-355*;*lax1-064* and WT, and between *aux1-21*;*lax1*;*2-1*;*3* and WT was not significant by Kruskal-Wallis and Mann-Whitney test with Bonferroni correction.
