## Supplemental Table 9 for "Coordination of Tissue Cell Polarity by Auxin Transport and Signaling"

**Table S9. Embryo viability of *toz*, *mp*, *pin1*,*3*,*6* and *pin1*,*3*,*6*;*aux1*;*lax1***

| **Genotype of self-fertilized parent** | **Proportion of viable embryos in siliques of self-fertilized parent (no. of non-aborted seeds / total no. of seeds)** | **Percentage of viable seeds in siliques of self-fertilized parent** |
| --- | --- | --- |
| *TOZ*/*toz*-*1* | 185/244^***^ | 75.8 |
| *MP*/*mp^G12^* | 220/220 | 100 |
| *PIN1*/*pin1-1*,*pin3*/*pin3*,*PIN6*/*pin6* | 259/261^***^ | 99.2 |
| *PIN1*/*pin1-1*,*pin3*/*pin3*,*PIN6*/*pin6*;*aux1*/*aux1-355*;*lax1*/*lax1-064* | 280/282^***^ | 99.3 |

Difference between negative control for completely penetrant embryo lethality (*mp^G12^*) and positive control for completely penetrant embryo lethality (*toz-1*), between *pin1*-*1*,*3*,*6* and *toz*-*1*, and between *pin1-1*,*3*,*6*;*aux1-355*;*lax1-064* and *toz-1* was significant at *P*<0.001 (***) by Kruskal-Wallis and Mann-Whitney test with Bonferroni correction. Difference between *pin1-1*,*3*,*6* and *mp^G12^*, and between *pin1-1*,*3*,*6*;*aux1-355*;*lax1-064* and *mp^G12^* was not significant by Kruskal-Wallis and Mann-Whitney test with Bonferroni correction. Linkage in *cis* between *pin1-1* and *pin6* in *PIN1*/*pin1-1*,*pin3*/*pin3*,*PIN6*/*pin6* and *PIN1*/*pin1-1*,*pin3*/*pin3*,*PIN6*/*pin6*;*aux1*/*aux1-355*;*lax1*/*lax1-064* was confirmed by phenotyping the progeny of the self-fertilized plants used for the embryo viability analysis for the presence of seedlings with cup-shaped cotyledons, which are characteristic of *pin1*,*6* double homozygous mutant (Sawchuk et al., 2013).

**Sawchuk, M. G., Edgar, A. and Scarpella, E.** (2013). Patterning of leaf vein networks by convergent auxin transport pathways. *PLoS Genet.* **9**, e1003294.
