## Supplemental Table 10 for "Coordination of Tissue Cell Polarity by Auxin Transport and Signaling"

**Table S10. Embryo viability of *pin1*,*3*,*6* and *pin1*,*3*,*6*;*aux1*;*lax1***

| **Genotype of self-fertilized parent** | **Proportion of embryo-viable mutants in progeny of self-fertilized parent (no. of mutant seedlings / total no. of seedlings)** | **Percentage of embryo-viable mutants in progeny of self-fertilized parent** |
| --- | --- | --- |
| *PIN1*/*pin1-1*,*pin3*/*pin3*,*PIN6*/*pin6* | 87/390 | 22.3 |
| *PIN1*/*pin1-1*,*pin3*/*pin3*,*PIN6*/*pin6*;*aux1*/*aux1-355*;*lax1*/*lax1-064* | 109/489 | 22.3 |

Difference between observed and theoretical frequency distributions of embryo-viable mutants in the progeny of self-fertilized heterozygous parents was not significant by Pearson’s chi-squared (χ2) goodness-of-fit test (α=0.05, dF=1). Genotype of the mutants seedlings of both *PIN1*/*pin1-1*,*pin3*/*pin3*,*PIN6*/*pin6* and *PIN1*/*pin1-1*,*pin3*/*pin3*,*PIN6*/*pin6*;*aux1*/*aux1-355*;*lax1*/*lax1-064* was confirmed by genotyping all mutant seedlings for *pin1-1* and *pin6* mutation.
