## Supplemental Table 11 for "Coordination of Tissue Cell Polarity by Auxin Transport and Signaling"

**Table S11. Embryo viability of *axr1*;*axl*, *tir1*;*afb2*, *gn*;*pin1*,*3*;*4*;*7* and *gn*;*pin1*,*3*,*6*;*4*;*7*;*8***

| **Genotype of self-fertilized parent** | **Proportion of viable embryos in siliques of self-fertilized parent**  **(no. of non-aborted seeds / total no. of seeds)** | **Percentage of viable seeds in siliques of self-fertilized parent** |
| --- | --- | --- |
| *AXR1/axr1-12*;*AXL/axl* | 900/978 | 92 |
| *TIR1/tir1*;*AFB2/afb2* | 777/781^***^ | 99.5 |
| *GN/gn-13*;*PIN1/pin1-1*,*pin3/pin3*;*pin4/pin4*;*pin7/pin7* | 482/484^***^ | 99.6 |
| *GN/gn-13*;*PIN1/pin1-1*,*pin3/pin3*,*pin6/pin6*;*pin4/pin4*;*pin7/pin7*;*pin8/pin8* | 571/575^***^ | 99.3 |

Difference between negative control for completely penetrant embryo lethality (*tir1*;*afb2*) and positive control for completely penetrant embryo lethality (*axr1-12*;*axl*), between *gn*;*pin1-1*,*3*;*4*;*7* and *axr1-12*;*axl*, and between *gn*;*pin1-1*,*3*,*6*;*4*;*7*;*8* and *axr1-12*;*axl* was significant at *P*<0.001 (***) by Kruskal-Wallis and Mann-Whitney test with Bonferroni correction. Difference between *gn*;*pin1-1*,*3*;*4*;*7* and *tir1*;*afb2*, and between *gn*;*pin1-1*,*3*,*6*;*4*;*7*;*8* and *tir1*;*afb2* was not significant by Kruskal-Wallis and Mann-Whitney test with Bonferroni correction.
