## Supplemental Table 12 for "Coordination of Tissue Cell Polarity by Auxin Transport and Signaling"

**Table S12. Embryo viability of *gn*;*pin1*,*3*;*4*;*7* and *gn*;*pin1*,*3*,*6*;*4*;*7*;*8***

| **Genotype of self-fertilized parent** | **Proportion of embryo-viable mutants in progeny of self-fertilized parent (no. of mutant seedlings / total no. of seedlings)** | **Percentage of embryo-viable mutants in progeny of self-fertilized parent** |
| --- | --- | --- |
| *GN/gn-13*;*PIN1*/*pin1-1*,*pin3*/*pin3*;*pin4*/*pin4*;*pin7*/*pin7* | 256/3624 | 7.1 |
| *GN/gn-13*;*PIN1*/*pin1-1*,*pin3*/*pin3,pin6*/*pin6;pin4*/*pin4*;*pin7*/*pin7*;*pin8*/*pin8* | 222/3231 | 6.9 |

Difference between observed and theoretical frequency distributions of embryo-viable mutants in the progeny of self-fertilized heterozygous parents was not significant by Pearson’s chi-squared (χ^2^) goodness-of-fit test (α=0.05, dF=1).
