## Supplemental Table 13 for "Coordination of Tissue Cell Polarity by Auxin Transport and Signaling"

**Table S13. Embryo viability of WT, *axr1* and *tir1*;*afb2***

| **Genotype of self-fertilized parent** | **Proportion of viable embryos in siliques of self-fertilized parent (no. of non-aborted seeds / total no. of seeds)** | **Percentage of viable seeds in siliques of self-fertilized parent** |
| --- | --- | --- |
| WT | 408/412 | 99 |
| *axr1-3* | 391/403 | 97 |
| *tir1*;*afb2* | 300/303 | 99 |

Difference between *axr1-3* and WT, and between *tir1*;*afb2* and WT was not significant by Kruskal-Wallis and Mann-Whitney test with Bonferroni correction.
