## Supplemental Table 14 for "Coordination of Tissue Cell Polarity by Auxin Transport and Signaling"

**Table S14. Embryo viability of *toz*, *mp*, *pin1*,*3*,*6*;*4*;7;*8*, *pin1*,*3*,*6*;*4*;*7*;*8*;*axr1*, *pin1*,*3*,*6*;*4*;*7*;*8*;*tir1*;*afb2***

| **Genotype of self-fertilized parent** | **Proportion of viable embryos in siliques of self-fertilized parent (no. of non-aborted seeds / total no. of seeds)** | **Percentage of viable seeds in siliques of self-fertilized parent** |
| --- | --- | --- |
| *TOZ*/*toz*-*1* | 190/239 | 79.5 |
| *MP*/*mp^G12^* | 261/262^***^ | 99.6 |
| *PIN1/pin1-1*,*pin3/pin3*,*pin6/pin6*;*pin4/pin4*;*pin7/pin7*;*pin8/pin8* | 243/244^***^ | 99.6 |
| *PIN1/pin1-1*,*pin3/pin3*,*pin6/pin6*;*pin4/pin4*;*pin7/pin7*;*pin8/pin8*;*axr1*/*axr1-3* | 240/248^***^ | 96.8 |
| *PIN1/pin1-1*,*pin3/pin3*,*pin6/pin6*;*pin4/pin4*;*pin7/pin7*;*pin8/pin8*;*tir1*/*tir1*;*afb2*/*afb2* | 473/475^***^ | 99.6 |

Difference between negative control for completely penetrant embryo lethality (*mp^G12^*) and positive control for completely penetrant embryo lethality (*toz-1*), between *pin1-1*,*3*,*6*;*4*;*7*;*8* and *toz-1*, between *pin1-1*,*3*,*6*;*4*;*7*;*8*;*axr1-3* and *toz-1*, and between *pin1-1*,*3*,*6*;*4*;*7*;*8*;*tir1*;*afb2* and *toz-1* was significant at *P*<0.001 (***) by Kruskal-Wallis and Mann-Whitney test with Bonferroni correction. Difference between *pin1-1*,*3*,*6*;*4*;*7*;*8* and *mp^G12^*, between *pin1-1*,*3*,*6*;*4*;*7*;*8*;*axr1-3* and *mp^G12^*, and between *pin1-1*,*3*,*6*;*4*;*7*;*8*;*tir1*;*afb2* and *mp^G12^* was not significant by Kruskal-Wallis and Mann-Whitney test with Bonferroni correction.
