## Supplemental Table 15 for "Coordination of Tissue Cell Polarity by Auxin Transport and Signaling"

**Table S15. Embryo viability of *pin1*,*3*,*6*;*4*;*7*;*8*;*axr1* and *pin1*,*3*,*6*;*4*;*7*;*8*;*tir1*;*afb2***

| **Genotype of self-fertilized parent** | **Proportion of embryo-viable mutants in progeny of self-fertilized parent (no. of mutant seedlings / total no. of seedlings)** | **Percentage of embryo-viable mutants in**  **progeny of self-fertilized parent** |
| --- | --- | --- |
| *PIN1/pin1-1*,*pin3/pin3*,*pin6/pin6*;*pin4/pin4*;*pin7/pin7*;*pin8/pin8*;*axr1*/*axr1-3* | 66/277 | 23.8 |
| *PIN1/pin1-1*,*pin3/pin3*,*pin6/pin6*;*pin4/pin4*;*pin7/pin7*;*pin8/pin8*;*tir1/tir1*;*afb2/afb2* | 77/324 | 23.8 |

Difference between observed and theoretical frequency distributions of embryo-viable mutants in the progeny of self-fertilized heterozygous parents was not significant by Pearson’s chi-squared (χ^2^) goodness-of-fit test (α=0.05, dF=1).
