## Supplemental Table 16 for "Coordination of Tissue Cell Polarity by Auxin Transport and Signaling"

**Table S16. Embryo viability of *toz*, *mp*, *gn* and *gn*;*axr1***

| **Genotype of self-fertilized parent** | **Proportion of viable embryos in siliques of self-fertilized parent (no. of non-aborted seeds / total no. of seeds)** | **Percentage of viable seeds in siliques of self-fertilized parent** |
| --- | --- | --- |
| *TOZ*/*toz*-*1* | 206/259 | 79.5 |
| *MP*/*mp^G12^* | 243/247^***^ | 98.4 |
| *GN*/*gn-13* | 248/252^***^ | 98.4 |
| *GN*/*gn-13*;*axr1*/*axr1-3* | 264/270^***^ | 97.8 |
| *GN*/*gn-13*;*axr1*/*axr1-12* | 214/224^***^ | 95.6 |

Difference between negative control for completely penetrant embryo lethality (*mp^G12^*) and positive control for completely penetrant embryo lethality (*toz-1*), between *gn-13* and *toz-1*, between *gn-13*;*axr1-3* and *toz-1*, and between *gn-13*;*axr1-12* and *toz-1* was significant at *P*<0.001 (***) by Kruskal-Wallis and Mann-Whitney test with Bonferroni correction. Difference between *gn-13* and *mp^G12^*, between *gn-13*;*axr1-3* and *mp^G12^*, and between *gn-13*;*axr1-12* and *mp^G12^* was not significant by Kruskal-Wallis and Mann-Whitney test with Bonferroni correction.
