## Supplemental Table 17 for "Coordination of Tissue Cell Polarity by Auxin Transport and Signaling"

**Table S17. Embryo viability of *gn* and *gn*;*axr1***

| **Genotype of self-fertilized parent** | **Proportion of embryo-viable mutants in progeny of self-fertilized parent (no. of mutant seedlings / total no. of seedlings)** | **Percentage of embryo-viable mutants in progeny of self-fertilized parent** |
| --- | --- | --- |
| *GN*/*gn-13* | 101/411 | 24.6 |
| *GN*/*gn1-13*;*axr1-3* | 74/321 | 23.0 |
| *GN*/*gn1-13*;*axr1-12* | 70/276 | 25.4 |

Difference between observed and theoretical frequency distributions of embryo-viable mutants in the progeny of self-fertilized heterozygous parents was not significant by Pearson’s chi-squared (χ^2^) goodness-of-fit test (α=0.05, dF=1).
