## Supplemental Table 18 for "Coordination of Tissue Cell Polarity by Auxin Transport and Signaling"

**Table S18. Genotyping strategies**

| **Line** | **Strategy** |
| --- | --- |
| *gn-13* | *GN*: ‘SALK_045424 gn LP’ and ‘SALK_045424 gn RP’; *gn*: ‘SALK_045424 gn RP’ and ‘LBb1.3’ |
| *gn-18* | *GN*: ‘Salk026031 LP gnp close’ and ‘Salk026031 RP gnp close’; *gn*: ‘Salk026031 RP gnp close’ and ‘LBb1.3’ |
| *fwr* (*gn^fwr^*) | ‘FWR for’ and ‘FWR REV2’; *EcoR*I |
| *van7*/*emb30-7* (*gn^van7^*) | ‘van7 Hpa1 FP’ and ‘van7 Hpa1 RP’; *Hpa*I |
| *pin1-1* | ‘pin1-1 F’ and ‘pin1-1 R’; *Tat*I |
| *pin1-134* | ‘pin1-1 F’ and ‘pin1-134 R mse-I’; *Mse*I |
| *pin3-3* | ‘pin3-3 F’ and ‘pin3-3 R’; *Sty*I |
| *pin4-2* | *PIN4*: ‘PIN4 forw geno II’ and ‘PIN4en rev Ikram’; *pin4*: ‘PIN4en rev Ikram’ and ‘en primer’ |
| *pin7^En^* | *PIN7*: ‘PIN7en forw Ikram’ and ‘PIN7en rev’; *pin7*: ‘PIN7en rev Ikram II’ and ‘en primer’ |
| *eir1-1* (*pin2*) | ‘eir1-1 F’ and ‘eir1-1 R’; *Bse*LI |
| *pin6* | *PIN6*: ‘PIN6 spm F’ and ‘PIN6 spm R’; *pin6*: ‘PIN6 spm F’ and ‘Spm32’ |
| *pin8-1* | *PIN8*: ‘SALK_107965 LP’ and ‘SALK_107965 RP’; *pin8*: ‘SALK_107965 RP’ and ‘LBb1.3’ |
| *pgp1-100* (*abcb1*) | *ABCB1*: ‘SALK_083649 pgp1-100 LP’ and ‘SALK_083649 pgp1-100 RP’; *abcb1*: ‘SALK_083649 pgp1-100 RP’ and ‘LBb1.3’ |
| *mdr1-101* (*abcb19*) | *ABCB19*: ‘SALK_033455 atmdr1-101 LP’ and ‘SALK_033455 atmdr1-101 RP’; *abcb19*: ‘SALK_033455 atmdr1-101 RP’ and ‘LBb1.3’ |
| *ucu2-4* (*twd1*) | *UCU2*: ‘SALK_012836 twd1 LP’ and ‘SALK_012836 twd1 RP’; *ucu2*: ‘SALK_012836 twd1 RP’ and ‘LBb1.3’ |
| *aux1-21* | ‘aux1-21 Fwd’ and ‘aux1-21 Rev’; *ApaL*I |
| *lax1* | *LAX1*: ‘lax1 Fwd’ and ‘lax1 WT Rev’; *lax1*: ‘lax1 fwd’ and ‘lax123 mutant Rev’ |
| *lax2-1* | *LAX2*: ‘lax2 Fwd’ and ‘lax2 WT Rev’; *lax2*: ‘lax2 fwd’ and ‘lax123 mutant Rev’ |
| *lax3* | *LAX3*: ‘lax3 Fwd’ and ‘lax3 WT Rev’; *lax3*: ‘lax3 fwd’ and ‘dSpm5’ |
| *aux1-355* | *AUX1*: ‘SALK_020355 LP (aux1)’ and ‘SALK_020355 RP (aux1)’; *aux1*: ‘SALK_020355 RP (aux1)’ and ‘LBb1.3’ |
| *lax1-064* | *LAX1*: ‘SALK_071064 lax1 LP’ and ‘SALK_071064 lax1 RP’; *lax1*: ‘SALK_071064 lax1 RP’ and ‘LBb1.3’ |
| *axr1-3* | ‘AXR1-Acc1’ and ‘AXR1-15’; *Sal*I |
| *axr1-12* | ‘axr1-12 forw’ and ‘axr1-12 rev’; *Dra*I |
| *axl* | *AXL*: ‘AXL SAIL LP’ and ‘AXL SAIL RP’; *axl*: ‘AXL SAIL RP’ and ‘LB3’ |
| *tir1-1* | ‘tir1-1F2’ and ‘tir1-1R2’, *Bsa*I |
| *afb2-3* | *AFB2*: ‘AFB2+F’ and ‘AFB2-TR’; *afb2*: ‘pROK-LB’ and ‘AFB2-TR’ |
