## Supplemental Table 19 for "Coordination of Tissue Cell Polarity by Auxin Transport and Signaling"

**Table S19. Oligonucleotide sequences**

| **Name** | **Sequence (5’ to 3’)** |
| --- | --- |
| SALK_045424 gn LP | TGATCCAAATCACTGGGTTTC |
| SALK_045424 gn RP | AGCTGAAGATAGGGAATTCGC |
| LBb1.3 | ATTTTGCCGATTTCGGAAC |
| Salk026031 LP gnp close | TGAAAGAGACATGTCCTTCGG |
| Salk026031 RP gnp close | GACACGTCTCGCTAAATCTCG |
| FWR for | AAGAGCCAAGATCACAGCCTACTG |
| FWR REV2 | GAGAGCACGCGCAAGCTGCAACAAG |
| van7 Hpa1 FP | ATCCGTGCCCTTGATCTAATGGGAG |
| van7 Hpa1 RP | CACTTTTCTTAGTCCTTGAACAAGCGTTAA |
| GN Fwd NotI | TCTGCGGCCGCTCTAGAGGTGTGTATGATAATG |
| GN Rev NotI | TTTGCGGCCGCTCTAGAAATCGAAATCCGTCTC |
| fwr-mutagenesis F | GCTTGCGCGTGCTCTCATTTGGGC |
| fwr-mutagenesis R | TGCAACAAAAATTCAGCTTGTAGAAACTTGCTTTCG |
| pin1-1 F | ATGATTACGGCGGCGGACTTCTA |
| pin1-1 R | TTCCGACCACCACCAGAAGCC |
| pin1-134 R mse-I | CTCAGCTTCAGTTTCCAAAGGTTG |
| pin3-3 F | GGAGCTCAAACGGGTCACCCG |
| pin3-3 R | GCTGGATGAGCTACAGCTATATTC |
| PIN4 forw geno II | GTCCGACTCCACGGCCTTC |
| PIN4en rev Ikram | ATCTTCTTCTTCACCTTCCACTCT |
| en primer | GAGCGTCGGTCCCCACACTTCTATAC |
| PIN7en forw Ikram | CCTAACGGTTTCCACACTCA |
| PIN7en rev | TAGCTCTTTAGGGTTTAGCTC |
| PIN7en rev Ikram II | GGTTTAGCTCTGCTGTGGAGTT |
| eir1-1 F | TTGTTGATCATTTTACCTGGGACA |
| eir1-1 R | GGTTGCAATGCCATAAATAGAC |
| PIN6 spm F | CATAACGAAGCTAACTAAGGGGTAATCTC |
| PIN6 spm R | GGAGTTCAAAGAGGAATAGTAGCAGAG |
| Spm32 | TACGAATAAGAGCGTCCATTTTAGAGTG |
| SALK_107965 LP | TGAAAGACATTTTGATGGCATC |
| SALK_107965 RP | CCAAATCAAGCTTTGCAAGAC |
| SALK_083649 pgp1-100 LP | GAAGACTGCGACAAGGACAAG |
| SALK_083649 pgp1-100 RP | GCAAGAGCGATGTTGAAGAAC |
| SALK_033455 atmdr1-101 LP | GCAATTGCAATTCTCTGCTTC |
| SALK_033455 atmdr1-101 RP | CTCAGGCAATTGCTCAAGTTC |
| SALK_012836 twd1 LP | GTGAAGCTGAGGTCTTGGATG |
| SALK_012836 twd1 RP | TATGGCCTGAAACAGCAAACC |
| aux1-21 Fwd | CTGGAAAGCACTAGGACTCGC |
| aux1-21 Rev | AAGCGGCGAAGAAACGATACAG |
| lax1 Fwd | ATATGGTTGCAGGTGGCACA |
| lax1 WT Rev | GTAACCGGCAAAAGCTGCA |
| lax123 mutant Rev | AAGCACGACGGCTGTAGAATAG |
| lax2 Fwd | ATGGAGAACGGTGAGAAAGCAGC |
| lax2 WT Rev | CGCAGAAGGCAGCGTTAGCG |
| lax3 Fwd | TACTTCACCGGAGCCACCA |
| lax3 WT Rev | TGATTGGTCCGAAAAAGG |
| dSpm5 | CGGGATCCGACACTCTTTAATTAACTGACACTC |
| SALK_020355 LP (aux1) | GGCTCCCGTAAAATAAAGCAC |
| SALK_020355 RP (aux1) | AATTATCGTTGGTTTCAGGTGG |
| SALK_071064 lax1 LP | CAATAGTAGTCTCCGGGGAGG |
| SALK_071064 lax1 RP | ACAACACAAGCTTGGTTGGAC |
| AXR1-Acc1 | AAACCAACTTAACGTTTGCATGTCG |
| AXR1-15 | TCTCATATGTACTTTTCCTCGTCCTCTTCAC |
| axr1-12 forw | CCGAGCAGCATCCCAAAAC |
| axr1-12 rev | GTTGGCAGCAAATCTGTCCG |
| AXL SAIL LP | TGGACTTACTGGGTTTGTTCG |
| AXL SAIL RP | CAAACCTTGAGTGCTGCTACC |
| LB3 | TAGCATCTGAATTTCATAACCAATCTCGATACAC |
| SALK_045424 gn LP | TGATCCAAATCACTGGGTTTC |
| SALK_045424 gn RP | AGCTGAAGATAGGGAATTCGC |
| tir1-1F2 | AGCGACGGTGATTAGGAGG |
| tir1-1R2 | CAGGAACAACGCAGCAAAA |
| AFB2+F | TTCTCCTTCGATCATTGTCAAC |
| AFB2-TR | TAGCGGCAATAGAGGCAAGA |
| pROK-LB | GGAACCACCATCAAACAGGA |
| GN_qFb | ACTTGTCAACAGAGCTGGTAGC |
| GN_qRb | GCTGCAAACCATCGAAAGAATC |
| ROC1 F | CAAACCTCTTCTTCAGTCTGATAGAGA |
| ROC1 R | GAGTGCTCATTCCTTATTTCTGGTAG |
