## Supplemental Table 20 for "Coordination of Tissue Cell Polarity by Auxin Transport and Signaling"

**Table S20. Light paths**

| **Fluorophore** | **Laser** | **Wavelength**  **(nm)** | **Main dichroic beam splitter** | **First secondary dichroic beam splitter** | **Second secondary dichroic beam splitter** | **Emission filter (detector)** |
| --- | --- | --- | --- | --- | --- | --- |
| YFP | Ar | 514 | HFT 405/514/594 | NFT 595 | NFT 515 | BP 520-555 IR (PMT3) |
| GFP;  Autofluorescence | Ar | 488 | HFT 405/488/594 | NFT 545 | NFT 490 (PMT3); Plate (META) | BP 505-530 (PMT3);  550-574 (META) |
| GFP | Ar | 488 | HFT 405/488/594 | NFT 545 | NFT 490 | BP 505-530 (PMT3) |
